# A fluorescent probe enables the discovery of improved antagonists targeting the intracellular allosteric site of the chemokine receptor CCR7

**DOI:** 10.1101/2024.08.27.607356

**Authors:** Silas L. Wurnig, Max E. Huber, Corinna Weiler, Hanna Baltrukevich, Nicole Merten, Isabel Stötzel, Yinshui Chang, René H. L. Klammer, Dirk Baumjohann, Eva Kiermaier, Peter Kolb, Evi Kostenis, Matthias Schiedel, Finn K. Hansen

## Abstract

Intracellularly acting ligands of G protein-coupled receptors (GPCRs) are gaining significant interest in GPCR drug discovery. In this study, we report the development of the fluorescent ligand Mz437 (**4**) targeting the CC chemokine receptor CCR7 at an intracellular allosteric site. We demonstrate its experimental power by applying **4** to identify two improved intracellular CCR7 antagonists, SLW131 (**10**) and SLW132 (**21m**), developed by converting two weakly active antagonists into single- or double-digit nanomolar ligands with minimal modifications. The thiadiazoledioxide **10** was derived from the CCR7 antagonist Cmp2105 by removing a methyl group from the benzamide moiety, while the squaramide **21m** was obtained from the CXCR1/CXCR2 antagonist and clinical candidate navarixin by replacing the ethyl substituent by a *tert*-butyl group to engage a lipophilic subpocket. We show that **10** and **21m** qualify to probe CCR7 biology both in recombinant cells and in the endogenous signaling environment of immune cells. Our novel probes are expected to facilitate the design of next-generation intracellular CCR7 ligands and serve as molecular tools to interrogate CCR7 biology in human and murine endogenous settings.

**Entry for the Table of Contents:** 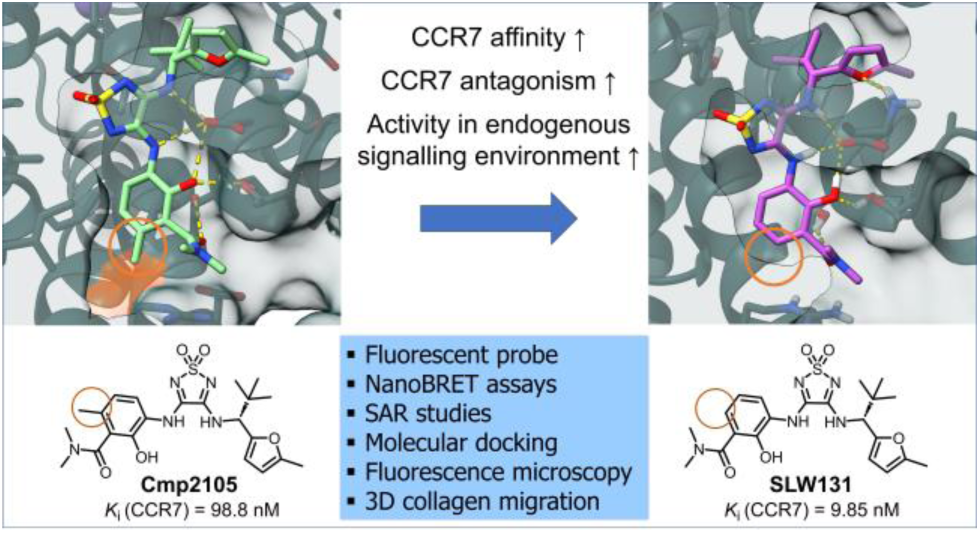

## INTRODUCTION

G-protein coupled receptors (GPCRs) are of high importance for drug development, as they represent more than 30% of the targets of currently used medications.^1,2^ Historically, the majority of GPCR ligands have targeted the orthosteric site, which is located within a conserved helical bundle accessible from the extracellular side.^1^ In addition to the orthosteric site, a conserved intracellular allosteric binding site (IABS) has recently been identified for several GPCRs. So far, the presence of an IABS has been verified through X-ray co-crystallography for the chemokine receptors CXCR2, CCR2, CCR7, and CCR9, as well as the beta-2-adrenergic receptor (β_2_-AR).^3–9^ Furthermore, the existence of a druggable IABS has been proposed for other GPCRs (*e.g.*, CCR1, CCR6, CXCR1). ^10,11^ The allosteric nature of these intracellularly binding ligands enables a novel dual-mechanism of action approach to GPCR modulation. This mode of action involves (i) the stabilization of the inactive conformation of the receptor, thereby leading to negative cooperativity with the orthosteric agonist, and (ii) the resulting steric hindrance blocks the binding of intracellular transducers and regulatory molecules, such as G-proteins, G protein-coupled receptor kinases, and β-arrestins, respectively. This strategy is particularly promising for GPCRs for which orthosteric antagonists are therapeutically ineffective.^12^ Notably, most of the receptors identified with an IABS are chemokine receptors, which are implicated in several cancers associated with poor prognosis but also in non-oncological conditions such as immune cell trafficking and homing as well as inflammatory processes.^13–16^ Thus, targeting the IABS in chemokine receptors represents a new avenue for therapeutic interventions in a variety of diseases. The CC chemokine receptor type 7 (CCR7) is a receptor specifically interacting with the C–C motif chemokines CCL19 and CCL21.^17^ Its primary expression occurs in diverse lymphoid tissues.^17,18^ CCR7 plays a vital role in lymphocyte and dendritic cell (DC) homing to secondary lymphoid organs and has been recognized as a potential therapeutic target in diseases such as rheumatoid arthritis, pathogenic infections and lymph node metastasis.^18^ Consequently, there is a clinical demand for the discovery and characterization of selective modulators capable of regulating CCR7 downstream signaling pathways. The absence of known orthosteric antagonists for CCR7 has posed challenges in the development of drugs targeting this receptor to date. However, a recent breakthrough by Jaeger *et al.* resolved the crystal structure of Cmp2105 (**1**, Figure 1) bound to the IABS of CCR7 through X-ray crystallography.^5^ They discovered that Cmp2105 (**1**) binds to the IABS similarly to vercirnon^1,19^ in CCR9 and Cmpd-15PA in the β_2_-AR.^7,20^ This, together with the observed inhibitory effect on CCL19 binding, identifies Cmp2105 (**1**) as an intracellular allosteric antagonist that may serve as a promising lead compound for the development of improved CCR7 antagonists. Additionally, Jaeger *et al.* screened known antagonists of other chemokine receptors and discovered, unexpectedly, that the established CXCR1/CXCR2 antagonist navarixin (**2**, Figure 1) also binds to CCR7. Docking studies suggested that navarixin (**2**) interacts with the IABS of CCR7 in a similar fashion as Cmp2105.^5^ Given their nearly identical substituents attached to the thiadiazoledioxide or squaramide core, these two compounds hold significant promise as lead structures for the development of novel intracellular CCR7 antagonists.

**Figure 1.**
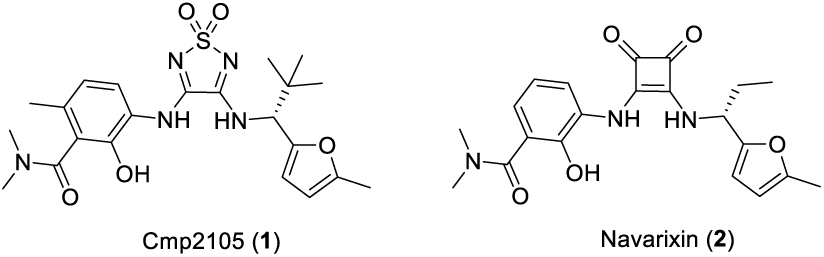
Chemical structures of the thiadiazoledioxide-based intracellular CCR7 antagonist Cmp2105 (**1**) and the squaramide-based CXCR1/CXCR2 antagonist navarixin (**2**).

To date, the characterization of CCR7 ligands targeting the IABS has focused on thermofluor stability assays and functional assays (e.g. cellular arrestin recruitment assays).^5,21^ Although thermofluor stability assays are a valuable tool for initial screening of protein-ligand interactions, they have several limitations. These limitations include: (i) applicability only to proteins that denature with heat, (ii) significant dependence on the specific fluorescent dye used, (iii) a substantial likelihood of misinterpretation, and (iv) a rather qualitative or at best semi-quantitative nature, making them unsuitable for accurate determination of binding affinities.

We recently developed squaramide-based fluorescent tracers targeting the IABS of CCR2 and CCR9.^22,23^ These tools have been effectively utilized in cell-free and cellular binding studies using the NanoBRET technology.^22,23^ In the present study, we report the development of a fluorescent thiadiazoledioxide-based intracellular CCR7 ligand derived from Cmp2105 (**1**). This molecular tool enabled to probe the binding of established and novel ligands to the IABS of CCR7 both in a cell-free and cellular environment. The most promising and selective CCR7 ligands identified in this screening were further characterized in a panel of functional assays in both recombinant and primary cells. This workflow allowed us to discover intracellular CCR7 antagonists with improved affinity and enhanced antagonistic behaviour compared to Cmp2105 (**1**), qualified as probes to investigate contribution of CCR7 to complex biological responses also in the endogenous signaling environment.

## RESULTS AND DISCUSSION

### Design and synthesis of a fluorescent probe targeting the IABS of CCR7

The co-crystal structure of human CCR7 in complex with the intracellular allosteric antagonist Cmp2105 (**1**) (PDB ID: 6QZH) provided the foundation for the structure-based design of a fluorescent tracer for the IABS of CCR7.^5^ The examination of this structure indicated that the *N*,*N*- dimethylamide moiety of Cmp2105 (**1**) is sufficiently solvent exposed to facilitate the attachment of a fluorophore for the development of a fluorescently labeled intracellular CCR7 ligand. To simplify the synthesis, we omitted the methyl group adjacent to the *N*,*N*-dimethylamide group. Our prior research on TAMRA-labeled ligands for other chemokine receptors, including CCR1, CCR2, CCR6, CCR9, CXCR1 and CXCR2, has shown that such ligands are effective molecular tools to investigate the binding of ligands to the IABS of GPCRs in both cell-free and, more importantly, cellular setups.^10,11,22–24^ Based on this experience, we aimed to attach a TAMRA fluorophore using a Cu-catalyzed Huisgen cycloaddition to a Cmp2105 analog. To visualize the correctness of this assumption, we conducted molecular docking studies with the triazole-based CCR7 ligand-linker conjugate **3** *post hoc* (Figure 2).

**Figure 2.**
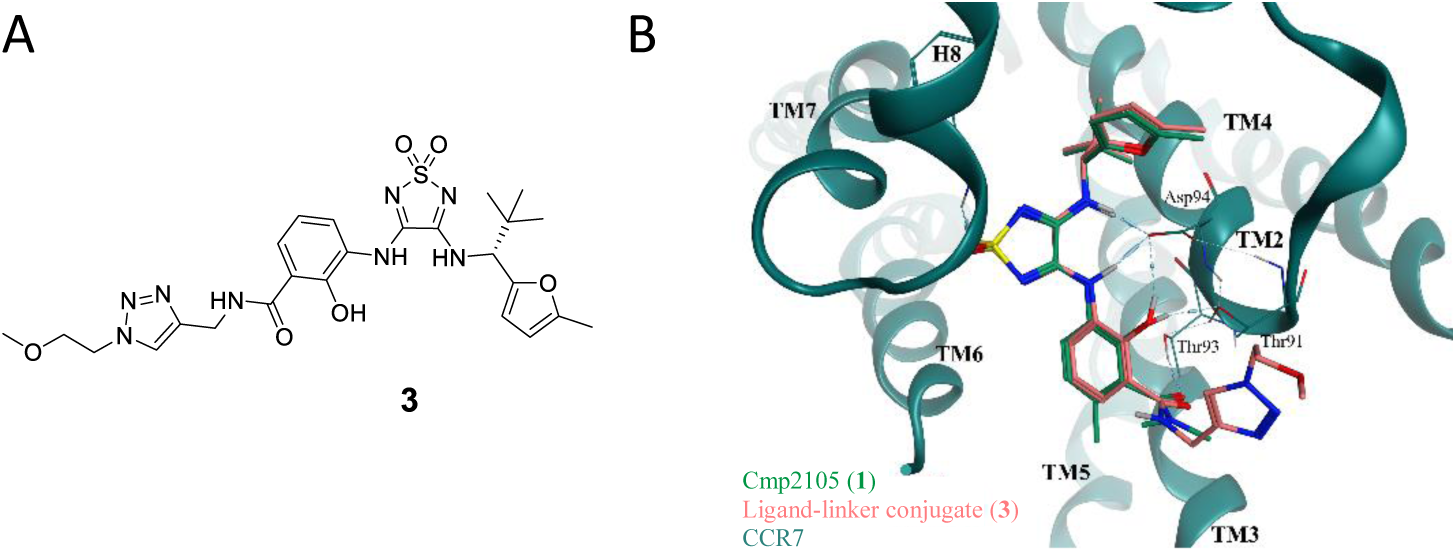
Design of a fluorescent ligand targeting the intracellular allosteric binding site of CCR7. A) Chemical structure of the triazole-based CCR7 ligand-linker conjugate **3** that was designed by introducing a linker which had already been successfully implemented in ligands targeting the IABS of other chemokine receptors. B) Overlay of the reported binding mode of Cmp2105 (**1**, green; PDB ID: 6QZH) with the predicted binding mode of the Cmp2105-derived ligand-linker conjugate **3** (salmon) in the IABS of CCR7 (green).

The synthesis of the designed fluorescent CCR7 probe **4** is summarized in Scheme 1. First, 3-nitrosalicylic acid (**5**) was converted into the corresponding propargyl amide **6** by an amide bond formation using PyBroP and DIPEA as coupling system to obtain the desired product. The reduction of the nitro group using tin(II) chloride monohydrate afforded the corresponding aniline **7**. In the next step, **7** was reacted with the readily available building block 3,4-dimethoxy-1,2,5- thiadiazole-1,1-dioxide to obtain the mono-substituted 1,2,5-thiadiazole-1,1-dioxide **8**. The subsequent second substitution reaction of intermediate **8** with (*R*)-2,2-dimethyl-1-(5- methylfuran-2-yl)propan-1-amine provided the clickable CCR7 ligand **9**. This ligand was then subjected to a Cu(I)-catalyzed azide−alkyne cycloaddition (CuAAC) with an azido-functionalized TAMRA analog to furnish the desired fluorescent tracer **4**.

**Scheme 1.**
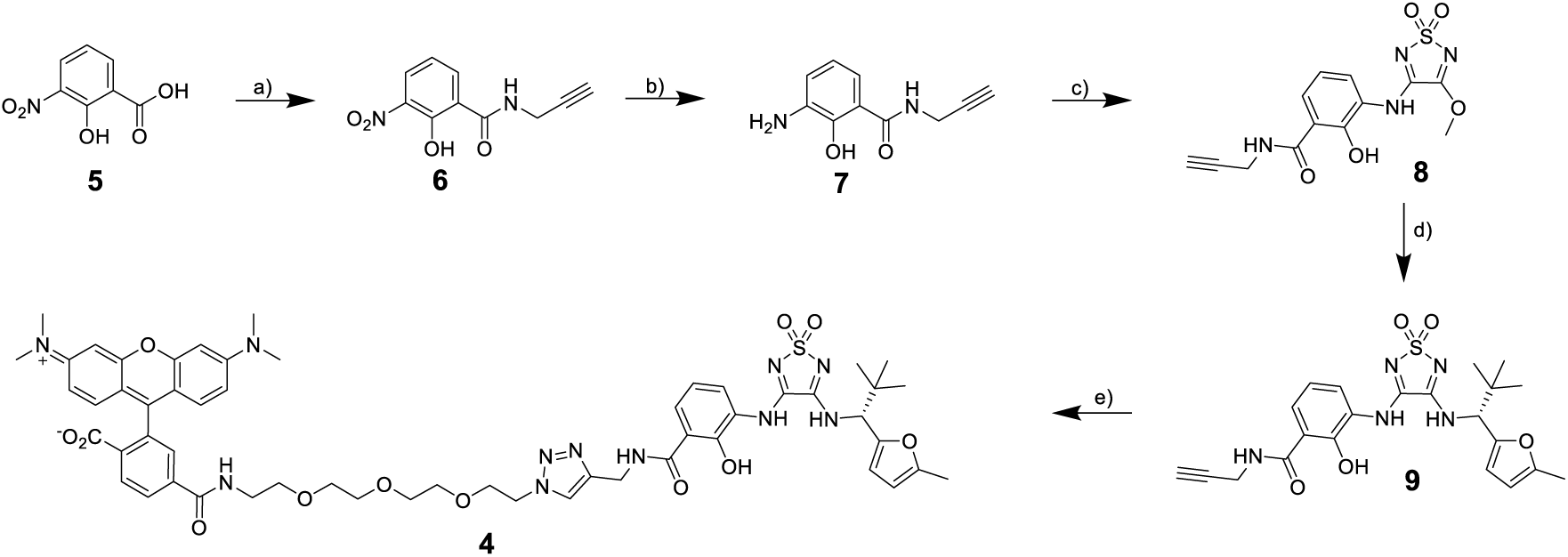
Synthesis of the fluorescent tracer **4**. *Reagents and conditions*: a) Propargylamine, PyBroP, DIPEA, CH_2_Cl_2_, rt, overnight, 79% yield. b) SnCl_2_ · H_2_O, MeOH, reflux, 3 h, quantitative yield. c) 3,4-Dimethoxy-1,2,5-thiadiazole-1,1-dioxide, MeOH, rt, 3 days, quantitative yield. d) (*R*)-2,2-Dimethyl-1-(5-methylfuran-2-yl)propan-1-amine hydrochloride, DIPEA, MeOH, rt, 7 days, 66% yield. e) 6-TAMRA-PEG_3_-azide, CuSO_4_·5 H_2_O, sodium ascorbate, TBTA, water/*tert*-BuOH/DMF mixture (1:1:1 (*v/v/v*)), rt, 6 h, 52% yield.

### NanoBRET Assay Development for CCR7

To evaluate the capacity of Mz437 (**4**) to serve as a molecular tool for studying ligand binding to the IABS of CCR7, we established a NanoBRET-based binding assay (Figures 3A-G and S1A-G). To this end, we fused the small bioluminescent Nanoluciferase (Nluc)^25^ either directly to the intracellular C terminus of CCR7 or with a GSSG linker between CCR7 and the Nluc-tag (Figures 3A and S1A), in a similar manner as reported for CCR2, CCR6 or CCR9.^10,20,22^ Successful surface expression of the CCR7-Nluc fusion proteins, hereafter referred to as CCR7_Nluc and CCR7_GSSG_Nluc, was confirmed by ELISA (Figure S1Β). In first cell-free experiments with membranes from HEK293T cells transiently expressing CCR7_Nluc and CCR7_GSSG_Nluc, respectively, we observed a slightly larger assay window for CCR7_GSSG_Nluc (Figure S1E), and thus used this construct for further investigations. In cell-free saturation binding experiments using CCR7_GSSG_Nluc membranes, Mz437 (**4**) was characterized as a high affinity intracellular CCR7 ligand with an equilibrium dissociation constant (*K*_D(eq.)_) of 212 nM (Figures 3B and S1F). Membrane-based kinetic binding experiments indicated both fast association (*k*_on_ = 4.28 × 10^6^ M^−1^ min^−1^) and dissociation (*k*_off_ = 2.23 × 10^−1^ min^−1^, *t*_r_ = 4.50 min) of the receptor-ligand complex (Figure 3D, Table S1). The derived kinetic dissociation constant (*K*_D(kin.)_) of 52.2 nM is in a similar range as the detected equilibrium *K*_D_ (*vide supra*). Then, we applied Mz437 (**4**) in a cell-free competition binding setup, to report the binding of nonfluorescent ligands to the IABS of CCR7. Using this setup, we detected a two-digit nanomolar affinity for thiadiazoledioxide Cmp2105 (**1**, *K*_i_ = 98.8 nM). The approximately 3-fold difference between the reported IC_50_ value of 35 nM, which was determined by means of a membrane-based competition experiment with the radioactively labeled orthosteric agonist CCL19, and the *K*i value from our NanoBRET-based competition binding assay can be rationalized by the strongly different concepts and setups of the applied GPCR binding assays. A much weaker affinity was detected for the squaramide navarixin (**2**, *K*_i_ = 7790 nM), which is in line with the results from dose response thermal shift assays reported by Jaeger *et al*.^5^ For the thiadiazoledioxide-based linker-ligand conjugate **3** (see Scheme S1, Supporting Information for synthetic details) that shares the same CCR7-binding unit as our fluorescent tracer Mz437 (**4**), we determined a *K*i value of 504 nM. This underlines that all changes at the scaffold of Cmp2105 (**1**) that were done in the course of the design of our fluorescent ligand (*i.e.*, attachment of a linker at the benzamide N-atom, switch from a tertiary to a secondary benzamide, removal of the methyl group in position 6 of the benzamide) were well tolerated. To probe the effect of the removal of the methyl group in position 6 of the benzamide, we synthesized and tested the respective desmethyl analogue **10** (Figure 3D, see experimental section for synthetic details) of Cmp2105 (**1**). Intriguingly, we observed an approximately 10-fold increased CCR7 affinity for **10** (*K*_i_ = 9.85 nM) compared to the parent compound Cmp2105 (**1**). Initial selectivity profiling, using previously reported NanoBRET assays to detect intracellular binding to the closely related chemokine receptors CCR2 and CCR9, indicated that **10** displays an improved CCR7 selectivity compared to Cmp2105 (Table S2, Supporting Information). In a next step, we assessed the suitability of our cell-free NanoBRET competition binding assay for high-throughput screening (HTS) approaches via a Z’-factor determination according to Zhang et *al.*^26^ These investigations resulted in a Z’-factor of 0.73 (Figure 3E), thus indicating that our cell-free CCR7 binding assay can be considered as excellent and suitable for HTS approaches.

**Figure 3.**
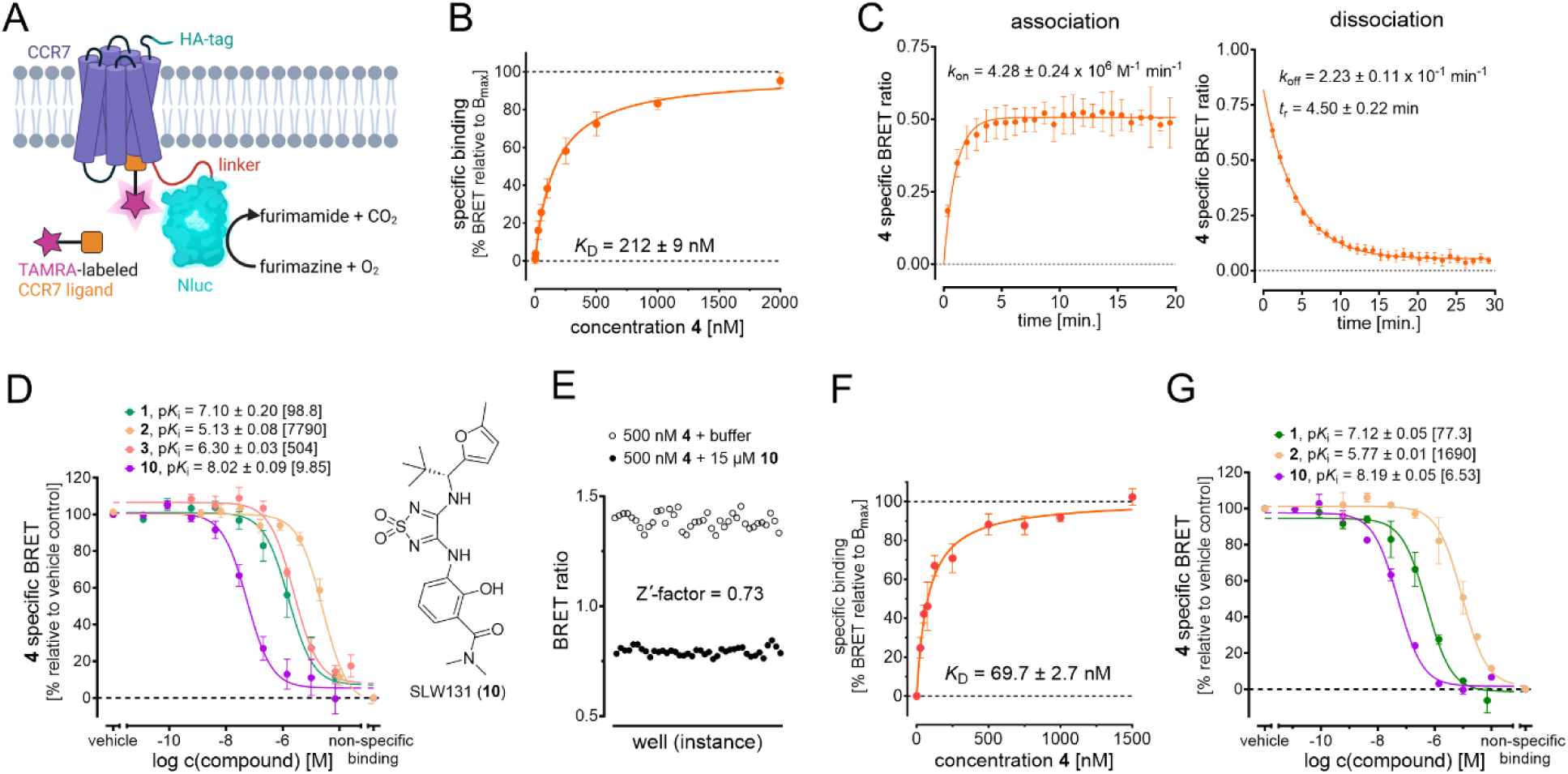
Application of Mz437 (**4**) as a fluorescent tool for cell-free and cellular NanoBRET- based binding studies targeting the IABS of CCR7. A) Cartoon representation of the NanoBRET strategy to detect binding to the IABS of CCR7. B) Specific saturation binding curve of the fluorescent tracer **4** in a NanoBRET-based assay using CCR7_GSSG_Nluc membranes (mean ± SEM, triplicate measurement, n = 3). C) Kinetic binding studies at room temperature. Representative association and dissociation curves with **4** (250 nM) using CCR7_GSSG_Nluc membranes (duplicate measurement). D) Competition binding curves and detected p*K*_i_ values (mean ± SEM, triplicate measurement, n ≥ 3) for the intracellular CCR7 antagonist Cmp2105 (**1**),^5^ the CXCR1/CXCR2 antagonist navarixin (**2**) with reported off-target binding to CCR7,^5^ the linker-ligand conjugate SLW151 (**3**), and **10** (chemical structure on the right), which is the desmethyl analogue of Cmp2105 (**1**). Data was obtained with **4** (500 nM) and CCR7_GSSG_Nluc membranes. *K*_i_ values are given in square brackets. E) Z’-factor determination for the cell-free NanoBRET-based setup using Mz437 (**4**, 500 nM) and CCR7_GSSG_Nluc membranes (36-fold determination). F) Saturation binding curve of Mz437 (**4**) in a cellular NanoBRET-based experiment (mean ± SEM, quadruplicate measurement, n = 4) using live HEK293T cells transiently expressing CCR7_GSSG_Nluc. G) Cellular competition binding curve and p*K*_i_ values (mean ± SEM, quadruplicate measurement, n ≥ 3) for Cmp2105 (**1**), navarixin (**2**), and **10** obtained with **4** (500 nM) and live HEK293T cells transiently expressing CCR7_GSSG_Nluc. *K*_i_ values are given in square brackets.

After having shown that Mz437 (**4**) is a highly valuable tool for studying ligand binding to the IABS of CCR7 in a high-throughput manner under cell-free conditions, we aimed for transferring our NanoBRET-based assay protocol to a live cell environment. Methods to directly monitor ligand binding to a specific target or target site in a cellular environment, also referred to as cellular target engagement, are highly important in early drug discovery. This is especially important for intracellular target proteins or transmembrane proteins with an intracellular binding site, such as the IABS of GPCRs. To the best of our knowledge, no small-molecule tracers have been reported thus far that enabled cellular binding assays for the IABS of CCR7. In a cellular saturation binding assay setup, using live HEK293T cells overexpressing CCR7_GSSG_Nluc, we detected a *K*_D_ value of 67.7 nM for our fluorescent CCR7 tracer Mz437 (**4**, Figure 3F). With this, we demonstrated that Mz437 (**4**) is able to pass the cell membrane of live cells and bind to the IABS of CCR7 in a live cell environment. Using Mz437 (**4**) as a tracer for cell-based competition binding assays, we detected a two-digit nanomolar affinity for Cmp2105 (**1**, *K*_i_ = 77.3 nM), a strongly reduced affinity for navarixin (**2**, *K*_i_ = 1690 nM), and a single-digit nanomolar affinity for **10** (*K*_i_ = 6.53 nM, Figure 3G). These results are (i) highly consistent with the affinity data from our cell- free experiments (Figure 3B); (ii) clearly showing the suitability of Mz437 (**4**) as a tracer to study cellular target engagement for the IABS of CCR7, and; (iii) highlighting **10** as an intracellular CCR7 ligand with significantly improved affinity in a cell-free and cellular environment, as compared to the prior gold standard Cmp2105 (**1**).

### Docking studies of antagonist 10 vs. Cmp2105 (1)

Cell-based and cell-free competition binding assays showed an unexpected 10-fold affinity improvement of the desmethylated derivative **10** compared to the parent compound Cmp2105 (**1**). Intrigued by such a shift in the binding affinity, we performed molecular docking calculations to rationalize the effect of this single methyl group removal. Not surprisingly, we obtained almost identical binding modes for **10** and Cmp2105 (**1**). Taking a closer look at the crystal structure of Cmp2105 in CCR7’s IABS (PDB ID: 6QZH), we noted that the methyl group of interest is solvent-exposed. We therefore hypothesized that the binding affinity was affected by the different interactions between the ligands and the solvent in that region. To test the hypothesis, we ran SZMAP calculations to predict whether any water molecules were potentially influenced by the introduction of the methyl group. The SZMAP calculations with Cmp2105 (**1**) and desmethylated **10** showed that the main difference between the two predicted water maps lies in the region highlighted as a red solid surface in Figure 4A. Exactly in that location we observed a region in the displacement grid where the methyl group of Cmp2105 displaces an energetically favorable water molecule (predicted SZMAP free energy value −5.97 kcal/mol) from the *apo* grid (Figure 4A; sampled orientations of the water molecule shown as sticks). This is corroborated by stabilization calculations (depicted in green mesh in Figure 4A), which indicated that this water molecule has a stabilizing effect on the protein-ligand complex. Displacement of such a stabilizing water molecule could in theory be counterbalanced by newly emerging energetically favorable interactions made by functional groups of a ligand — yet not via a methyl group, as in Cmp2105 (**1**). Consistent with this assumption, in the protein-ligand complex of CCR7 and **10**, the corresponding energetically favorable water molecule at the equivalent position is not displaced by the ligand and has essentially the same calculated free energy value of −6.08 kcal/mol (Figure 4B). Therefore, we hypothesize that the methyl group of Cmp2105 (**1**) displaces an energetically favorable water molecule, which – when present – stabilizes the protein-ligand complex. Since the methyl group of Cmp2105 (**1**) cannot compensate for the lost interactions, this would lead to a net loss in binding free energy. When the desmethylated ligand **10** is bound, a water molecule at this position would be left unperturbed, an explanation consistent with the experimentally observed increased potency.

**Figure 4.**
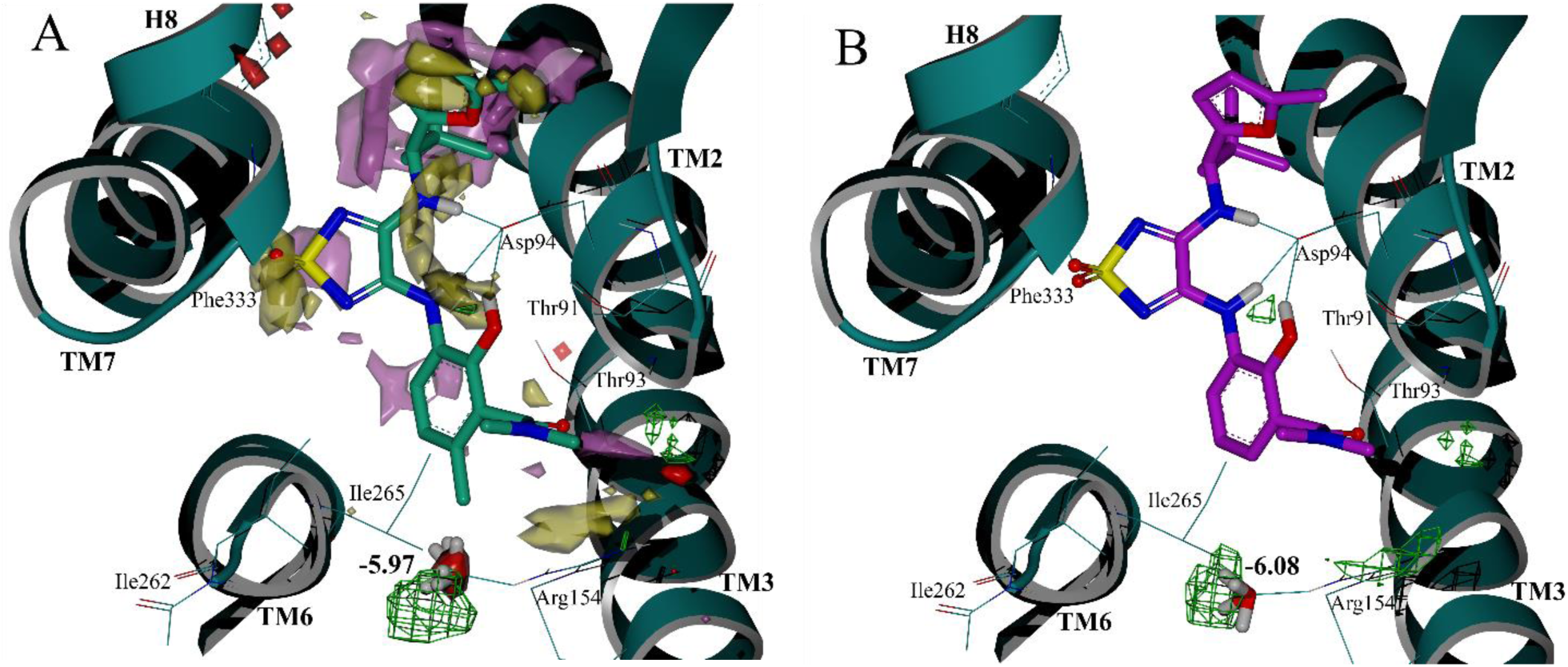
A) SZMAP grid calculations in the CCR7 IABS (PDB ID: 6QZH), shown as green ribbon: ligand displacement water map of Cmp2105 (**1**, green carbons). Residues within 4.5 Å from the methyl group of interest are shown as wires. Solid surfaces correspond to the water regions having negative (yellow) or positive (purple) neutral difference free energies. Water molecules are expected to enhance binding affinity when located in negative regions and decrease it when located in positive regions. Calculated difference between the Cmp2105 (**1**) and **10** grid values is highlighted as red solid surface. Region where water stabilizes the protein-ligand complex is shown in green mesh. Sampled water molecule orientations are shown as sticks. This water molecule is hypothesized to be displaced by the methyl group of Cmp2105 (**1**), but not by **10**, and has a predicted energy value of −5.97 kcal/mol. B) SZMAP grid calculations in the CCR7 IABS (PDB ID: 6QZH), shown as green ribbon: protein-ligand complex water map of **10** (magenta carbons). The water molecule corresponding to the one displaced by Cmp2105 (**1**) is shown as sticks. Free energy value of −6.08 kcal/mol is similar to the energy value of the displaced water molecule in the Cmp2105 (**1**) complex (−5.97 kcal/mol). Stabilization grid is shown in green mesh.

### SAR study for thiadiazoledioxide- and squaramide-based intracellular CCR7 antagonists

To further expand the knowledge on the structure-activity relationships (SARs) of intracellular allosteric CCR7 antagonists, a series of novel Cmp2105 and navarixin analogues were synthesized as outlined in Scheme 2. The arylalkylamino part was synthesized in a similar fashion as previously published (Scheme 2A).^10,11^ Starting from the aldehydes **11a-e**, the imines **13a-e** were created by a condensation reaction with Ellmańs sulfinimide ((*S*)-2-methylpropane-2-sulfinamide, **12**)^27,28^ using titanium ethoxide as catalyst. The imines were then subjected to a Grignard type reaction with ethyl-, isopropyl- or *tert*-butyl magnesium chloride to obtain the products **14a-g**. This diastereoselective reaction produced exclusively the desired diastereomers (see Experimental Section for details), with the exception of the less sterically hindered ethyl derivative **14b**, which was obtained as a mixture of diastereomers. In the last step, the chiral auxiliary was removed by treatment of **14a-g** with hydrochloric acid to obtain the free amine hydrochlorides **15a-g**. The benzamide part was synthesized as summarized in Scheme 2B by the reaction of 3-nitrosalicylic acid (**5**) with the corresponding amines in a PyBroP-mediated amide coupling. The newly synthesized amides **16a-c** were then treated with tin(II)chloride monohydrate to obtain the free anilines **17a-c**. In the next step, the benzamide moiety was coupled to either 3,4-dimethoxy-1,2,5- thiadiazole-1,1-dioxide or dimethyl squarate to afford the intermediates **18a-c** and **19a-b**. Lastly, the synthesis of the desired Cmp2105 and navarixin analogues was finalized by the introduction of the second amine component by treatment of the corresponding thiadiazole-1,1-dioxide and squaramide intermediates with **15a-g** or commercially available amines to generate the final compounds **20a-e** and **21a-m**. The final compounds are summarized in Table 1 and 2, and Table S3 (Supporting Information).

**Table 1.**
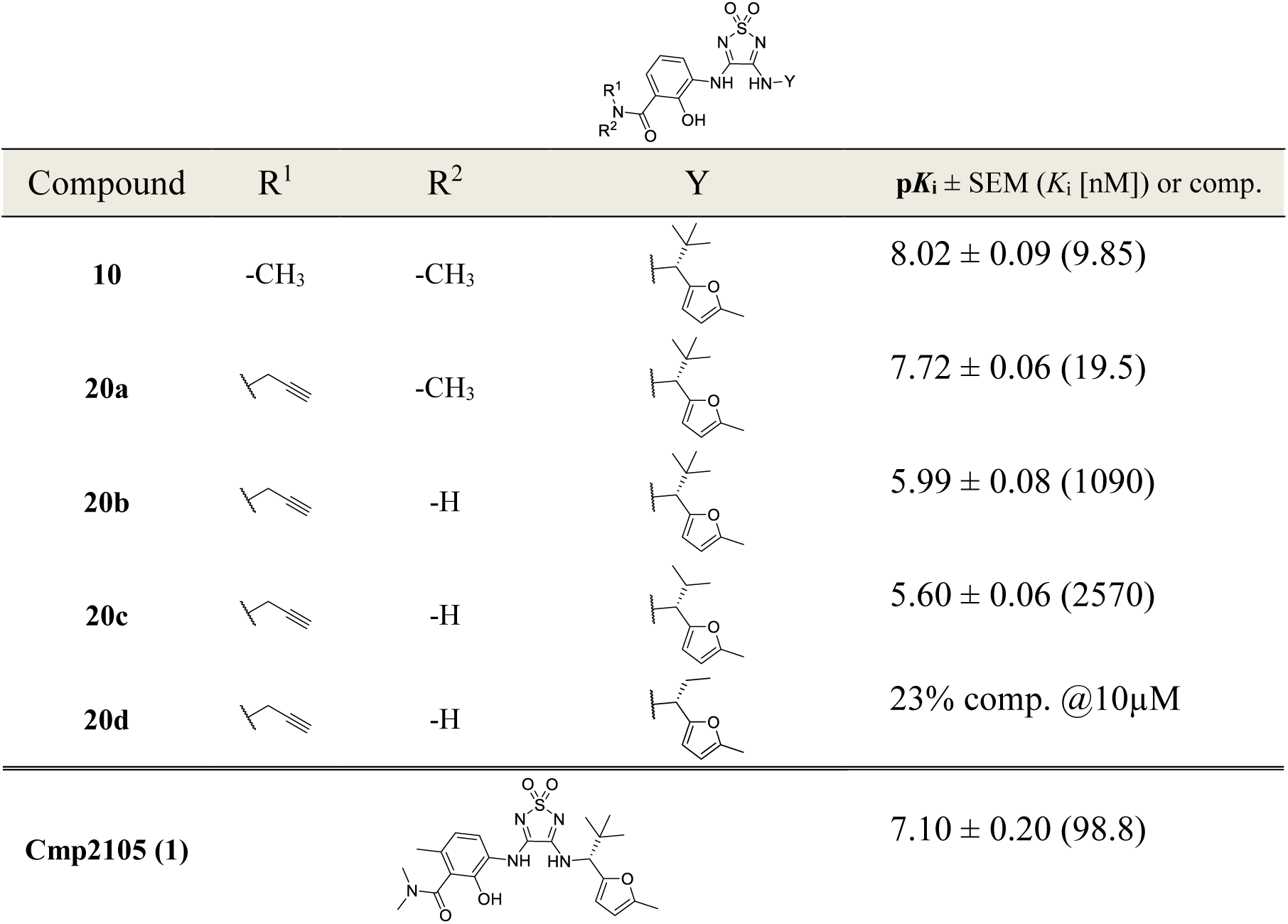
Overview of the synthesized thiadiazole-1,1-dioxide-based Cmp2105 analogues and their respective p*K*_i_ values (mean ± SEM, triplicate measurement, n ≥ 3) obtained from our cell-free NanoBRET competition binding assay. Data was obtained with **4** (500 nM) and CCR7_GSSG_Nluc membranes. *K*_i_ values are given in brackets.

|  |  |  |  |  |
| --- | --- | --- | --- | --- |
| Compound | R<sup>1</sup> | R<sup>2</sup> | Y | $pK_i \pm \text{SEM}$ ($K_i$ [nM]) or comp. |
| **10** | -CH<sub>3</sub> | -CH<sub>3</sub> | | $8.02 \pm 0.09$ (9.85) |
| **20a** | | -CH<sub>3</sub> | | $7.72 \pm 0.06$ (19.5) |
| **20b** | | -H | | $5.99 \pm 0.08$ (1090) |
| **20c** | | -H | | $5.60 \pm 0.06$ (2570) |
| **20d** | | -H | | 23% comp. @10μM |
| **Cmp2105 (1)** | | | | $7.10 \pm 0.20$ (98.8) |

**Scheme 2.**
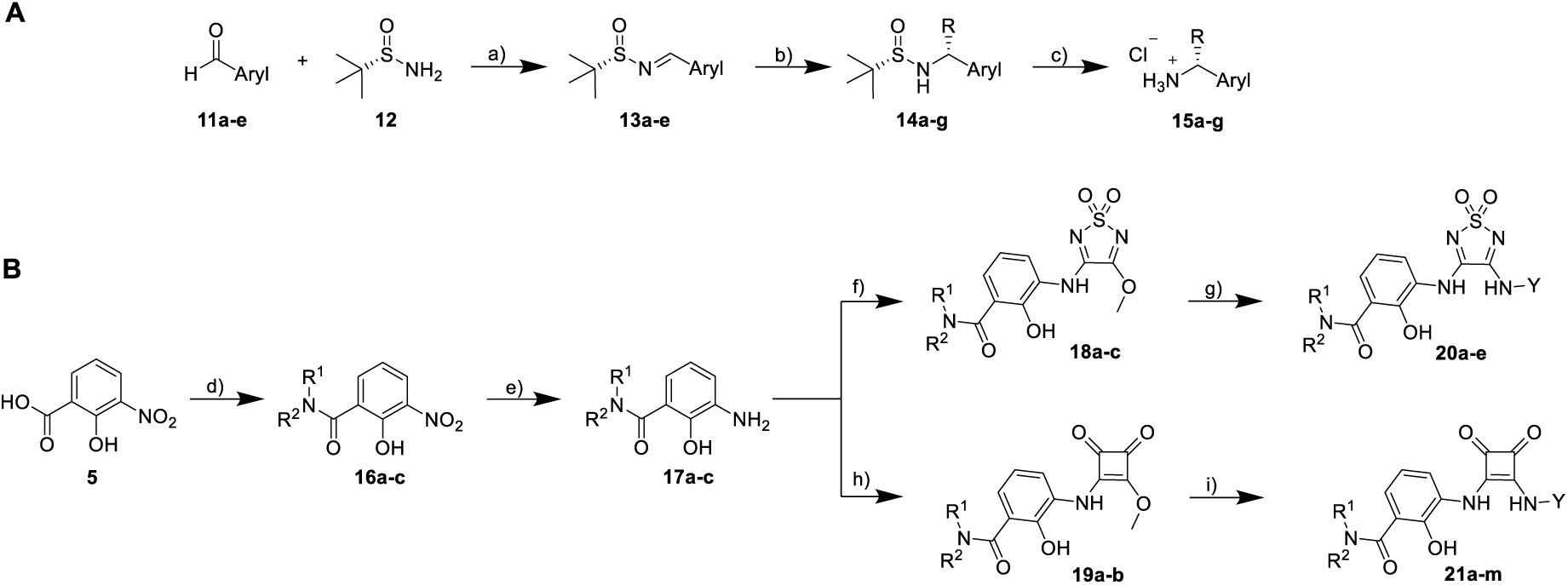
Synthesis of novel ligands for SAR studies. *Reagents and conditions*: A) a) Ti(OEt)_4_, DCM, rt, overnight, quantitative yield. b) Respective alkylmagnesium chloride, THF, rt, 3 days, 5 - 54% yield. c) HCl, Et_2_O, 0 °C, 1 h, quantitative yield. B) d) Respective amine, PyBroP, DIPEA, CH_2_Cl_2_, rt, overnight, 26 - 95% yield. e) SnCl_2_ · H_2_O, MeOH, reflux, 3 h, quantitative yield. f) **17a-c**, 3,4-dimethoxy-1,2,5-thiadiazole-1,1-dioxide, MeOH, rt, 3 days, quantitative yield. g) Respective amine hydrochloride, DIPEA, MeOH, rt, 7 days, 5 - 66% yield. h) **17a-c**, dimethyl squarate, MeOH, rt, 3 days, quantitative yield. i) Respective amine hydrochloride, DIPEA, MeOH, rt, 7 days, 5 - 66% yield.

The screening of the thiadiazole-1,1-dioxide series in our cell-free NanoBRET-based CCR7 competition binding assay provided important SARs (Table 1). The replacement of one methyl group of the tertiary amide in **10** by a propargyl residue yielded compound **20a** with a *K*_i_ value of 19.5 nM. This modification was well tolerated, resulting only in a 2-fold reduced binding affinity compared to **10** (*K*_i_ = 9.85 nM) and, notably, a 5-fold improvement in affinity relative to Cmp2105 (**1**, *K*_i_ = 98.5 nM). As the propargyl group provides a valuable functional handle for the creation of bifunctional tools such as fluorescent ligands and targeted protein degraders, we chose to retain the propargyl residue at the R^1^ position throughout this SAR study. The subsequent conversion of the tertiary amide in **20a** to a secondary amide in **20b** (*K*_i_ = 1090 nM) led to a 56-fold decreased binding affinity. Interestingly, when the hydrophobicity and bulkiness of the *tert*-butyl moiety realized in **20b** was reduced to isopropyl (**20c**) and ethyl (**20d**), the affinities decreased in the order **20b** > **20c** > **20d**.

To further investigate the SARs of our intracellular allosteric CCR7 antagonists, we performed a scaffold hopping approach to the navarixin-derived squaramide-based compounds **21a-m**. This enabled a straightforward access to a systematic variation of the furanylalkylamino part. However, the replacement of the chiral furanylalkylamino residue by various aliphatic, aromatic, and heterocyclic aromatic amines (see **21a-k**, Table S3, Supporting Information) resulted in a complete loss of affinity towards CCR7. The only exemption was the thiophene derivative **21l** featuring a *tert*-butyl residue at the chiral carbon. This modification resulted in low, yet quantifiable affinity towards CCR7 (*K*_i_ = 4200 nM, Table 2). When we screened the direct squaramide analog of **10**, compound **21m**, we observed the highest CCR7 affinity (*K*_i_ = 36.2 nM, Table 2) in the squaramide series. Similar to **10**, we detected a strongly reduced CCR2 and CCR9 affinities for **21m** (Table S2, Supporting Information). As a result, our SAR data for CCR7 underline the importance of a tertiary amide at the benzamide moiety and the *tert*-butyl moiety at the chiral carbon.

**Table 2.**
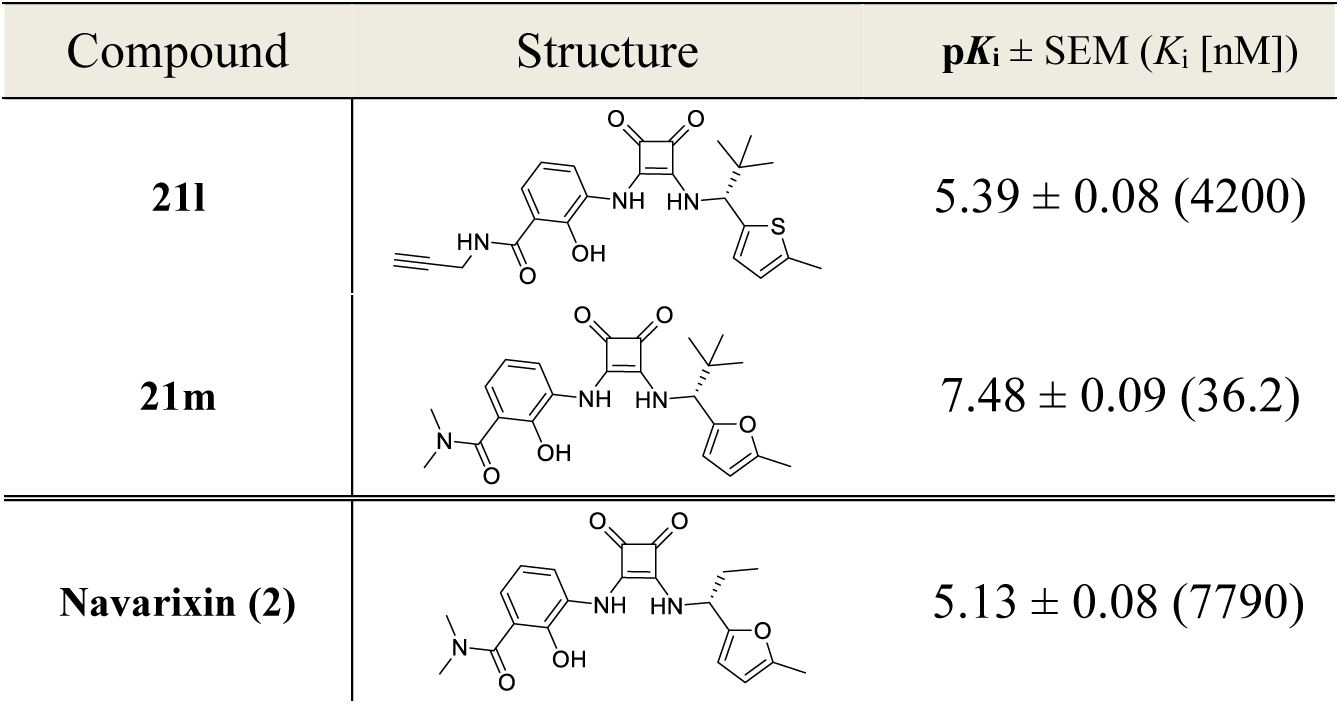
Overview of the synthesized squaramide-based navarixin analogues and their respective p*K*_i_ values (mean ± SEM, triplicate measurement, n ≥ 3) obtained from our cell-free NanoBRET competition binding assay. Data was obtained with **4** (500 nM) and CCR7_GSSG_Nluc membranes. *K*_i_ values are given in brackets.

| Compound | Structure | $pK_i \pm \text{SEM}$ ($K_i$ [nM]) |
| --- | --- | --- |
| **21l** | | $5.39 \pm 0.08$ (4200) |
| **21m** | | $7.48 \pm 0.09$ (36.2) |
| **Navarixin (2)** | | $5.13 \pm 0.08$ (7790) |

The critical role of the *tert*-butyl group is further confirmed by comparing the affinity data obtained for **21m** and navarixin (**2**). These compounds differ only in the residue at the chiral carbon (navarixin (**2**): (*R*)-ethyl; **21m**: (*R*)-*tert*-butyl). This modification led to a 215-fold decrease in binding affinity, which is in excellent agreement with the results from the thiadiazole-1,1-dioxide series. In our previous study on squaramide-based intracellular antagonists targeting the IABS of CCR6 and CXCR1, we identified the *tert*-butyl moiety at the chiral carbon as an absolute sweet spot.^10^ Docking studies indicated that the *tert*-butyl group fits perfectly into an aromatic cage in the IABS of CCR6, thereby facilitating crucial hydrophobic interactions. We visualized aromatic residues within a distance of 4.5 Å around the *tert*-butyl group of Cmp2105 (**1**) in CCR7’s IABS (PDB ID: 6QZH) and identified a lipophilic subpocket analogous to CCR6’s IABS (Figure 5). This lipophilic subpocket is formed by Val79^1x53^, the methyl group of Thr82^1x56^, Tyr83^1x57^, Leu97^2x43^, Tyr326^7x53^ and Phe333^8x50^. All of these residues are identical in the IABS of CCR6, except for Tyr83^1x57^, which corresponds to Phe71^1x57^ in CCR6. Similarly, to the CCR6 IABS, we speculate that this aromatic cage in CCR7’s IABS allows for an optimal shape complementarity with the *tert*-butyl group, resulting in a favorable binding affinity. As a result, this could be an important contribution to the increased binding affinity of **21m** relative to navarixin (**2**).

**Figure 5.**
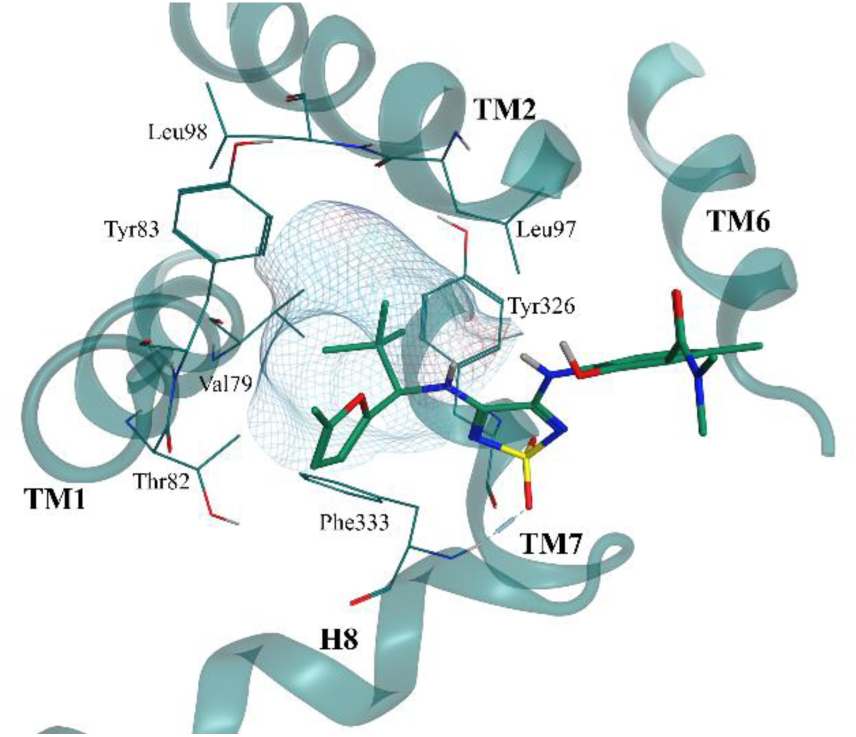
Visualization of the IABS of CCR7 (PDB ID: 6QZH) with a focus on the aromatic cage. Residues surrounding the *tert*-butyl group of Cmp2105 (**1**) are show in lines. Molecular surface (green mesh) highlights the shape complementarity of the receptor’s subpocket and the *tert*-butyl substituent.

### Fluorescence microscopy visualizes IABS targeting of intracellular CCR7 ligands

Development of IABS targeting CCR7 ligands demands complementary methods to determine intracellular target engagement. Based on the results from the SAR study, we selected the thiadiazole-1,1-dioxide **10** and the squaramide **21m** as well as Cmp2105 (**1**) as intracellular competitors for our newly developed fluorescent probe Mz437 using fluorescence microscopy. Distinct from and complementary to NanoBRET-based binding studies, which require the use of labelled receptors, fluorescence microscopy enables visualization of ligand binding to native receptors. We used fixed naïve and stable CCR7-HEK293 cells to obtain fluorescence profiles in costaining experiments with a CCR7-directed fluorescent antibody and the fluorescent probe Mz437. We found clear fluorescence overlap at the plasma membrane which is evident from both the merged images and the two co-incident sharp fluorescence peaks in line scan analysis (Figure 6). No detectable plasma membrane-associated fluorescence was observed in naive HEK293 cells, neither for the probe nor for the CCR7 antibody, entirely consistent with CCR7-specific labelling by Mz437. To test the competitive nature of this binding interaction, we added Mz437 to cells previously incubated with **10**, **21m**, Cmp2105 (**1**) or vehicle control for 60 minutes. We observed complete reduction of plasma membrane fluorescence by **10** and **21m**, consistent with specific and quantitative probe displacement from plasma membrane-resident CCR7. Cmp2105 (**1**), on the contrary, only partially promoted probe displacement in agreement with its lower CCR7 affinity as determined in the earlier NanoBRET-based binding assays. Neither of the intracellular antagonists diminished CCR7 recognition by the CCR7-directed fluorescent antibody. These data prove that Mz437 is a suitable tool for visualizing cellular target engagement of nonfluorescent intracellular CCR7 ligands, and reveal that **10** and **21m** clearly surpass the only available reference Cmp2105 (**1**) in binding to CCR7.

**Figure 6.**
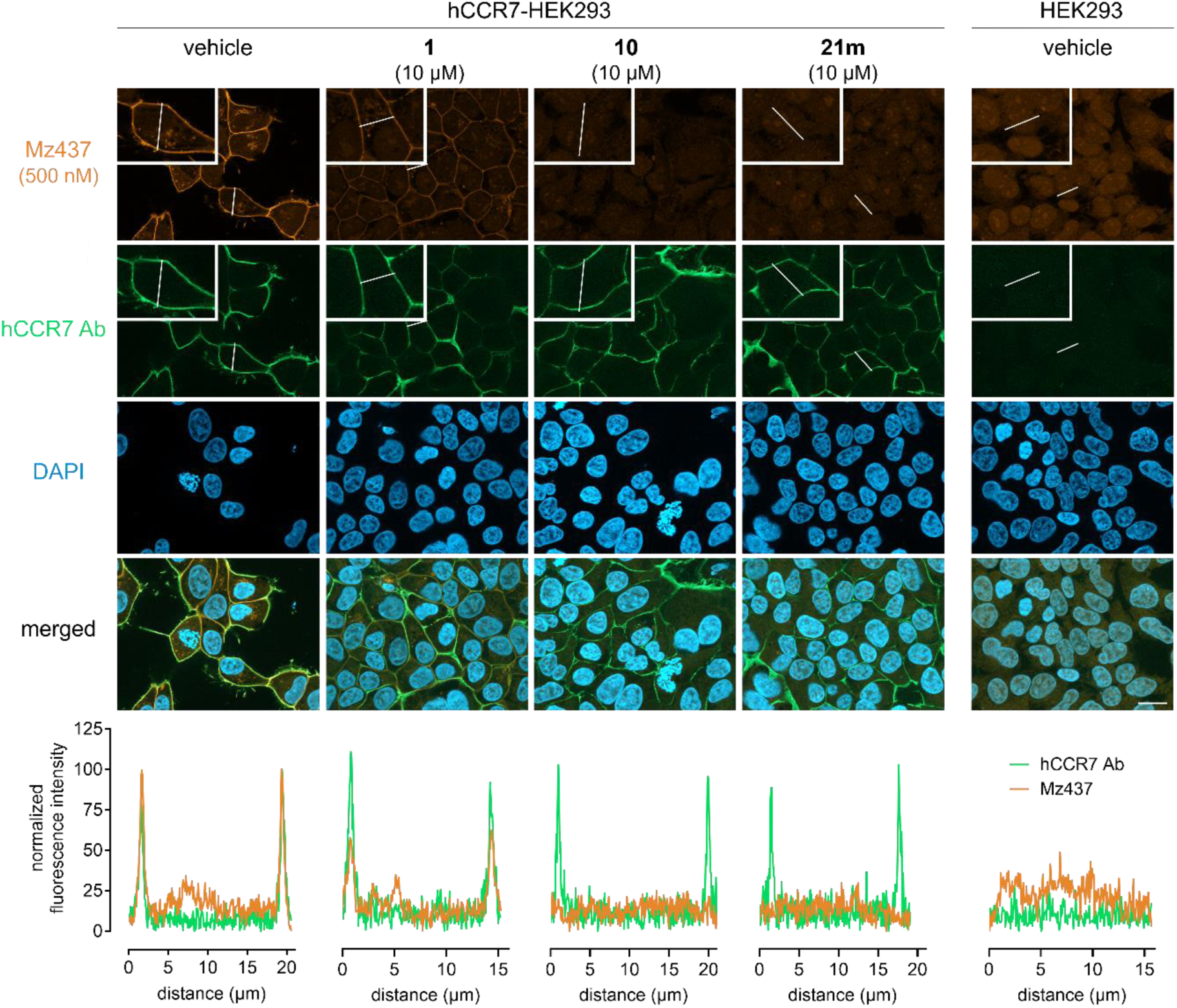
Intracellular CCR7 ligands competitively displace the fluorescent CCR7 probe Mz437. Naïve and stable HEK293-CCR7 cells were fixed and immunostained with an anti-CCR7 antibody and with the fluorescent CCR7 probe Mz437 as indicated. Fluorescence intensity profiles along the white lines were quantified by line scan analysis in the representative images, showing clear colocalization of Mz437 and receptor immunostaining on the plasma membrane as well as the displacement of the probe by the intracellular CCR7 ligands. Scale bar is 10 μm.

### 10 and 21m are efficacious CCR7 inhibitors in recombinant and primary cells

To investigate whether intracellular CCR7 binding also translates to inhibition of CCR7 function, we tested whether **10**, **21m**, and Cmp2105 (**1**) impede CCR7 interaction with signal transducing G proteins and signal-regulating arrestins. In HEK293T cells transiently expressing human CCR7 (hCCR7) increasing concentrations of CCL19 resulted in a concentration-dependent Go activation (Figure S2, Supporting Information). Pretreatment of cells with increasing concentrations of CCR7 binders prior to stimulation with CCL19 at its EC_85_ yielded single and double digit micromolar inhibition of Go by **10** and **21m**, respectively, but, to our surprise, no detectable inhibition by Cmp2105 (**1**) whatsoever (Figure 7A). We obtained comparable results assessing inhibition of CCL19-induced β-arrestin2 recruitment, in that **21m** surpassed the inhibitory potency of **10** by a factor of 17, while Cmp2105 (**1**) was again not able to mimic the inhibitory effects of the desmethyl compounds (Figure 7B and Figure S3, Supporting Information). To avoid any potential bias in our results due to functional selectivity of the synthesized ligands, we took advantage of holistic label-free whole cell biosensing based on detection of dynamic mass redistribution (DMR). Previously, we have shown that DMR is a powerful assay platform reporting whole cell activation upon GPCR stimulation regardless of the primary signaling pathway.^19,29,30^ Stimulation of HEK293 cells stably transfected to express hCCR7 with CCL19 provoked a feature-reach DMR response, that was characterized by a transient sharp rise that is followed by an initial rapid and a second slower descending phase, which is characteristic for many Gi-coupled receptors (Figure 7C). In this experimental setup, CCL19 activation of CCR7 was diminished by **10** and more so by **21m** but not by Cmp2105 (**1**) (Figure 7C). These data suggest that (i) lack of antagonistic efficacy of Cmp2105 (**1**) in Go activation and β-arrestin2 recruitment assays unlikely results from biased inhibition of only some of the G protein effector pathways downstream of CCR7, and (ii) that **10** and **21m** should qualify to probe contribution of CCR7 to biological responses also in the endogenous signaling environment. Because CCR7 is expressed in mature bone marrow-derived dendritic cells (BMDCs),^31^ we reasoned that **10** and **21m** should impede cell shape change induced by CCL19 but not by CXC chemokine or unrelated lipid stimuli. Indeed, pretreatment of BMDCs with both inhibitors significantly diminished CCL19-mediated cell shape changes but not those induced by CXCL12 and prostaglandin E_1_ (PGE_1_) acting via endogenous CXCR4 and EP2/EP4 receptors, respectively (Figure 7D). These data confirm CCR7 specificity of antagonist action in the label-free assay platform and imply suitability of both ligands to probe CCR7 biology also in primary cell environments. Consistent with specific inhibition of CCR7 function in the endogenous setting, **10** and **21m** blunted CCR7- but not CXCR4-dependent transmigration of primary murine CD4^+^ T cells (Figure 7E, F).

**Figure 7.**
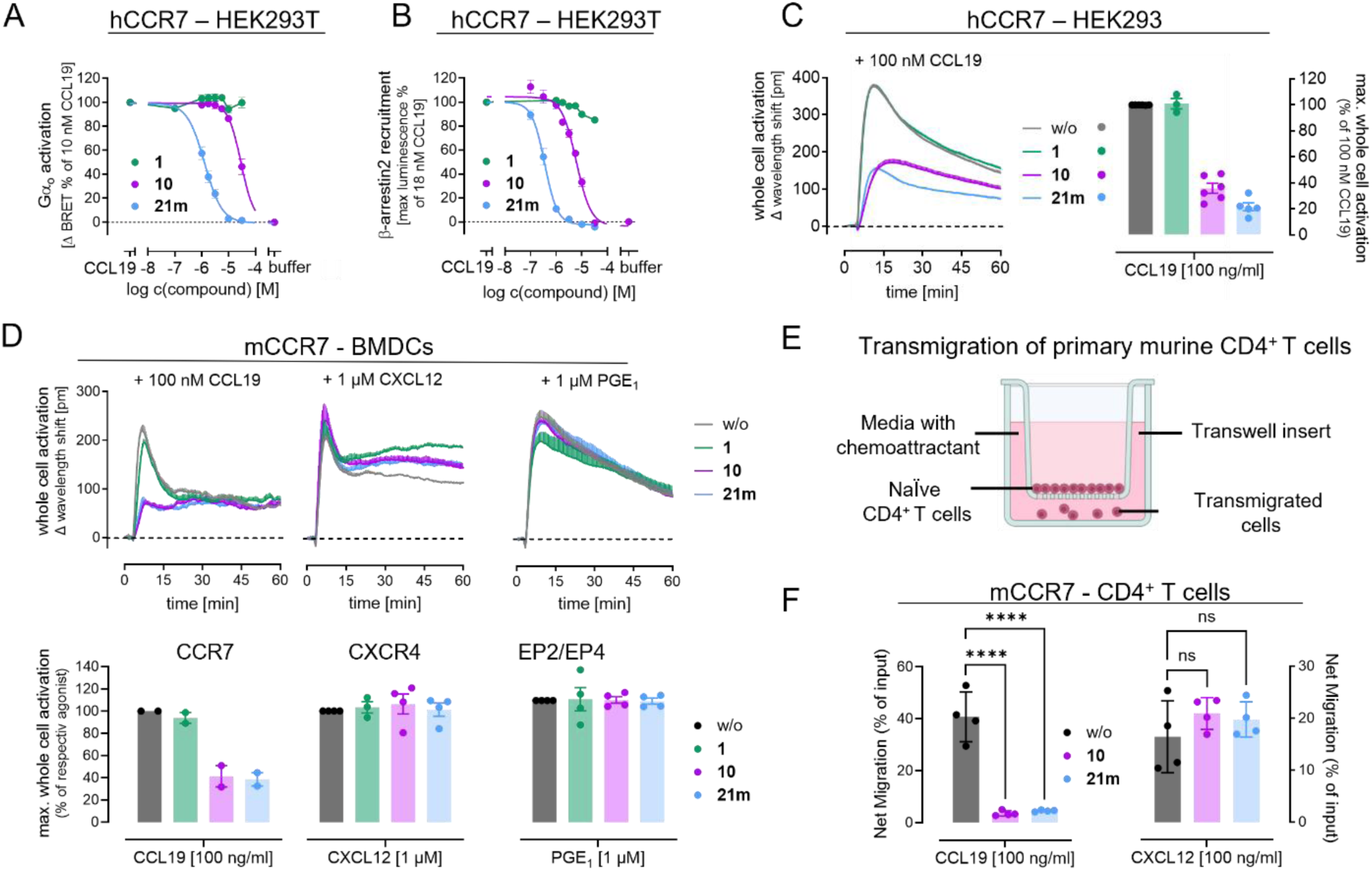
**10** and **21m** dampen CCR7 functionality in recombinant and primary cells. A) Inhibition of CCL19 (10 nM)-mediated Go activation, determined as BRET between Gαo-derived Gβγ (Venus-labelled) and the Gβγ effector mimic masGRK3ct-Nluc by CCR7 antagonists. MasGRK3ct-Nluc denotes the C-terminal fragment of the G protein-coupled receptor kinase isoform 3 fused to a myristic acid attachment peptide (mas) and NanoLuc luciferase (Nluc); **10** (pIC_50_ = 4.53 ± 0.03 [29.4 µM]) and **21m** (pIC_50_ = 5.89 ± 0.03 [1.28 µM]). B) Inhibition of CCL19- induced β-arrestin2 recruitment using a NanoBiT-based β-arrestin2 recruitment assay after stimulation of hCCR7 with 18 nM CCL19 (EC_25_); **10** (pIC_50_ = 5.22 ± 0.03 [6.0 µM]) and **21m** (pIC_50_ = 6.45 ± 0.04 [0.35 µM]). C) Representative whole cell activation profiles and their bar chart summaries of 100 nM CCL19 in the absence and presence of the indicated CCR7 ligands, obtained with label-free DMR whole cell biosensing in HEK293 cells stably expressing human CCR7. Label-free signatures are means + SEM of n= 3 experiments, performed in technical quadruplicates. Bar charts are the means ± SEM of n= 3 experiments. D) Representative DMR traces and their bar chart summaries evoked by 100 nM CCL19 in murine BMDCs after preincubation with the indicated CCR7 ligands. Label-free signatures are mean + SEM of technical triplicates. Summarized data are means ± SEM of n= 2-4 independent experiments. E) Experimental setup of transmigration assay. F) Inhibition of CCR7-mediated transmigration of murine CD4^+^ T cells towards a 100 ng/ml CCL19 or 100 ng/ml CXCL12 gradient by **10** and **21m**. Final concentration of **1**, **10**, and **21m** in C, D, and F is 10 µM. Statistical significance was calculated with a one-way ANOVA with Dunnett’s post-hoc analysis; ****, *P* < 0.0001; ns, not significant. Cartoon created with BioRender.com.

As CCR7-directed migration is an inherent feature of dendritic cells, we used 3D collagen matrices to probe the capacity of BMDCs to migrate along soluble gradients of CCL19 (Fig. 8A). Live cell video-microscopy allowed us to follow individual cells over time and extract migration parameters such as velocity and directionality (Fig. 8B, C). Manual and automized image analysis confirmed efficient reduction of directional migration towards higher CCL19 concentrations in the presence of **10** (Fig. 8C, D), consistent with the role of CCR7 as master regulator of DC migration. By contrast, pre-treatment with **21m** did not impact CCL19-guided BMDC migration (Fig. 8C, D). Although we cannot rule out that the observed functional difference between the two CCR7 ligands may be rationalized by insufficient target engagement in the complex 3D collagen matrix, our data clearly show the antagonistic efficacy of **10**. Moreover, they further emphasize context- and cell type-specific regulation of downstream CCR7 signaling as demonstrated before between T cells and DCs.^31^

**Figure 8.**
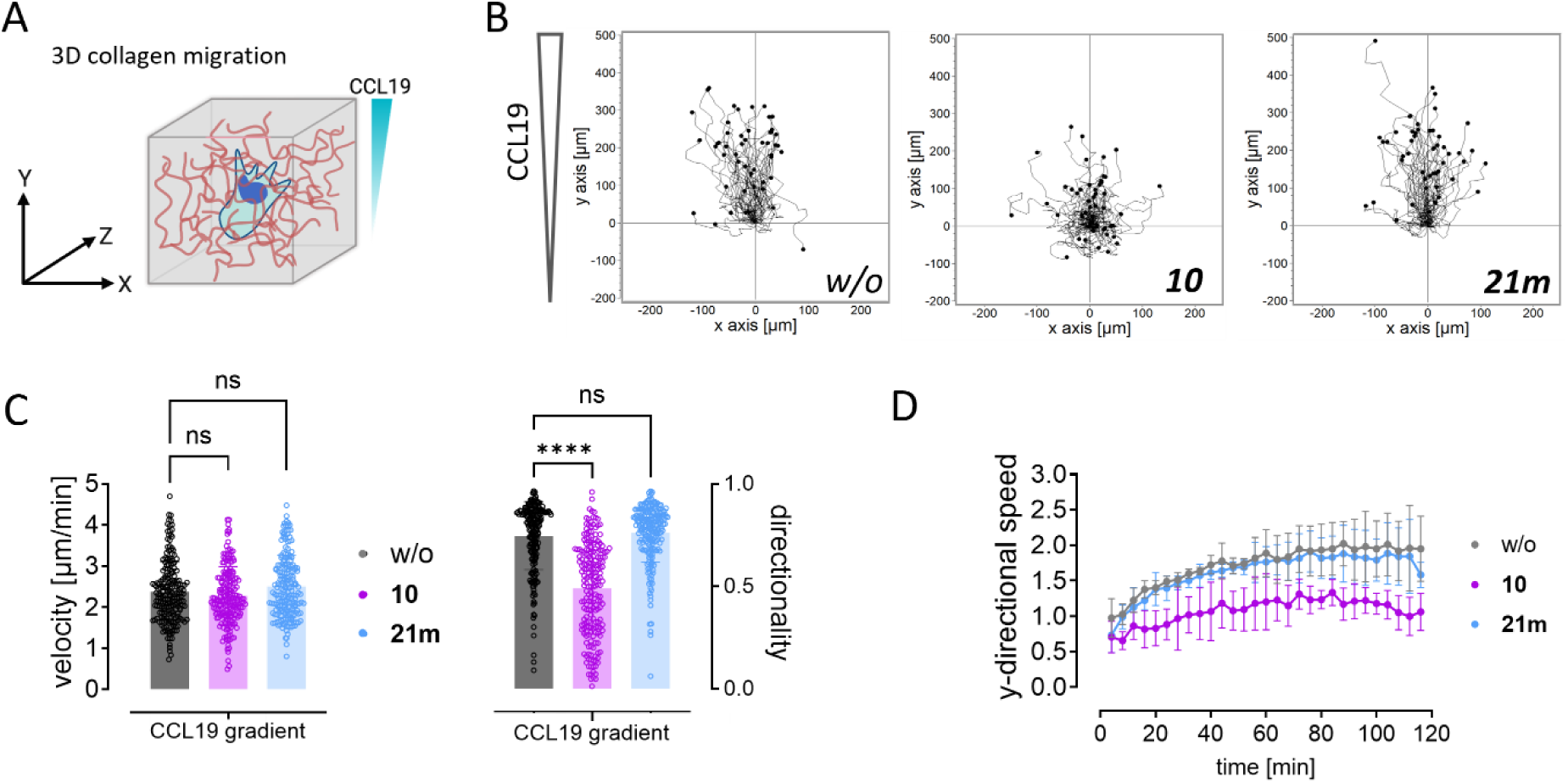
Reconstitution of DC migration in 3D collagen matrices. A) Schematic representation of experimental set-up. DCs are embedded within collagen fibers and exposed to a soluble gradient of CCL19. B) Individual tracks of single cells migrating in the absence (left) or presence of **10** (middle) and **21m** (right) each at 10 µM final concentration. Each track represents one cell pooled from two independent experiments. C) Quantification of migration velocity and directionality (defined as ratio of displacement to trajectory length). Each data point represents one cell. Graph shows mean values ± sd of four independent experiments with cells generated from two different mice. ****, *P* < 0.0001 (one-way ANOVA with Dunnett’s post-hoc analysis). ns, non-significant. D) Automated analysis of y-directional speed of all cells over time. Graph shows mean values ± sd of four independent experiments with cells generated from two different mice.

## CONCLUSION

Modulation of CCR7 is therapeutically attractive for two reasons: to dampen immune activation in autoimmune disorders and to improve immunity for anti-tumor targeting.^17,18^ In this work, we developed the TAMRA-derived fluorescent probe Mz437 (**4**) targeting the IABS of CCR7. We demonstrated its utility in enabling both cell-free and cellular NanoBRET-based binding studies. Additionally, we showed that Mz437 is an effective tool for visualizing intracellular target engagement of CCR7 through fluorescence microscopy. By employing Mz437, we identified the two intracellular CCR7 ligands **10** (hereafter dubbed SLW131) and **21m** (hereafter named SLW132). SLW131, a thiadiazoledioxide, was developed from Cmp2105 (**1**) by removing a methyl group from the benzamide core. In contrast, SLW132, a squaramide, was obtained from the CXCR1/CXCR2 antagonist navarixin by replacing its ethyl group with a *tert*-butyl group to target a lipophilic subpocket. Both compounds exhibit improved binding affinities and antagonistic properties compared to the prior gold standard intracellular CCR7 ligand Cmp2105 (**1**). In detail, CCR7 G protein activation and β-arrestin recruitment assays revealed that the increased affinity of SLW131 and SLW132 translates into an improved antagonistic behaviour compared to Cmp2105 (**1**), which was further corroborated by label-free whole cell biosensing based on detection of dynamic mass redistribution in both stable CCR7-HEK293 cells and primary BMDCs.

In conclusion, we successfully identified SLW131 and SLW132 as promising leads for future studies of CCR7 pharmacology and function in both recombinant and endogenous signaling environments. It is anticipated that these molecular tools will considerably advance our understanding of CCR7 functionality.

## ASSOCIATED CONTENT

### Supplementary Information

Electronic Supplementary Information (ESI) available: Supplementary Figures, Schemes, and Tables, experimental procedures and characterization data for all compounds, NMR spectra, and HPLC traces. (PDF)

## AUTHOR INFORMATION

### Author Contributions

The manuscript was written through contributions of all authors. All authors have given approval to the final version of the manuscript.

### Funding Sources

The new NMR console for the 500 MHz NMR spectrometer used in this research was funded by the Deutsche Forschungsgemeinschaft (DFG, German Research Foundation) under project number 507275896. D.B. was supported by Deutsche Forschungsgemeinschaft (DFG, German Research Foundation) under Germany’s Excellence Strategy EXC2151 (390873048, ImmunoSensation^2^). M.S. (Li 204/04) was supported by the Verband der Chemischen Industrie (VCI).

### Notes

Any additional relevant notes should be placed here.

## ACKNOWLEDGMENT

We acknowledge the assistance of the Flow Cytometry Core Facility at the Institute of Experimental Immunology, Medical Faculty, University of Bonn, for providing equipment. We thank Lara Toy for supporting the design of the graphical abstract.

## ABBREVIATIONS

BMDC: bone-marrow derived dendritic cell
BRET: bioluminescence-resonance energy transfer
BSA: bovine serum albumin
CCL19: C-C chemokine ligand 19
CCR1: C-C chemokine receptor 1
CCR2: C-C chemokine receptor 2
CCR6: C-C chemokine receptor 6
CCR7: C-C chemokine receptor 7
CCR9: C-C chemokine receptor 9
CD4: cluster of differentiation 4
CDCl_3_: deuterated chloroform
CXCL12: C-X-C chemokine ligand 12
CXCR1: C-X-C chemokine recptor 1
CXCR2: C-X-C chemokine receptor 2
CXCR4: C-X-C chemokine receptor 4
DC: dendritic cell
DCM: dichloromethane
DIPEA: *N,N-*diisopropylethylamin
DMF: *N*,*N*-dimethylformamide
DMR: dynamic mass redistribution
DMSO: dimethylsulfoxide
dr: diastereomeric ratio
EC: effective concentration
EDTA: ethylendiaminetetraacetic acid
ELISA: enzyme-linked immunosorbent assay
eq.: equivalent
ESI: electrospray ionization
Et_2_O: diethyl ether
GPCR: G-protein coupled receptor
HEK293: human embryonic kidney cells
HPLC: high-performance liquid chromatography
HR-MS: high-resolution mass spectrometry
IABS: intracellular allosteric binding site
IC: inhibitory concentration
*K*_i_: inhibitory constant
LR-MS: low-resolution mass spectrometry
MEM: Minimum essential medium eagle
MeOH: methanol
MOE: molecular open environment
NMR: nuclear magnetic resonance
PBS: phosphate buffered saline
ppm: parts per million
PyBroP: Bromo-tris-pyrrolidino-phosphonium hexafluorophosphate
SAR: structure- activity relationship
SEM: standard error of mean
SnCl_2_: tin(II)chloride
TAMRA: 5- carboxytetramethylrhodamine
TBTA: Tris[(1-benzyl-1*H*-1,2,3-triazol-4-yl)methyl]amin)
TFA: trifluoroacetic acid
THF: tetrahydrofurane
Tris: tris(hydroxymethyl)aminomethane.

## Notes

### Competing Interest Statement

The authors have declared no competing interest.

