## Supporting Information for "A fluorescent probe enables the discovery of improved antagonists targeting the intracellular allosteric site of the chemokine receptor CCR7"

### Table of Contents

SUPPLEMENTAL FIGURES, TABLES, AND SCHEMES ........ S4
EXPERIMENTAL SECTION ........ S10
NMR SPECTRA OF SYNTHESIZED COMPOUNDS ........ S32
HPLC CHROMATOGRAMS OF TARGET COMPOUNDS ........ S57
REFERENCES ........ S67

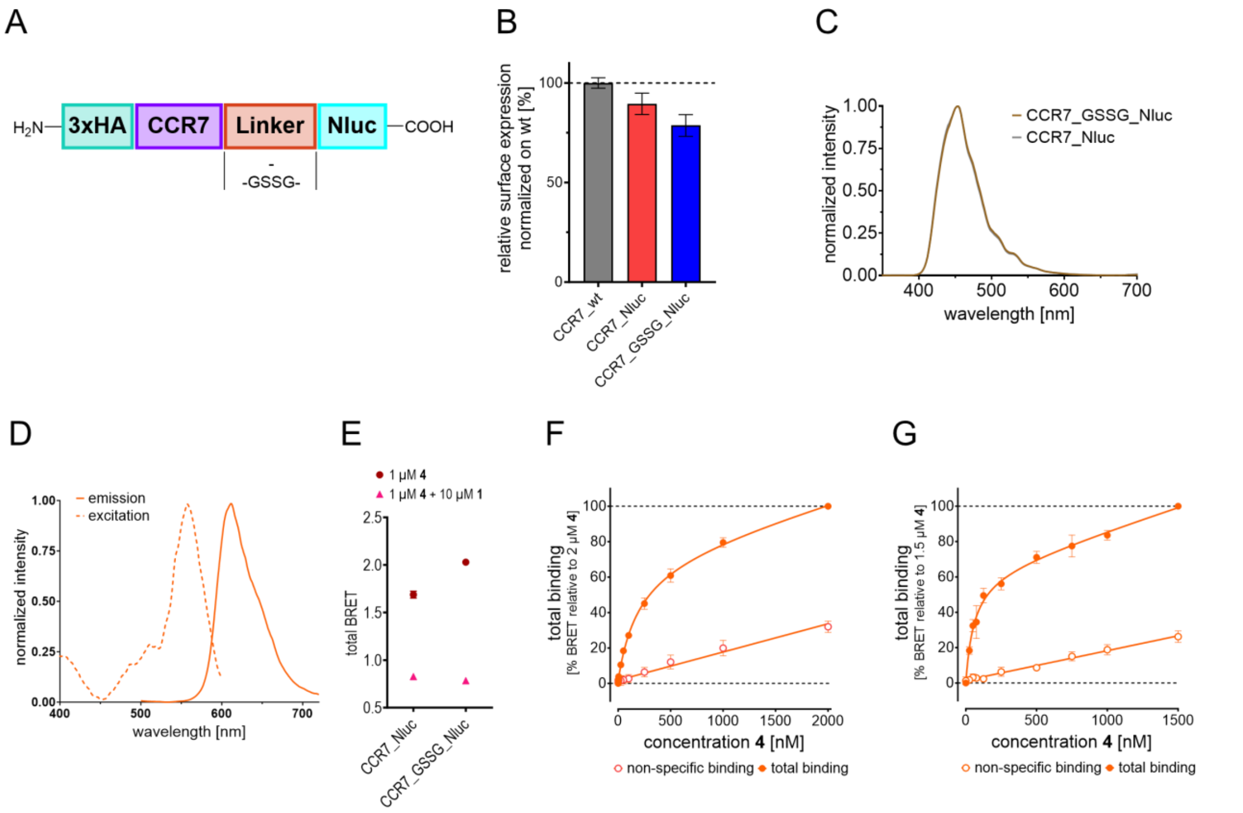


**Figure S1.** Development of a cell-free and cellular NanoBRET-based binding assay for the IABS of CCR7. A) Schematic representation of the genetic C-terminally Nluc-labeled constructs of CCR7 that were investigated in the course of assay development. B) Expression level of different 3xHA-CCR7-Nluc constructs detected via ELISA and normalized to the expression of wild-type 3xHA-CCR7 (CCR7_wt). Bar diagram represents the mean values ± SEM ($n \geq 3$) with each test performed in quadruplicate. The experiment indicates that all CCR7-Nluc constructs are well-expressed in HEK293T cells. C) Emission spectra of membrane preparations from HEK293T cells transiently overexpressing the CCR7_Nluc and CCR7_GSSG_Nluc construct, respectively. D) Spectral properties of fluorescent ligand Mz437 (**4**). Absorption and fluorescence emission spectra were normalized to the respective maximum signal of each sample. E) Representative NanoBRET assay windows for Mz437 (**4**), generated with membrane preparations from HEK293T cells expressing different CCR7-Nluc constructs. Tests were performed in duplicate. The signals for non-specific binding (full displacement of **4**, red triangles) were detected in the presence of **4** (1 µM) and **1** (10 µM), whereas the signals obtained for the vehicle controls represent total binding of **4** at a concentration of 1 µM (blue circles). As the application of CCR7_GSSG_Nluc resulted in a slightly larger assay window, we used this construct for further investigations. F) Binding curves for total and non-specific binding (triplicate measurement, $n = 3$) of **4** to CCR7_GSSG_Nluc membranes. G) Binding curves for total and non-specific binding (quadruplicate measurement, $n = 4$) of **4** to live HEK293T cells transiently overexpressing CCR7_GSSG_Nluc.

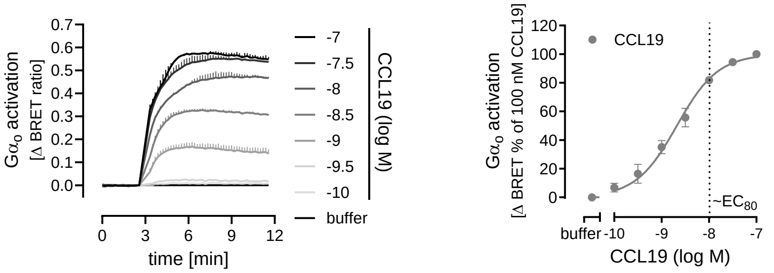


**Figure S2.** Concentration-dependent Go activation of CCR7 by CCL19. Representative kinetic recordings of BRET changes between masGRK3ct-Nluc and G$\alpha_o$-derived G$\beta\gamma$ (Venus-labeled) in transiently transfected HEK293T cells after treatment with increasing concentrations of CCL19. Summarized concentration-effect curve is shown as means ± SEM of n= 6 independent experiments. CCR7 binders were tested against 10 nM CCL19 (~$EC_{80}$ as indicated in the curve).

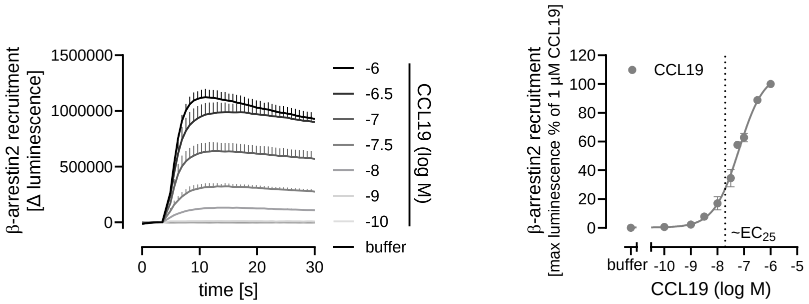


**Figure S3.** CCL19-induced β-arrestin2 recruitment to CCR7. Representative traces of NanoBiT complementation between CCR7-SmBiT and LgBiT-β-arrestin2 mutant (R393E, R395E) in transiently transfected HEK293T cells after stimulation with CCL19. Averaged concentration-effect curve derived from the traces shows means ± SEM of n= 2-5 independent experiments. CCR7 binders were tested against 18 nM CCL19 (~$EC_{25}$ as indicated in the curve).

**Table S1.** Kinetic parameters detected for the interaction between Mz437 (4) and CCR7_GSSG_Nluc using our membrane-based NanoBRET assay and by applying the indicated conditions. A) Dissociation at room temperature. B) Association at room temperature. For the calculation of $k_{on}$, a pre-determined $k_{off}$ of 0.2234 $min^{-1}$ was used.

A

**Dissociation at room temperature (membranes)**

| | $k_{off}$ [$min^{-1}$] | $t_R$ [min] |
|---|---|---|
| 1000 nM | 0.2414 | 4.14 |
| 500 nM | 0.2036 | 4.91 |
| 250 nM | 0.2253 | 4.44 |
| mean | **0.2234** | **4.50** |
| SEM | 0.0110 | 0.22 |

B

**Association at room temperature (membranes)**

| | $k_{on}$ [$M^{-1}$ $min^{-1}$] |
|---|---|
| 1000 nM | 5003923 |
| | 4576441 |
| 500 nM | 3961621 |
| | 5455546 |
| | 3868786 |
| 250 nM | 3210495 |
| | 3018297 |
| 100 nM | 4501295 |
| | 4327296 |
| | 4852031 |
| mean | **4277573** |
| SEM | 244459 |

**Table S2.** Selectivity binding studies with **10** (SLW131) and **21m** (SLW132). CCR9 was selected because it is the most closely related chemokine receptor to CCR7.[1] CCR2 was selected as a second example of a chemokine receptor with a known intracellular allosteric binding pocket.[2] Cmp2105 was used for comparison. NanoBRET-based binding studies were performed as previously described.[3,4] Tests were performed in triplicate (n = 3). $K_i$ values are given in brackets.

| Compound | **CCR2**<br>**p*K*i** ± SEM (*K*i [nM]) or comp. | **CCR9**<br>**p*K*i** ± SEM (*K*i [nM]) or comp. |
|---|---|---|
| **10** | 13% comp. @ 20 µM | **5.83** ± 0.05 (1479) |
| **21m** | 30% comp. @ 20 µM | **5.82** ± 0.02 (1514) |
| **Cmp2105 (1)** | 3% comp. @ 10 µM | **6.28** ± 0.06 (530) |

**Table S3.** Overview of the synthesized squaramide-based navarixin analogues and their respective $pK_i$ values (mean ± SEM, triplicate measurement, n ≥ 3) obtained from our cell-free NanoBRET competition binding assay. Data was obtained with **4** (500 nM) and CCR7_GSSG_Nluc membranes. $K_i$ values are given in brackets.

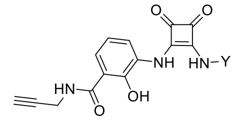


| Compound | Y | $\mathbf{p}\boldsymbol{K}_\mathbf{i}$ ± SEM ($K_i$ [nM]) or comp. |
|---|---|---|
| **21a** | 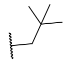 | 9% comp. @10µM |
| **21b** | 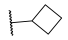 | 1% comp. @10µM |
| **21c** | 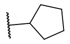 | 6% comp. @10µM |
| **21d** | 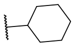 | 15% comp. @10µM |
| **21e** | 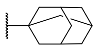 | 23% comp. @10µM |
| **21f** | 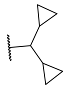 | 13% comp. @10µM |
| **21g** | 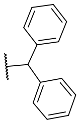 | 5% comp. @10µM |
| **21h** | 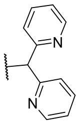 | 13% comp. @10µM |
| **21i** | 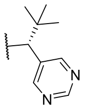 | 5% comp. @10µM |
| **21j** | 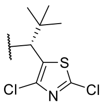 | 18% comp. @10µM |
| **21k** | 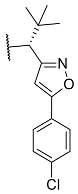 | 24% comp. @10µM |
| **Navarixin (2)** | | 5.13 ± 0.08 (7790) |

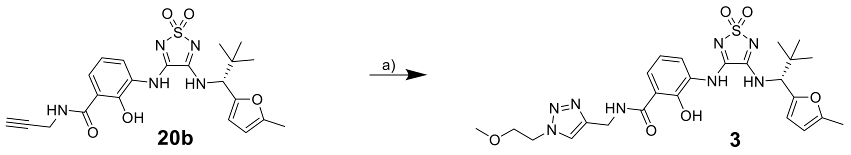


**Scheme S1.** Synthesis of the triazole-based CCR7 ligand-linker conjugate **3**. *Reagents and conditions*: a) 1-azido-2-methoxyethane, $CuSO_4 \cdot 5\,H_2O$, sodium ascorbate, TBTA, water/*tert*-BuOH/DMF mixture (1:1:1 (*v/v/v*)), rt, 2 days, 36%.

EXPERIMENTAL SECTION

#### Chemistry

*General remarks.* Chemicals were obtained from abcr GmbH (Karlsruhe, Germany), Acros Organics (Geel, Belgien), Carbolution Chemicals (Sankt Ingberg, Germany), Sigma-Aldrich (Steinheim, Germany,) TCI Chemicals (Eschborn, Germany) or VWR (Langenfeld, Germany) and used without further purification. Cmp2105 (**1**) was obtained from Hölzel Diagnostika (Köln, Germany). Technical-grade solvents were distilled prior to use. For all HPLC purposes, acetonitrile in HPLC-grade quality (HiPerSolv CHROMANORM, VWR, Langenfeld, Germany) was used. Water was purified with a PURELAB flex® (ELGA VEOLIA, Celle, Germany). Thin-layer chromatography (TLC) was carried out on prefabricated plates (silica gel 60, F254, Merck). Components were visualized either by irradiation with ultraviolet light (254 nm or 366 nm) or by appropriate staining. Column chromatography was carried out on silica gel (60 Å, 40–60 µm, Acros Organics, Geel, Belgium). If no solvent is stated, an aqueous solution was prepared with demineralized water. Mixtures of two or more solvents are specified as "solvent A"/"solvent B", 3/1, *v*/*v*; meaning that 100 mL of the respective mixture consists of 75 mL of "solvent A" and 25 mL of "solvent B". The uncorrected melting points were determined using a Büchi (Essen, Germany) Melting Point M-560 apparatus. Diastereomeric ratios were determined by $^{1}H$ NMR spectroscopy. Proton ($^{1}H$) and carbon ($^{13}C$) NMR spectra were recorded either on a Bruker AVANCE 500 MHz at a frequency of 500 MHz ($^{1}H$) and 126 MHz ($^{13}C$) or a Bruker AVANCE III HD 600 MHz at a frequency of 600 MHz ($^{1}H$) and 151 MHz ($^{13}C$). The chemical shifts are given in parts per million (ppm). As solvents, deuterated chloroform ($CDCl_3$), deuterated methanol (methanol-$d_4$) and deuterated dimethyl sulfoxide (DMSO-$d_6$) were used. The residual solvent signal ($CDCl_3$: $^{1}H$ NMR: 7.26 ppm, $^{13}C$ NMR: 77.1 ppm; DMSO-$d_6$: $^{1}H$ NMR: 2.50 ppm, $^{13}C$ NMR: 39.52 ppm; methanol-$d_4$: $^{1}H$ NMR: 3.31 ppm, 4.87 ppm, $^{13}C$ NMR: 49.00 ppm) was used for calibration. The multiplicity of each signal is reported as singlet (s), doublet (d), triplet (t), quartet (q), multiplet (m) or combinations thereof. Multiplicities and coupling constants are reported as measured and might disagree with the expected values. High-resolution electrospray-ionization mass spectra (HRMS-ESI) were acquired with a Bruker Daltonik GmbH micrOTOF coupled to a an LC Packings Ultimate HPLC system and controlled by micrOTOFControl3.4 and HyStar 3.2-LC/MS, with a Bruker Daltonik GmbH ESI-qTOF Impact II coupled to a Dionex UltiMateTM 3000 UHPLC system and controlled by micrOTOFControl 4.0 and HyStar 3.2-LC/MS or with a micrOTOF-Q mass spectrometer (Bruker, Bremen, Germany) with ESI-source coupled with an HPLC Dionex UltiMate 3000 (Thermo Scientific, Heysham, United Kingdom). Low-resolution electrospray-ionization mass spectra (LRMS-ESI) were acquired with an Advion expression® compact mass spectrometer (CMS) coupled with an automated TLC plate reader Plate Express® (Advion, Ithaca, NY, USA). A Thermo Fisher Scientific (Heysham, United Kingdom) UltiMateTM 3000 UHPLC system with a Nucleodur 100-5 C18 (250 × 4.6 mm, Macherey Nagel, Düren, Germany) with a flow rate of 1 mL/min and a temperature of 25 °C, or a 100-5 C18 (100 × 3 mm, Macherey Nagel, Düren, Germany) with a flow rate of 0.5 mL/min and a temperature of 25 °C was used with an appropriate gradient. For preparative purposes, an AZURA Prep. 500/1000 gradient system with a Nucleodur 110-5 C18 HTec (150 × 32 mm, Macherey Nagel, Düren, Germany) column with 20 mL/min was used. Detection was implemented with UV absorption measurement at wavelengths of λ = 220 nm and λ = 250 nm. Bidest. $H_2O$ (A) and MeCN (B) were used as eluents with an addition of 0.1% TFA in case of eluent A. Purity: The purity of all final compounds was 95% or higher. Purity analysis for compound **4** was performed using an Agilent

1200 series HPLC system employing a diode array detector (DAD, detection at 200, 220, 254 or 560 nm) and a ZORBAX ECLIPSE, XDB-C8 column (4.6 mm x 150 mm, 5 µm) with a flow rate of 0.5 mL·min$^{-1}$. The indicated purity was determined at a wavelength of 254 nm. As solvent systems the following binary solvent systems were used. Elution was performed at room temperature under gradient conditions. Eluent A was water containing 0.1% (v/v) TFA; eluent B was acetonitrile. Linear gradient conditions were as follows: 0–3.0 min: A=90%, B=10%; 3.0-18.0 min: linear increase to A=5%, B=95%; 18.0-24.0 min: A=5%, B=95%; 24.0–27.0 min: linear decrease to A=90%, B=10%; 27.0-30.0 min: A=90%, B=10%. Purity analyses for compounds **3**, **10**, **20a-d**, **21a-m**, were performed on a Thermo Fisher Scientific UltiMateTM 3000 UHPLC system with a Nucleodur 100-5 C18 (250 × 4.6 mm, Macherey Nagel, Düren, Germany) at 250 nm. Elution was performed at a flow rate of 1 mL/min and a temperature of 25 °C. After column equilibration for 5 min, a linear gradient from 5% A to 95% B in 7 min followed by an isocratic regime of 95% B for 10 min was used.

**Synthesis and compound characterization**

Detailed information on experimental procedures for compound synthesis as well as compound characterization data are listed below. NMR spectra and HPLC chromatograms are given in the Supporting Information. Compounds **11c**, **13a**, **14a-c**, **15a-c**, **16a,b**, **17a,b**, **19a,b**, and 3,4-dimethoxy-1,2,5-thiadiazole-1,1-dioxide were synthesized according to previously published procedures.[5–9]

**(*R*)-3-[(4-{[2,2-dimethyl-1-(5-methylfuran-2-yl)propyl]amino}-1,1-dioxido-1,2,5-thiadiazol-3-yl)amino]-2-hydroxy-*N*-{[1-(2-methoxyethyl)-1H-1,2,3-triazol-4-yl]methyl}benzamide** (**3**)

(*R*)-3-[(4-{[2,2-Dimethyl-1-(5-methylfuran-2-yl)propyl]amino}-1,1-dioxido-1,2,5-thiadiazol-3-yl)amino]-2-hydroxy-*N*-(prop-2-yn-1-yl)benzamide (**20b,** 10 mg, 0.097 mmol, 1.0 *eq.*) and 1-azido-2-methoxyethane (48 mg, 0.097 mmol, 1.0 *eq.*) were dissolved in a water/*tert*-BuOH mixture (2 mL, 1:1). Tris(benzyltriazolylmethyl)amine (TBTA, 5.1 mg, 0.010 mmol, 0.10 *eq.*) was dissolved in DMF (1 mL) and added to the mixture. An aqueous $CuSO_4$ solution (0.1 M, 0.48 mL, 0.048 mmol, 0.50 *eq.*) and an aqueous solution of sodium ascorbate (0.1 M, 0.24 mL, 0.024 mmol, 0.25 *eq.*) were added in that order. The resulting reaction mixture was stirred for 2 h at room temperature. After completion of the reaction, volatiles were removed under reduced pressure. The residue was purified by preparative HPLC (acetonitrile/water (0.1%TFA): gradient 5−95%) to obtain the title compound as a colorless solid (20 mg, 36%). $^{1}$H NMR (500 MHz, DMSO-$d_6$, δ [ppm]): 13.72 (s, 1H), 10.59 (s, 1H), 9.62 (t, $J$ = 5.8 Hz, 1H), 9.25 (d, $J$ = 9.1 Hz, 1H), 8.00 (s, 1H), 7.87 (d, $J$ = 7.9 Hz, 1H), 7.01 (t, $J$ = 8.0 Hz, 1H), 6.32 (d, $J$ = 3.1 Hz, 1H), 6.07 (dd, $J$ = 3.1, 1.2 Hz, 1H), 4.87 (d, $J$ = 9.0 Hz, 1H), 4.59 (d, $J$ = 5.8 Hz, 2H), 4.50 (t, $J$ = 5.2 Hz, 2H), 3.72 (t, $J$ = 5.2 Hz, 2H), 3.23 (s, 3H), 2.30 (s, 2H), 1.01 (s, 9H); $^{13}$C NMR (126 MHz, DMSO-$d_6$, δ [ppm]): 169.3, 155.8, 154.0, 153.4, 151.3, 149.3, 143.8, 129.6, 125.5, 125.2, 123.4, 118.1, 115.0, 109.6, 106.4, 70.1, 61.8, 57.9, 49.1, 35.7, 34.7, 26.5, 13.4; HRMS m/z (ESI$^+$) [found: 573.2238, $C_{25}H_{33}N_8O_6S^+$ requires [M + H]$^+$ 573.2166]; HPLC retention time: 12.45 min, purity: 95.1%.

**(*R*)-4-((2-(2-(2-(2-(4-((3-((4-((2,2-Dimethyl-1-(5-methylfuran-2-yl)propyl)amino)-1,1-dioxido-1,2,5-thiadiazol-3-yl)amino)-2-hydroxybenzamido)methyl)-1*H*-1,2,3-triazol-1-**

**yl)ethoxy)ethoxy)ethoxy)ethyl)carbamoyl)-2-(6-(dimethylamino)-3-(dimethyliminio)-3*H*-xanthen-9-yl)benzoate (Mz437, 4)**

(*R*)-3-((4-((2,2-Dimethyl-1-(5-methylfuran-2-yl)propyl)amino)-1,1-dioxido-1,2,5-thiadiazol-3-yl)amino)-2-hydroxy-*N*-(prop-2-yn-1-yl)benzamide (**20b**, 4.0 mg, 8.5 µmol, 1.0 *eq.*), the 6-TAMRA-PEG3-azide (5.4 mg, 8.5 µmol, 1.0 *eq.*), and tris(benzyltriazolylmethyl)amine (TBTA, 0.50 mg, 0.85 µmol, 0.10 *eq.*) were dissolved in a water/*tert*-BuOH/DMF mixture (1.5 mL, 1:1:1 (v/v/v)). An aqueous $CuSO_4$ solution (8.5 µL, 0.10 M, 0.10 *eq.*) and an aqueous solution of sodium ascorbate (17 µL, 0.10 M, 0.20 *eq.*) were added in that order. The resulting reaction mixture was stirred for 6 h at room temperature under nitrogen atmosphere. After completion, volatiles were removed under reduced pressure. The residue was purified by preparative HPLC (acetonitrile/water (0.1% TFA): gradient 30 % - 55 %) to obtain the TFA salt of title compound as a pink solid (5.4 mg, 52%). $^1$H NMR (600 MHz, DMSO-$d_6$, δ [ppm]): 13.71 (s, 1H), 13.36 (s, 1H), 10.60 (s, 1H), 9.61 (t, *J* = 5.7 Hz, 1H), 9.26 (d, *J* = 9.0 Hz, 1H), 8.79 (t, *J* = 5.5 Hz, 1H), 8.32 - 8.25 (m, 1H), 8.22 (d, *J* = 8.2 Hz, 1H), 7.99 (s, 1H), 7.89 - 7.81 (m, 3H), 7.18 - 6.79 (m, 6H), 6.98 (t, *J* = 8.0 Hz, 1H), 6.31 (d, *J* = 3.1 Hz, 1H), 6.07 (dq, *J* = 3.1 Hz, 1.1 Hz, 1H), 4.86 (d, *J* = 9.0 Hz, 1H), 4.54 (d, *J* = 5.7 Hz, 2H), 4.46 (t, *J* = 5.3 Hz, 2H), 3.76 (t, *J* = 5.3 Hz, 2H), 3.50$^{\#}$ (t, *J* = 5.9 Hz, 2H), 3.47 - 3.40$^{\#}$ (m, 10H), 3.24 (s, 12H), 2.29 (d, *J* = 1.1 Hz, 3H), 1.00 (s, 9H); $^{13}$C NMR$^a$ (DEPTQ, 151 MHz, DMSO-$d_6$, δ [ppm]): 169.3, 164.5, 157.6 (q, *J* = 32.6 Hz), 155.8, 154.0, 153.4, 151.3, 149.3, 143.8, 133.0*, 130.7*, 130.6, 129.6, 129.0, 128.6*, 125.5, 125.2, 123.5, 118.1, 114.4*, 114.9, 109.7, 106.4, 96.3, 69.6, 69.6, 69.5, 69.5, 68.7, 68.6, 61.8, 49.3, 40.4, 39.4*$^{\#}$, 35.7, 34.6, 26.6, 13.4; $^a$the $^{13}$C NMR (DEPTQ) signals for 3*H*-xanthene C-3,4a,6,8a,9,9a,10a, carbamoyl benzoate C-2,4, -COOH, and $F_3\underline{C}$-$COO^-$ could not be detected. *Could only be detected in HSQC/HMBC spectra; $^{\#}$partially overlaid by solvent signal. HRMS *m/z* (ESI$^+$) [found: 1102.4455, $C_{55}H_{64}N_{11}O_{12}S^+$ requires [M+H]$^+$ 1102.4451]; HPLC retention time: 17.63 min, 95.3%).

**(*R*)-3-[(4-{[2,2-Dimethyl-1-(5-methylfuran-2-yl)propyl]amino}-1,1-dioxido-1,2,5-thiadiazol-3-yl)amino]-2-hydroxy-*N*,*N*-dimethylbenzamide (SLW131, 10)**

2-Hydroxy-3-[(4-methoxy-1,1-dioxido-1,2,5-thiadiazol-3-yl)amino]-*N*,*N*-dimethylbenzamide (**18a**, 50 mg, 0.20 mmol, 1.0 *eq.*) and (*R*)-2,2-dimethyl-1-(5-methylfuran-2-yl)propan-1-amine hydrochloride (**15c**, 31 mg, 0.20 mmol, 1.0 *eq.*) were dissolved in methanol (10 mL). After the addition of *N*,*N*-diisopropylethylamine (54 µL, 0.40 mmol, 2.0 *eq.*), the reaction was stirred for 3 days. Extraction between water and ethyl acetate (3 × 50 mL), drying over sodium sulfate, filtration, and evaporation of the solvent gave the crude product which was purified by preparative HPLC (acetonitrile/water (0.1% TFA): gradient 5−95%) to obtain the title compound as a white-brown amorphous solid (17 mg, 24%). $^1$H NMR (500 MHz, DMSO-$d_6$, δ [ppm]): 10.48 (s, 1H), 9.88 (s, 1H), 8.89 (d, *J* = 9.1 Hz, 1H), 7.46 (dd, *J* = 7.9, 1.7 Hz, 1H), 7.12 (dd, *J* = 7.6, 1.7 Hz, 1H), 6.96 (t, *J* = 7.8 Hz, 1H), 6.32 (d, *J* = 3.1 Hz, 1H), 6.07 (dd, *J* = 3.2, 1.3 Hz, 1H), 4.85 (d, *J* = 9.1 Hz, 1H), 2.92 (s, 6H), 2.29 (s, 3H), 1.02 (s, 9H); $^{13}$C NMR (126 MHz, DMSO-$d_6$, δ [ppm]): 167.7, 156.0, 154.6, 151.2, 149.4, 147.3, 127.3, 126.8, 126.5, 124.5, 119.7, 109.5, 106.3, 61.6, 35.7, 26.5, 23.9, 13.4; HRMS m/z (ESI$^+$) [found: 462.1806, $C_{21}H_{28}N_5O_5S^+$ requires [M + H]$^+$ 462.1766]; HPLC retention time: 12.33 min, purity: 95.6%.

**(*S*,*E*)-2-Methyl-*N*-[(5-methylthiophen-2-yl)methylene]propane-2-sulfinamide (13b)**

(*S*)-2-Methylpropane-2-sulfinamide (1.00 g, 8.09 mmol, 1.00 *eq.*) was dissolved in dichloromethane (10 mL). 5-Methylthiophene-2-carboxaldehyde (918 µL, 8.09 mmol, 1.00 *eq.*), titanium ethoxide (3.73 mL, 17.8 mmol, 2.20 *eq*), and sodium sulfate (2 g) were added under stirring. The reaction mixture was stirred at room temperature overnight, filtered through celite, and rinsed with dichloromethane. Evaporation of the solvent gave the crude product in quantitative yield, which was used without further purification.

**(*S*,*E*)-2-Methyl-*N*-(pyrimidin-5-ylmethylene)propane-2-sulfinamide (13c)**
(*S*)-2-Methylpropane-2-sulfinamide (1.00 g, 8.09 mmol, 1.00 *eq.*) was dissolved in dichloromethane (10 mL). Pyrimidine-5-carboxaldehyde (920 mg., 8.09 mmol, 1.00 *eq.*), titanium ethoxide (3.73 mL, 17.8 mmol, 2.20 *eq*), and sodium sulfate (2 g) were added under stirring. The reaction mixture was stirred at room temperature overnight, filtered through celite, and rinsed with dichloromethane. Evaporation of the solvent gave the crude product in quantitative yield, which was used without further purification.

**(*S*,*E*)-*N*-[(2,4-Dichlorothiazol-5-yl)methylene]-2-methylpropane-2-sulfinamide (13d)**
(*S*)-2-Methylpropane-2-sulfinamide (650 mg, 5.26 mmol, 1.00 *eq.*) was dissolved in dichloromethane (10 mL). 2,4-Dichlorothiazole-5-carboxaldehyde (1.01 g, 5.26 mmol, 1.00 *eq.*), titanium ethoxide (2.42 mL, 11.6 mmol, 2.20 *eq.*), and sodium sulfate (2 g) were added under stirring. The reaction mixture was stirred at room temperature overnight, filtered through celite, and rinsed with dichloromethane. Evaporation of the solvent gave the crude product in quantitative yield, which was used without further purification.

**(*S*,*E*)-*N*-{[5-(4-Chlorophenyl)isoxazol-3-yl]methylene}-2-methylpropane-2-sulfinamide (13e)**
(*S*)-2-Methylpropane-2-sulfinamide (600 mg, 4.85 mmol, 1.00 *eq.*) was dissolved in dichloromethane (10 mL). 5-(4-Chlorophenyl)isoxazole-3-carboxaldehyde (1.06 g, 4.85 mmol, 1.00 *eq.*), titanium ethoxide (2.24 mL, 10.7 mmol, 2.20 *eq*), and sodium sulfate (2 g) were added under stirring. The reaction mixture was stirred at room temperature overnight, filtered through celite, and rinsed with dichloromethane. Evaporation of the solvent gave the crude product in quantitative yield, which was used without further purification.

**(*S*)-*N*-[(*R*)-2,2-Dimethyl-1-(5-methylthiophen-2-yl)propyl]-2-methylpropane-2-sulfinamide (14d)**
A 250 mL flask was sealed with a septum flushed with $N_2$ and filled with THF (20 mL). A 1 M *tert*-butylmagnesium chloride solution in THF (18.0 mL, 17.8 mmol, 2.20 *eq.*) was added and the solution was cooled to 0 °C. (*S*,*E*)-2-Methyl-*N*-[(5-methylthiophen-2-yl)methylene]propane-2-sulfinamide (**13b**, 1.86 g, 8.10 mmol, 1.00 *eq.*) was dissolved in THF (20 mL) and added dropwise to the vigorously stirred solution. After addition, the mixture was stirred for 72 h. The reaction mixture was then quenched by the addition of saturated ammonium chloride solution (50 mL) and extracted using ethyl acetate (3 x 100 mL). Drying over sodium sulfate, filtration, and evaporation of the solvent resulted in the crude product which was purified by column chromatography using a gradient of dichloromethane and ethyl acetate (dichloromethane to dichloromethane/ethyl acetate (8/2) to dichloromethane/ethyl acetate (5/5) (*v/v*)). The title compound was obtained as a dark orange to red oil (264 mg, 11%). $^1$H NMR (500 MHz, DMSO-$d_6$ δ [ppm]) 6.77 (d, $J$ = 3.4 Hz, 1H), 6.63 (dq, $J$ = 3.4, 1.2 Hz, 1H), 4.40 (d, $J$ = 4.4 Hz, 1H), 4.18 (d, $J$ = 4.4 Hz, 1H), 2.39 (d, $J$ = 1.1 Hz, 3H), 1.08 (s, 9H), 0.94 (s, 9H); $^{13}$C NMR (126 MHz, DMSO-$d_6$ δ [ppm]) 140.9, 138.1, 126.7, 124.2, 64.3, 55.2, 35.1, 26.5, 22.3, 14.9; LRMS *m/z* ($ESI^+$) [Found: 288.5, $C_{14}H_{26}NOS_2^+$

requires $[M+H]^+$ 288.5]; $R_f$: 0.34 (cyclohexane/ethyl acetate (1/1) (*v/v*)); Diastereomeric ratio: *dr* = 1:0.

**(*S*)-*N*-[(*R*)-2,2-Dimethyl-1-(pyrimidin-5-yl)propyl]-2-methylpropane-2-sulfinamide (14e)**
A 250 mL flask was sealed with a septum flushed with $N_2$ and filled with THF (20 mL). A 1 M *tert*-butylmagnesium chloride solution in THF (18.0 mL, 17.8 mmol, 2.20 *eq.*) was added and the solution was cooled to 0 °C. (*S,E*)-2-methyl-*N*-(pyrimidin-5-ylmethylene)propane-2-sulfinamide (**13c**, 1.71 g, 8.10 mmol, 1.00 *eq.*) was dissolved in THF (20 mL) and added dropwise to the vigorously stirred solution. After addition, the mixture was stirred for 72 h. The reaction mixture was then quenched by the addition of saturated ammonium chloride solution (50 mL) and extracted using ethyl acetate (3 x 100 mL). Drying over sodium sulfate, filtration, and evaporation of the solvent resulted in the crude product which was purified by column chromatography using a gradient of dichloromethane and ethyl acetate (dichloromethane to dichloromethane/ethyl acetate (8/2) to dichloromethane/ethyl acetate (5/5) (*v/v*)). The title compound was obtained as a dark orange to red oil (115 mg, 5%). $^1$H NMR (500 MHz, DMSO-$d_6$ δ [ppm]) 9.06 (s, 1H), 8.77 (s, 2H), 5.17 (d, *J* = 7.5 Hz, 1H), 4.16 (d, *J* = 7.6 Hz, 1H), 1.08 (s, 9H), 0.93 (s, 9H); $^{13}$C NMR (126 MHz, DMSO-$d_6$ δ [ppm]) 157.0, 156.6, 134.4, 64.9, 55.5, 53.9, 35.2, 26.3, 22.2; LRMS *m/z* ($ESI^+$) [Found: 288.5, $C_{13}H_{24}N_3OS^+$ requires $[M+H]^+$ 270.4]; $R_f$: 0.20 (cyclohexane/ethyl acetate (1/1) (*v/v*)); Diastereomeric ratio: *dr* = 1:0.

**(*S*)-*N*-[(*R*)-1-(2,4-Dichlorothiazol-5-yl)-2,2-dimethylpropyl]-2-methylpropane-2-sulfinamide (14f)**
A 250 mL flask was sealed with a septum flushed with $N_2$ and filled with THF (20 mL). A 1 M *tert*-butylmagnesium chloride solution in THF (12.0 mL, 11.6 mmol, 2.20 *eq.*) was added and the solution was cooled to 0 °C. (*S,E*)-*N*-[(2,4-dichlorothiazol-5-yl)methylene]-2-methylpropane-2-sulfinamide (**13d**, 1.5 g, 5.3 mmol, 1. *eq.*) was dissolved in THF (20 mL) and added dropwise to the vigorously stirred solution. After addition, the mixture was stirred for 72 h. The reaction mixture was then quenched by the addition of saturated ammonium chloride solution (50 mL) and extracted using ethyl acetate (3 x 100 mL). Drying over sodium sulfate, filtration, and evaporation of the solvent resulted in the crude product which was purified by column chromatography using a gradient of dichloromethane and ethyl acetate (dichloromethane to dichloromethane/ethyl acetate (8/2) to dichloromethane/ethyl acetate (5/5) (*v/v*)). The title compound was obtained as a dark orange to red oil (447 mg, 25%). $^1$H NMR (500 MHz, DMSO-$d_6$ δ [ppm]) 4.97 (d, *J* = 5.2 Hz, 1H), 4.40 (d, *J* = 5.2 Hz, 1H), 1.08 (s, 9H), 1.00 (s, 9H); $^{13}$C NMR (126 MHz, DMSO-$d_6$ δ [ppm]) 150.2, 134.9, 134.1, 61.9, 55.8, 36.2, 26.1, 22.2; LRMS *m/z* ($ESI^+$) [Found: 345.5, $C_{12}H_{21}Cl_2NOS_2^+$ requires $[M+H]^+$ 345.3.]; $R_f$: 0.51 (cyclohexane/ethyl acetate (1/1) (*v/v*)); Diastereomeric ratio: *dr* = 1:0.

**(*S*)-*N*-(*R*)-1-[5-(4-Chlorophenyl)isoxazol-3-yl)-2,2-dimethylpropyl]-2-methylpropane-2-sulfinamide (14g)**
A 250 mL flask was sealed with a septum flushed with $N_2$ and filled with THF (20 mL). A 1 M *tert*-butylmagnesium chloride solution in THF (11.2 mL, 11.2 mmol, 2.20 *eq.*) was added and the solution was cooled to 0 °C. (*S,E*)-*N*-{[5-(4-Chlorophenyl)isoxazol-3-yl]methylene}-2-methylpropane-2-sulfinamide (**13e**, 1.6 g, 5.1 mmol, 1.0 *eq.*) was dissolved in THF (20 mL) and added dropwise to the vigorously stirred solution. After addition, the mixture was stirred for 72 h. The reaction mixture was then quenched by the addition of saturated ammonium chloride solution

(50 mL) and extracted using ethyl acetate (3 x 100 mL). Drying over sodium sulfate, filtration, and evaporation of the solvent resulted in the crude product which was purified by column chromatography using a gradient of dichloromethane and ethyl acetate (dichloromethane to dichloromethane/ethyl acetate (8/2) to dichloromethane/ethyl acetate (5/5) (*v/v*)). The title compound was obtained as a dark orange to red oil (447 mg, 25%). $^{1}$H NMR (500 MHz, DMSO-$d_6$ δ [ppm]) 7.89 - 7.80 (m, 2H), 7.66 - 7.57 (m, 2H), 7.14 (s, 1H), 5.15 (d, *J* = 10.1 Hz, 1H), 4.10 (d, *J* = 10.1 Hz, 1H), 1.16 (s, 9H), 0.95 (s, 9H); $^{13}$C NMR (126 MHz, DMSO-$d_6$ δ [ppm]) 166.9, 165.0, 134.9, 129.4, 127.2, 125.7, 100.9, 61.8, 56.2, 35.3, 26.5, 22.7; LRMS *m/z* (ESI$^+$) [Found: 369.5, $C_{18}H_{26}ClNO_2S^+$ requires [M+H]$^+$ 369.9]; $R_f$: 0.51 (cyclohexane/ethyl acetate (1/1) (*v/v*)); Diastereomeric ratio: *dr* = 1:0.

**(*R*)-2,2-Dimethyl-1-(5-methylthiophen-2-yl)propan-1-amine hydrochloride (15d)**
(*S*)-*N*-[(*R*)-2,2-Dimethyl-1-(5-methylthiophen-2-yl)propyl]-2-methylpropane-2-sulfinamide (200 mg, 0.700 mmol, 1.00 *eq.*) was dissolved in diethyl ether (10 mL) and cooled to 0 °C. After addition of 2 M HCl in diethyl ether (873 µL, 1.75 mmol, 2.50 *eq.*) the mixture was stirred for 1 h. After evaporation of the solvents, **15d** was obtained as a reddish oil that was used in the next step without further purification. LRMS *m/z* (ESI$^+$) [Found: 185.3, $C_{10}H_{19}NS^+$ requires [M+H]$^+$ 185.3].

**(*R*)-2,2-Dimethyl-1-(pyrimidin-5-yl)propan-1-amine hydrochloride (15e)**
(*S*)-*N*-[(*R*)-2,2-Dimethyl-1-(pyrimidin-5-yl)propyl]-2-methylpropane-2-sulfinamide (80 mg, 0.28 mmol, 1.0 *eq.*) was dissolved in diethyl ether (10 mL) and cooled to 0 °C. After addition of 2 M HCl in diethyl ether (1.6 mL, 0.70 mmol, 2.5 *eq.*) the mixture was stirred for 1 h. After evaporation of the solvents, **15e** was obtained as a reddish oil that was used in the next step without further purification. LRMS *m/z* (ESI$^+$) [Found: 166.4, $C_9H_{17}N_3^+$ requires [M+H]$^+$ 166.2].

**(*R*)-1-(2,4-Dichlorothiazol-5-yl)-2,2-dimethylpropan-1-amine hydrochloride (15f)**
(*S*)-*N*-[(*R*)-1-(2,4-Dichlorothiazol-5-yl)-2,2-dimethylpropyl]-2-methylpropane-2-sulfinamide (300 mg, 0.872 mmol, 1.00 *eq.*) was dissolved in diethyl ether (10 mL) and cooled to 0 °C. After addition of 2 M HCl in diethyl ether (1.10 mL, 2.18 mmol, 2.50 *eq.*) the mixture was stirred for 1 h. After evaporation of the solvents, **15f** was obtained as a reddish oil that was used in the next step without further purification. LRMS *m/z* (ESI$^+$) [Found: 240.4, $C_8H_{14}Cl_2N_2S^+$ requires [M+H]$^+$ 240.2].

**(*R*)-1-[5-(4-Chlorophenyl)isoxazol-3-yl]-2,2-dimethylpropan-1-amine hydrochloride (15g)**
(*S*)-*N*-(*R*)-1-[5-(4-Chlorophenyl)isoxazol-3-yl)-2,2-dimethylpropyl]-2-methylpropane-2-sulfinamide (300 mg, 0.872 mmol, 1.00 *eq.*) was dissolved in diethyl ether (10 mL) and cooled to 0 °C. After addition of 2 M HCl in diethyl ether (1.10 mL, 2.18 mmol, 2.50 *eq.*) the mixture was stirred for 1 h. After evaporation of the solvents, **15g** was obtained as a reddish oil that was used in the next step without further purification. LRMS *m/z* (ESI$^+$) [Found: 266.8, $C_{14}H_{19}ClN_2O^+$ requires [M+H]$^+$ 266.8].

**2-Hydroxy-*N*-methyl-3-nitro-*N*-(prop-2-yn-1-yl)benzamide (16c)**
2-Hydroxy-3-nitrobenzoic acid (1.00 g, 5.35 mmol, 1.00 *eq.*), bromotrispyrrolidinophosphonium hexafluorophosphate (3.02 g, 6.42 mmol, 1.20 *eq.*) and *N,N*-diisopropylethylamine (3.78 mL, 21.4 mmol, 4.00 *eq.*) were dissolved in dichloromethane (30 mL) and stirred at room temperature for 30 min. *N*-Methylpropargylamine (666 µL, 7.49 mmol, 1.40 *eq.*) was added and the resulting

mixture was stirred overnight. The reaction was then extracted with 1 N sodium hydroxide solution (3 x 100 mL) and the organic phase was discarded. The aqueous phase was then acidified using 1 M hydrochloric acid (300 mL) and the organic products were extracted using ethyl acetate (3 x 150 mL). The organic phase was dried over sodium sulfate, filtered, and concentrated *in vacuo* to afford the crude product, which was then purified by column chromatography using a mixture of cyclohexane and ethyl acetate (6/4 (v/v)). The title compound was obtained as a pale yellow oil (1.2 g, 95%). $^{1}$H NMR (500 MHz, DMSO-$d_6$ δ [ppm]): 10.71 (s, 1H), 8.05 (dd, *J* = 8.3, 1.7 Hz, 1H), 7.56 (dd, *J* = 7.5, 1.7 Hz, 1H), 7.11 (dd, *J* = 8.4, 7.4 Hz, 1H), 4.32 (s, 1H), 3.39 - 3.21 (m, 2H), 3.10 - 2.81 (m, 3H); $^{13}$C NMR (151 MHz, DMSO-$d_6$ δ [ppm]): 166.0, 148.6, 136.2, 134.3, 134.2, 128.4, 125.9, 120.1, 78.9, 74.6, 35.3; LRMS m/z (ESI$^+$) [Found: 235.1, $C_{11}H_{11}N_2O_4^+$ requires [M+H]$^+$ 235.1]; Rf: 0.55 (cyclohexane/ethyl acetate (6/4) (v/v)).

**3-Amino-2-hydroxy-*N*-methyl-*N*-(prop-2-yn-1-yl)benzamide (17c)**

2-Hydroxy-*N*-methyl-3-nitro-*N*-(prop-2-yn-1-yl)benzamide (**16c,** 1.15 g, 4.86 mmol, 1.00 *eq.*) and tin(II) chloride dihydrate (5.48 g, 24.3 mmol, 5.00 *eq.*) were dissolved in methanol (25 mL) and heated under reflux for an hour. After cooling, the mixture was extracted with ethyl acetate (4 x 100 mL). After drying over sodium sulfate, filtration, and evaporation of the solvent, the title compound was obtained as a brown oil in quantitative yield that was used without further purification. LRMS *m/z* (ESI$^+$) [Found: 205.1, $C_{11}H_{13}N_2O_2^+$ requires [M+H]$^+$ 205.1].

**2-Hydroxy-3-[(4-methoxy-1,1-dioxido-1,2,5-thiadiazol-3-yl)amino]-*N*,*N*-dimethylbenzamide (18a)**

3-Amino-2-hydroxy-*N*,*N*-dimethylbenzamide (**17a**, 0.1 g, 0.55 mmol, 1.0 *eq.*) and 3,4-dimethoxy-1,2,5-thiadiazole-1,1-dioxide (99 mg, 0.55 mmol, 1.0 *eq.*) were dissolved in methanol (10 mL) and stirred overnight. Filtration of the solid and drying under reduced pressure afforded the title compound as a green gum (130 mg, 72% yield) that was used without further purification. LRMS *m/z* (ESI$^+$) [Found: 327.5, $C_{12}H_{15}N_4O_5S^+$ requires [M+H]$^+$ 327.3].

**2-Hydroxy-3-[(4-methoxy-1,1-dioxido-1,2,5-thiadiazol-3-yl)amino]-*N*-(prop-2-yn-1-yl)benzamide (18b)**

3-Amino-2-hydroxy-*N*-(prop-2-yn-1-yl)benzamide (**17b**, 250 mg, 1.31 mmol, 1.00 *eq.*) and 3,4-dimethoxy-1,2,5-thiadiazole-1,1-dioxide (234 mg, 1.31 mmol, 1.00 *eq.*) were dissolved in methanol (10 mL) and stirred overnight. Filtration of the solid and drying under reduced pressure afforded the title compound as a green gum (200 mg, 45% yield) that was used without further purification. LRMS *m/z* (ESI$^+$) [Found: 337.3, $C_{13}H_{13}N_4O_5S^+$ requires [M+H]$^+$ 337.3].

**2-Hydroxy-3-[(4-methoxy-1,1-dioxido-1,2,5-thiadiazol-3-yl)amino]-*N*-methyl-*N*-(prop-2-yn-1-yl)benzamide (18c)**

3-Amino-2-hydroxy-*N*-methyl-*N*-(prop-2-yn-1-yl)benzamide (**17c**, 0.10 g, 0.49 mmol, 1.0 *eq.*) and 3,4-dimethoxy-1,2,5-thiadiazole-1,1-dioxide (87 mg, 0.49 mmol, 1.0 *eq.*) were dissolved in methanol (10 mL) and stirred overnight. Filtration of the solid and drying under reduced pressure afforded the title compound as a green gum (125 mg, 73% yield) that was used without further purification. LRMS *m/z* (ESI$^+$) [Found: 351.5, $C_{14}H_{15}N_4O_5S^+$ requires [M+H]$^+$ 351.3].

**(*R*)-3-[(4-{[2,2-Dimethyl-1-(5-methylfuran-2-yl)propyl]amino}-1,1-dioxido-1,2,5-thiadiazol-3-yl)amino]-2-hydroxy-*N*-methyl-N-(prop-2-yn-1-yl)benzamide (20a)**

2-Hydroxy-3-[(4-methoxy-1,1-dioxido-1,2,5-thiadiazol-3-yl)amino]-*N*-methyl-*N*-(prop-2-yn-1-yl)benzamide (**18c**, 45 mg, 0.10 mmol, 1.0 *eq.*) and (*R*)-2,2-dimethyl-1-(5-methylfuran-2-yl)propan-1-amine hydrochloride (**15c**, 31 mg, 0.10 mmol, 1.0 *eq.*) were dissolved in methanol (10 mL). After the addition of *N*,*N*-diisopropylethylamine (46 µL, 0.20 mmol, 2.0 *eq.*), the reaction was stirred for 3 days. Extraction between water and ethyl acetate (3 × 50 mL), drying over sodium sulfate, filtration, and evaporation of the solvent gave the crude product which was purified by preparative HPLC (acetonitrile/water (0.1% TFA): gradient 5−95%) to obtain the title compound as a white-brown amorphous solid (28 mg, 45%). $^1$H NMR (500 MHz, DMSO-$d_6$, δ [ppm]): 10.49 (s, 1H), 9.94 (s, 1H), 8.85 (d, *J* = 9.3 Hz, 1H), 7.46 (dd, *J* = 8.0, 1.7 Hz, 1H), 7.13 (dd, *J* = 7.5, 1.6 Hz, 1H), 6.98 (t, *J* = 7.8 Hz, 1H), 6.32 (s, 1H), 6.07 (dd, *J* = 3.1, 1.2 Hz, 1H), 4.84 (d, *J* = 9.1 Hz, 1H), 4.16 (s, 2H), 3.26 (s, 1H), 2.92 (s, 3H), 2.29 (s, 3H), 1.02 (s, 9H); $^{13}$C NMR (126 MHz, DMSO-$d_6$, δ [ppm]): 167.5, 156.0, 154.8, 151.2, 149.5, 147.5, 127.8, 126.8, 125.8, 124.5, 119.8, 109.5, 106.3, 79.1, 61.6, 40.1, 35.7, 26.5, 13.4; HRMS m/z (ESI$^+$) [found: 486.1766, $C_{23}H_{28}N_5O_5S^+$ requires [M + H]$^+$ 486.1806]; HPLC retention time: 12.09 min, purity: 97.9%.

**(*R*)-3-[(4-{[2,2-Dimethyl-1-(5-methylfuran-2-yl)propyl]amino}-1,1-dioxido-1,2,5-thiadiazol-3-yl)amino]-2-hydroxy-*N*-(prop-2-yn-1-yl)benzamide (20b)**

2-Hydroxy-3-[(4-methoxy-1,1-dioxido-1,2,5-thiadiazol-3-yl)amino]-*N*-(prop-2-yn-1-yl)benzamide (**18b**, 100 mg, 0.300 mmol, 1.00 *eq.*) and (*R*)-2,2-dimethyl-1-(5-methylfuran-2-yl)propan-1-amine hydrochloride (**15c**, 72 mg, 0.36 mmol, 1.2 *eq.*) were dissolved in methanol (10 mL). After the addition of *N*,*N*-diisopropylethylamine (106 µL, 0.600 mmol, 2.00 *eq.*), the reaction was stirred for 3 days. Extraction between water and ethyl acetate (3 × 50 mL), drying over sodium sulfate, filtration, and evaporation of the solvent gave the crude product which was purified by preparative HPLC (acetonitrile/water (0.1% TFA): gradient 5−95%) to obtain the title compound as a white-brown amorphous solid (93 mg, 66%). $^1$H NMR (600 MHz, methanol-$d_4$, δ [ppm]): NH and OH signals were not detectable due to proton exchange, 8.18 (d, *J* = 8.1 Hz, 1H), 7.63 (d, *J* = 8.1 Hz, 1H), 6.97 (t, *J* = 8.1 Hz, 1H), 6.24 (d, *J* = 3.1 Hz, 1H), 6.00 - 5.98 (m, 1H), 4.98 (s, 1H), 4.18 (d, *J* = 2.6 Hz, 2H), 2.63 (t, *J* = 2.5 Hz, 1H), 2.30 (s, 3H), 1.08 (s, 9H); $^{13}$C NMR (151 MHz, methanol-$d_4$, δ [ppm]): 171.2, 157.9, 154.9, 154.5, 153.2, 150.7, 129.3, 127.2, 125.5, 119.3, 115.8, 110.9, 107.2, 80.3, 72.3, 63.9, 37.0, 29.5, 27.2, 13.5; HRMS m/z (ESI$^+$) [found: 494.1469, $C_{22}H_{26}N_5O_5S^+$ requires [M + Na]$^+$ 494.1576]; HPLC retention time: 12.74 min, purity: 99.1%.

**(*R*)-2-Hydroxy-3-[(4-{[2-methyl-1-(5-methylfuran-2-yl)propyl]amino}-1,1-dioxido-1,2,5-thiadiazol-3-yl)amino]-*N*-(prop-2-yn-1-yl)benzamide (20c)**

2-Hydroxy-3-[(4-methoxy-1,1-dioxido-1,2,5-thiadiazol-3-yl)amino]-*N*-(prop-2-yn-1-yl)benzamide (**18b**, 130 mg, 0.370 mmol, 1.00 *eq.*) and (*R*)-2-methyl-1-(5-methylfuran-2-yl)propan-1-amine hydrochloride (**15b**, 57 mg, 0.37 mmol, 1.0 *eq.*) were dissolved in methanol (10 mL). After the addition of *N*,*N*-diisopropylethylamine (132 µL, 0.740 mmol, 2.00 *eq.*), the reaction was stirred for 3 days. Extraction between water and ethyl acetate (3 × 50 mL), drying over sodium sulfate, filtration, and evaporation of the solvent gave the crude product which was purified by preparative HPLC (acetonitrile/water (0.1% TFA): gradient 5−95%) to obtain the title compound as a white-brown amorphous solid (38 mg, 22%). $^1$H NMR (600 MHz, DMSO-$d_6$, δ [ppm]): 13.61 (s, 1H), 10.26 (s, 1H), 9.68 - 9.61 (m, 1H), 9.54 - 9.52 (m, 1H), 8.00 - 7.96 (m, 1H),

7.83 - 7.80 (m, 1H), 7.02 (t, *J* = 8.0 Hz, 1H), 6.37 - 6.32 (m, 1H), 6.08 - 6.06 (m, 1H), 5.04 - 4.62 (m, 1H), 4.14 - 4.12 (m, 2H), 3.21 (t, *J* = 2.5 Hz, 1H), 2.30 - 2.22 (m, 4H), 1.02 (d, *J* = 6.7 Hz, 2H), 1.02 (d, *J* = 6.7 Hz, 2H), 0.88 (d, *J* = 6.7 Hz, 2H); $^{13}$C NMR (151 MHz, DMSO-$d_6$, δ [ppm]): 169.3, 169.2, 155.1, 154.8, 153.6, 153.5, 153.2, 153.2, 151.6, 151.4, 150.3, 149.7, 128.9, 128.7, 125.5, 125.4, 125.0, 124.9, 118.2, 114.5, 114.4, 109.3, 108.7, 106.5, 106.4, 80.2, 73.5, 58.7, 52.2, 34.3, 31.1, 28.4, 19.2, 19.2, 18.8, 13.4, 13.4, 13.3; HRMS m/z (ESI$^+$) [found: 458.1493, $C_{21}H_{24}N_5O_5S^+$ requires [M + H]$^+$ 458.1420]; HPLC retention time: 12.59 min, purity: 97.9%

**(*R*)-2-Hydroxy-3-[(4-{[1-(5-methylfuran-2-yl)propyl]amino}-1,1-dioxido-1,2,5-thiadiazol-3-yl)amino]-*N*-(prop-2-yn-1-yl)benzamide (20d)**

2-Hydroxy-3-[(4-methoxy-1,1-dioxido-1,2,5-thiadiazol-3-yl)amino]-*N*-(prop-2-yn-1-yl)benzamide (**18b**, 130 mg, 0.370 mmol, 1.00 *eq.*) and (*R*)-1-(5-methylfuran-2-yl)propan-1-amine hydrochloride (**15a**, 77 mg, 0.52 mmol, 1.4 *eq.*) were dissolved in methanol (10 mL). After the addition of *N,N*-diisopropylethylamine (132 µL, 0.740 mmol, 2.00 *eq.*), the reaction was stirred for 3 days. Extraction between water and ethyl acetate (3 × 50 mL), drying over sodium sulfate, filtration, and evaporation of the solvent gave the crude product which was purified by preparative HPLC (acetonitrile/water (0.1% TFA): gradient 5−95%) to obtain the title compound as a white-brown amorphous solid (37 mg, 22%). $^1$H NMR (600 MHz, DMSO-$d_6$, δ [ppm]): 13.61 (s, 1H), 10.14 (s, 1H), 9.66 (d, *J* = 7.9 Hz, 1H), 9.53 (t, *J* = 5.5 Hz, 1H), 7.99 (dd, *J* = 8.1, 1.4 Hz, 1H), 7.81 (dd, *J* = 8.2, 1.5 Hz, 1H), 7.02 (t, *J* = 8.0 Hz, 1H), 6.37 (d, *J* = 3.1 Hz, 1H), 6.08 (d, *J* = 3.1 Hz, 1H), 4.60 (q, *J* = 7.5 Hz, 1H), 4.13 (dd, *J* = 5.5, 2.5 Hz, 2H), 3.21 (t, *J* = 2.5 Hz, 1H), 2.28 (s, 3H), 2.04 - 1.88 (m, 2H), 1.00 - 0.81 (m, 3H); $^{13}$C NMR (151 MHz, DMSO-$d_6$, δ [ppm]): 169.3, 163.1, 154.8, 154.0, 153.5, 153.2, 151.6, 151.2, 150.4, 150.1, 128.7, 125.5, 124.9, 118.2, 114.4, 108.9, 108.3, 106.5, 106.4, 80.2, 73.5, 70.2, 53.9, 53.7, 28.4, 25.5, 25.0, 13.6, 13.3, 13.3, 10.6, 10.5; HRMS m/z (ESI$^+$) [found: 444.1336, $C_{20}H_{22}N_5O_5S^+$ requires [M + H]$^+$ 444.1297]; HPLC retention time: 12.44 min, purity: 95.4%.

**2-Hydroxy-3-{[2-(neopentylamino)-3,4-dioxocyclobut-1-en-1-yl]amino}-*N*-(prop-2-yn-1-yl)benzamide (21a)**

2-Hydroxy-3-[(2-methoxy-3,4-dioxocyclobut-1-en-1-yl)amino]-*N*-(prop-2-yn-1-yl)benzamide (**19**, 37 mg, 0.12 mmol, 1.0 *eq.*) and neopentylamine (22 µL, 0.18 mmol, 1.5 *eq.*) were dissolved in methanol (10 mL). After the addition of *N,N*-diisopropylethylamine (44 µL, 0.24 mmol, 2.0 *eq.*), the reaction was stirred for 3 days. Extraction between water and ethyl acetate (3 × 50 mL), drying over sodium sulfate, filtration, and evaporation of the solvent gave the crude product which was purified by preparative HPLC (acetonitrile/water (0.1% TFA): gradient 5−95%) to obtain the title compound as a white-brown amorphous solid (20 mg, 45%). $^1$H NMR (500 MHz, DMSO-$d_6$, δ [ppm]): 13.67 (s, 1H), 9.47 (t, *J* = 5.6 Hz, 1H), 9.40 (s, 1H), 8.36 (t, *J* = 6.8 Hz, 1H), 8.04 - 7.99 (m, 1H), 7.54 (d, *J* = 8.2 Hz, 1H), 6.90 (t, *J* = 8.1 Hz, 1H), 4.12 (dd, *J* = 5.6, 2.5 Hz, 2H), 3.43 (d, *J* = 6.7 Hz, 2H), 3.20 (t, *J* = 2.5 Hz, 1H), 0.93 (s, 9H); $^{13}$C NMR (126 MHz, DMSO-$d_6$, δ [ppm]): 184.4, 180.1, 170.0, 169.8, 162.8, 150.9, 128.1, 123.4, 120.5, 118.3, 113.4, 80.3, 73.4, 54.8, 32.2, 28.4, 26.5; HRMS m/z (ESI$^+$) [found: 356.1605, $C_{19}H_{21}N_3O_4^+$ requires [M + H]$^+$ 356.1612]; HPLC retention time: 12.62 min, purity: 95.0%.

**3-{[2-(Cyclobutylamino)-3,4-dioxocyclobut-1-en-1-yl]amino}-2-hydroxy-*N*-(prop-2-yn-1-yl)benzamide (21b)**

2-Hydroxy-3-[(2-methoxy-3,4-dioxocyclobut-1-en-1-yl)amino]-*N*-(prop-2-yn-1-yl)benzamide (**19**, 30 mg, 0.075 mmol, 1.0 *eq.*) and cyclobutylamine (11 µL, 0.075 mmol, 1.0 *eq.*) were dissolved in methanol (10 mL). After the addition of *N,N*-diisopropylethylamine (36 µL, 0.15 mmol, 2.0 *eq.*), the reaction was stirred for 3 days. Extraction between water and ethyl acetate (3 × 50 mL), drying over sodium sulfate, filtration, and evaporation of the solvent gave the crude product which was purified by preparative HPLC (acetonitrile/water (0.1% TFA): gradient 5−95%) to obtain the title compound as a white-brown amorphous solid (28 mg, 82%). $^{1}$H NMR (500 MHz, DMSO-$d_6$, δ [ppm]): 13.63 (s, 1H), 9.46 (t, *J* = 5.6 Hz, 1H), 9.25 (s, 1H), 8.66 (d, *J* = 8.4 Hz, 1H), 8.00 (d, *J* = 8.0 Hz, 1H), 7.54 (dd, *J* = 8.2, 1.4 Hz, 1H), 6.90 (t, *J* = 8.0 Hz, 1H), 4.60 - 4.52 (m, 1H), 4.12 - 4.11 (m, 2H), 3.19 (t, *J* = 2.5 Hz, 1H), 2.37 - 2.30 (m, 2H), 2.10 - 1.97 (m, 2H), 1.77 - 1.59 (m, 2H); $^{13}$C NMR (126 MHz, DMSO-$d_6$, δ [ppm]): 184.2, 180.1, 169.8, 168.3, 162.9, 150.8, 128.1, 123.3, 120.6, 118.3, 113.4, 80.3, 73.4, 48.7, 31.6, 28.4, 14.0; HRMS m/z (ESI$^+$) [found: 340.1292, $C_{18}H_{18}N_3O_4^+$ requires [M + H]$^+$ 340.1296]; HPLC retention time: 12.12 min, purity: 96.8%.

**3-{[2-(Cyclopentylamino)-3,4-dioxocyclobut-1-en-1-yl]amino}-2-hydroxy-*N*-(prop-2-yn-1-yl)benzamide (21c)**

2-Hydroxy-3-[(2-methoxy-3,4-dioxocyclobut-1-en-1-yl)amino]-*N*-(prop-2-yn-1-yl)benzamide (**19**, 30 mg, 0.075 mmol, 1.0 *eq.*) and cyclopentylamine (13 µL, 0.075 mmol, 1.0 *eq.*) were dissolved in methanol (10 mL). After the addition of *N,N*-diisopropylethylamine (36 µL, 0.15 mmol, 2.0 *eq.*), the reaction was stirred for 3 days. Extraction between water and ethyl acetate (3 × 50 mL), drying over sodium sulfate, filtration, and evaporation of the solvent gave the crude product which was purified by preparative HPLC (acetonitrile/water (0.1% TFA): gradient 5−95%) to obtain the title compound as a white-brown amorphous solid (32 mg, 91%). $^{1}$H NMR (500 MHz, DMSO-$d_6$, δ [ppm]): 13.65 (s, 1H), 9.46 (t, *J* = 5.6 Hz, 1H), 9.24 (s, 1H), 8.38 (d, *J* = 7.9 Hz, 1H), 8.01 (d, *J* = 8.0 Hz, 1H), 7.53 (dd, *J* = 8.2, 1.4 Hz, 1H), 6.90 (t, *J* = 8.1 Hz, 1H), 4.46 - 4.40 (m, 1H), 4.12 - 4.10 (m, 2H), 3.19 (t, *J* = 2.5 Hz, 1H), 2.00 - 1.92 (m, 2H), 1.78 - 1.50 (m, 6H); $^{13}$C NMR (126 MHz, DMSO-$d_6$, δ [ppm]) 184.3, 179.9, 169.8, 168.8, 163.0, 150.8, 128.1, 123.3, 120.5, 118.3, 113.4, 80.3, 73.4, 55.4, 33.7, 28.4, 23.2; HRMS m/z (ESI$^+$) [found: 354.1448, $C_{19}H_{20}N_3O_4^+$ requires [M + H]$^+$ 354.1455]; HPLC retention time: 12.31 min, purity: 97.3%.

**3-{[2-(Cyclohexylamino)-3,4-dioxocyclobut-1-en-1-yl]amino}-2-hydroxy-*N*-(prop-2-yn-1-yl)benzamide (21d)**

2-Hydroxy-3-[(2-methoxy-3,4-dioxocyclobut-1-en-1-yl)amino]-*N*-(prop-2-yn-1-yl)benzamide (**19**, 30 mg, 0.075 mmol, 1.0 *eq.*) and cyclohexylamine (15 µl, 0.075 mmol, 1.0 *eq.*) were dissolved in methanol (10 mL). After the addition of *N,N*-diisopropylethylamine (36 µL, 0.15 mmol, 2.0 *eq.*), the reaction was stirred for 3 days. Extraction between water and ethyl acetate (3 × 50 mL), drying over sodium sulfate, filtration, and evaporation of the solvent gave the crude product which was purified by preparative HPLC (acetonitrile/water (0.1% TFA): gradient 5−95%) to obtain the title compound as a white-brown amorphous solid (27 mg, 73%). $^{1}$H NMR (500 MHz, DMSO-$d_6$, δ [ppm]): 13.64 (s, 1H), 9.46 (t, *J* = 5.6 Hz, 1H), 9.30 (s, 1H), 8.38 (d, *J* = 8.0 Hz, 1H), 8.00 (d, *J* = 8.0 Hz, 1H), 7.53 (dd, *J* = 8.1, 1.3 Hz, 1H), 6.90 (t, *J* = 8.1 Hz, 1H), 4.12 - 4.10 (m, 2H), 3.94 - 3.88 (m, 1H), 3.19 (t, *J* = 2.5 Hz, 1H), 1.96 - 1.91 (m, 2H), 1.75 - 1.69 (m, 2H), 1.59 - 1.54 (m, 1H), 1.40 - 1.15 (m, 5H); $^{13}$C NMR (126 MHz, DMSO-$d_6$, δ [ppm]): 184.2, 179.9, 169.8, 168.6, 163.0, 150.9, 128.1, 123.4, 120.5, 118.3, 113.4, 80.3, 73.4, 52.4, 33.6, 28.4,

24.7, 23.9; HRMS m/z (ESI$^+$) [found: 368.1605, $C_{20}H_{22}N_3O_4^+$ requires $[M + H]^+$ 368.1610]; HPLC retention time: 12.56 min, purity: 97.6%.

**3-{[2-(Adamantan-1-ylamino)-3,4-dioxocyclobut-1-en-1-yl]amino}-2-hydroxy-*N*-(prop-2-yn-1-yl)benzamide (21e)**

2-Hydroxy-3-[(2-methoxy-3,4-dioxocyclobut-1-en-1-yl)amino]-*N*-(prop-2-yn-1-yl)benzamide (**19**, 30 mg, 0.075 mmol, 1.0 *eq.*) and 1-adamantylamine (20 mg, 0.075 mmol, 1.0 *eq.*) were dissolved in methanol (10 mL). After the addition of *N*,*N*-diisopropylethylamine (36 µL, 0.15 mmol, 2.0 *eq.*), the reaction was stirred for 3 days. Extraction between water and ethyl acetate (3 × 50 mL), drying over sodium sulfate, filtration, and evaporation of the solvent gave the crude product which was purified by preparative HPLC (acetonitrile/water (0.1% TFA): gradient 5−95%) to obtain the title compound as a white-brown amorphous solid (24 mg, 57%). $^1$H NMR (500 MHz, DMSO-$d_6$, δ [ppm]): 13.65 (s, 1H), 9.50 - 9.41 (m, 2H), 8.52 (s, 1H), 7.94 (dd, *J* = 8.0, 1.4 Hz, 1H), 7.55 (dd, *J* = 8.3, 1.4 Hz, 1H), 6.90 (t, *J* = 8.1 Hz, 1H), 4.12 - 4.11 (m, 2H), 3.19 (t, *J* = 2.5 Hz, 1H), 2.13 - 2.07 (m, 3H), 2.03 - 1.98 (m, 6H), 1.69 - 1.64 (m, 6H); $^{13}$C NMR (126 MHz, DMSO-$d_6$, δ [ppm]): 182.9, 179.9, 169.8, 169.4, 164.0, 151.1, 127.9, 123.9, 120.8, 118.2, 113.4, 80.3, 73.4, 52.8, 42.4, 35.3, 29.0, 28.4; HRMS m/z (ESI$^+$) [found: 420.1918, $C_{24}H_{26}N_3O_4^+$ requires $[M + H]^+$ 420.1918]; HPLC retention time: 13.41 min, purity: 95.9%.

**3-({2-[(Dicyclopropylmethyl)amino]-3,4-dioxocyclobut-1-en-1-yl}amino)-2-hydroxy-*N*-(prop-2-yn-1-yl)benzamide (21f)**

2-Hydroxy-3-[(2-methoxy-3,4-dioxocyclobut-1-en-1-yl)amino]-*N*-(prop-2-yn-1-yl)benzamide (**19**, 30 mg, 0.075 mmol, 1.0 *eq.*) and dicyclopropylmethanamine (13 µL, 0.075 mmol, 1.0 *eq.*) were dissolved in methanol (10 mL). After the addition of *N*,*N*-diisopropylethylamine (36 µL, 0.15 mmol, 2.0 *eq.*), the reaction was stirred for 3 days. Extraction between water and ethyl acetate (3 × 50 mL), drying over sodium sulfate, filtration, and evaporation of the solvent gave the crude product which was purified by preparative HPLC (acetonitrile/water (0.1% TFA): gradient 5−95%) to obtain the title compound as a white-brown amorphous solid (20 mg, 53%) $^1$H NMR (500 MHz, DMSO-$d_6$, δ [ppm]): 13.64 (s, 1H), 9.47 (t, *J* = 5.5 Hz, 1H), 9.36 (s, 1H), 8.50 (d, *J* = 9.3 Hz, 1H), 8.01 (d, *J* = 7.9 Hz, 1H), 7.54 (d, *J* = 8.1 Hz, 1H), 6.90 (t, *J* = 8.0 Hz, 1H), 4.13 - 4.09 (m, 2H), 3.22 - 3.19 (m, 1H), 3.13 (q, *J* = 8.3 Hz, 1H), 1.13 - 1.04 (m, 2H), 0.59 - 0.53 (m, 2H), 0.47 - 0.44 (m, 2H), 0.37 - 0.34 (m, 4H); $^{13}$C NMR (126 MHz, DMSO-$d_6$, δ [ppm]): 184.1, 179.7, 169.8, 168.6, 163.0, 150.9, 128.1, 123.5, 120.6, 118.3, 113.4, 80.3, 73.4, 60.9, 28.4, 15.7, 2.9, 1.9; HRMS m/z (ESI$^+$) [found: 380.1605, $C_{21}H_{22}N_3O_4^+$ requires $[M + H]^+$ 380.1610; HPLC retention time: 12.47 min, purity: 97.4%.

**3-{[2-(Benzhydrylamino)-3,4-dioxocyclobut-1-en-1-yl)amino}-2-hydroxy-*N*-(prop-2-yn-1-yl)benzamide (21g)**

2-Hydroxy-3-[(2-methoxy-3,4-dioxocyclobut-1-en-1-yl)amino]-*N*-(prop-2-yn-1-yl)benzamide (**19**, 30 mg, 0.075 mmol, 1.0 *eq.*) and benzhydrylamine (24 mg, 0.075 mmol, 1.0 *eq.*) were dissolved in methanol (10 mL). After the addition of *N*,*N*-diisopropylethylamine (36 µL, 0.15 mmol, 2.0 *eq.*), the reaction was stirred for 3 days. Extraction between water and ethyl acetate (3 × 50 mL), drying over sodium sulfate, filtration, and evaporation of the solvent gave the crude product which was purified by preparative HPLC (acetonitrile/water (0.1% TFA): gradient 5−95%) to obtain the title compound as a white-brown amorphous solid (35 mg, 78%). $^1$H NMR (500 MHz, DMSO-$d_6$, δ [ppm]): 13.66 (s, 1H), 9.50 - 9.42 (m, 2H), 9.15 (d, *J* = 8.8 Hz, 1H), 8.00 (d, *J* = 7.9 Hz, 1H), 7.55 (dd, *J* = 8.2, 1.4 Hz, 1H), 7.46 - 7.38 (m, 4H), 7.36 - 7.29 (m, 6H), 6.91

(t, $J$ = 8.1 Hz, 1H), 6.52 (d, $J$ = 8.8 Hz, 1H), 4.14 - 4.09 (m, 2H), 3.20 (t, $J$ = 2.5 Hz, 1H); $^{13}$C NMR (126 MHz, DMSO-$d_6$, δ [ppm]): 184.0, 180.6, 169.8, 168.4, 163.5, 150.9, 141.6, 128.8, 128.0, 127.6, 127.0, 123.4, 120.7, 118.3, 113.5, 80.3, 73.4, 60.6, 28.4; HRMS m/z (ESI$^+$) [found: 452.1605, $C_{27}H_{22}N_3O_4^+$ requires [M + H]$^+$ 452.1611]; HPLC retention time: 12.87 min, purity: 97.8%.

**3-[(2-{[Di(pyridin-2-yl)methyl]amino}-3,4-dioxocyclobut-1-en-1-yl)amino]-2-hydroxy-*N*-(prop-2-yn-1-yl)benzamide (21h)**

2-Hydroxy-3-[(2-methoxy-3,4-dioxocyclobut-1-en-1-yl)amino]-*N*-(prop-2-yn-1-yl)benzamide (**19**, 30 mg, 0.075 mmol, 1.0 *eq.*) and bis(pyridine-2-yl)methanamine (25 mg, 0.075 mmol, 1.0 *eq.*) were dissolved in methanol (10 mL). After the addition of *N,N*-diisopropylethylamine (36 µL, 0.15 mmol, 2.0 *eq.*), the reaction was stirred for 3 days. Extraction between water and ethyl acetate (3 × 50 mL), drying over sodium sulfate, filtration, and evaporation of the solvent gave the crude product which was purified by preparative HPLC (acetonitrile/water (0.1% TFA): gradient 5−95%) to obtain the title compound as a white-brown amorphous solid (23 mg, 51%). $^1$H NMR (500 MHz, DMSO-$d_6$, δ [ppm]): 13.62 (s, 1H), 9.88 (s, 1H), 9.66 (d, $J$ = 9.0 Hz, 1H), 9.47 (t, $J$ = 5.6 Hz, 1H), 8.60 - 8.56 (m, 2H), 7.94 (d, $J$ = 8.0 Hz, 1H), 7.88 (td, $J$ = 7.7, 1.8 Hz, 2H), 7.63 - 7.53 (m, 3H), 7.38 - 7.34 (m, 2H), 6.90 (t, $J$ = 8.0 Hz, 1H), 6.72 (d, $J$ = 9.0 Hz, 1H), 4.15 - 4.08 (m, 2H), 3.20 (t, $J$ = 2.5 Hz, 1H); $^{13}$C NMR (126 MHz, DMSO-$d_6$, δ [ppm]): 184.3, 180.8, 169.8, 168.4, 163.8, 158.8, 151.2, 148.9, 137.7, 128.0, 123.8, 123.0, 122.2, 120.8, 118.2, 113.5, 80.3, 73.4, 62.5, 28.4; HRMS m/z (ESI$^+$) [found: 454.1510, $C_{25}H_{320}N_5O_4^+$ requires [M + H]$^+$ 454.1507]; HPLC retention time: 11.19 min, purity: 97.9%.

**(*R*)-3-[(2-{[2,2-Dimethyl-1-(pyrimidin-5-yl)propyl]amino}-3,4-dioxocyclobut-1-en-1-yl)amino]-2-hydroxy-*N*-(prop-2-yn-1-yl)benzamide (21i)**

2-Hydroxy-3-[(2-methoxy-3,4-dioxocyclobut-1-en-1-yl)amino]-*N*-(prop-2-yn-1-yl)benzamide (**19**, 40 mg, 0.10 mmol, 1.0 *eq.*) and (*R*)-2,2-dimethyl-1-(pyrimidin-5-yl)propan-1-amine hydrochloride (**15e**, 27 mg, 0.10 mmol, 1.0 *eq.*) were dissolved in methanol (10 mL). After the addition of *N,N*-diisopropylethylamine (71 µL, 0.30 mmol, 3.0 *eq.*), the reaction was stirred for 3 days. Extraction between water and ethyl acetate (3 × 50 mL), drying over sodium sulfate, filtration, and evaporation of the solvent gave the crude product which was purified by preparative HPLC (acetonitrile/water (0.1% TFA): gradient 5−95%) to obtain the title compound as a white-brown amorphous solid (17 mg, 29%). $^1$H NMR (500 MHz, DMSO-$d_6$, δ [ppm]): 13.77 (s, 1H), 9.51 - 9.46 (m, 2H), 9.14 (s, 1H), 8.77 (s, 2H), 8.68 (d, $J$ = 9.3 Hz, 1H), 7.92 (d, $J$ = 8.7 Hz, 1H), 7.56 (dd, $J$ = 8.2, 1.4 Hz, 1H), 6.90 (t, $J$ = 8.1 Hz, 1H), 5.13 (d, $J$ = 9.3 Hz, 1H), 4.12 (dd, $J$ = 5.6, 2.5 Hz, 2H), 3.20 (t, $J$ = 2.5 Hz, 1H), 0.99 (s, 9H); $^{13}$C NMR (126 MHz, DMSO-$d_6$, δ [ppm]): 183.9, 180.6, 169.8, 168.9, 163.8, 157.4, 156.0, 150.9, 133.5, 127.8, 123.7, 120.9, 118.3, 113.5, 80.3, 73.4, 62.6, 35.0, 28.4, 25.8; HRMS m/z (ESI$^+$) [found: 434.1823, $C_{23}H_{24}N_5O_4^+$ requires [M + H]$^+$ 434.1824]; HPLC retention time: 11.93 min, purity: 96.8%.

**(*R*)-3-[(2-{[1-(2,4-Dichlorothiazol-5-yl)-2,2-dimethylpropyl]amino}-3,4-dioxocyclobut-1-en-1-yl)amino]-2-hydroxy-*N*-(prop-2-yn-1-yl)benzamide (21j)**

2-Hydroxy-3-[(2-methoxy-3,4-dioxocyclobut-1-en-1-yl)amino]-*N*-(prop-2-yn-1-yl)benzamide (**19**, 40 mg, 0.10 mmol, 1.0 *eq.*) and (*R*)-1-(2,4-dichlorothiazol-5-yl)-2,2-dimethylpropan-1-amine hydrochloride (**15f**, 37 mg, 0.10 mmol, 1.0 *eq.*) were dissolved in methanol (10 mL). After the addition of *N,N*-diisopropylethylamine (71 µL, 0.30 mmol, 3.0 *eq.*), the reaction was stirred for 3 days. Extraction between water and ethyl acetate (3 × 50 mL), drying over sodium sulfate, filtration, and evaporation of the solvent gave the crude product which was purified by preparative HPLC (acetonitrile/water (0.1% TFA): gradient 5−95%) to obtain the title compound as a white-brown amorphous solid (19 mg, 28%). $^1$H NMR (500 MHz, DMSO-$d_6$, δ [ppm]): 13.75 (s, 1H), 9.49 (t, *J* = 5.6 Hz, 1H), 9.41 (s, 1H), 8.56 (d, *J* = 9.5 Hz, 1H), 7.93 (dd, *J* = 8.1, 1.3 Hz, 1H), 7.57 (dd, *J* = 8.2, 1.4 Hz, 1H), 6.91 (t, *J* = 8.1 Hz, 1H), 5.56 (d, *J* = 9.4 Hz, 1H), 4.12 (dd, *J* = 5.5, 2.5 Hz, 2H), 3.20 (t, *J* = 2.5 Hz, 1H), 1.05 (s, 9H); $^{13}$C NMR (126 MHz, DMSO-$d_6$, δ [ppm]): 183.6, 180.7, 169.8, 168.4, 163.6, 151.0, 149.9, 134.3, 133.1, 127.7, 123.7, 120.9, 118.3, 113.5, 80.3, 73.4, 59.8, 36.4, 28.4, 25.7; HRMS m/z (ESI$^+$) [found: 507.0669, $C_{22}H_{21}Cl_2N_4O_4S^+$ requires [M + H]$^+$ 507.0655]; HPLC retention time: 13.18 min, 98.2%.

**(*R*)-3-{[2-({1-[5-(4-Chlorophenyl)isoxazol-3-yl]-2,2-dimethylpropyl}amino)-3,4-dioxocyclobut-1-en-1-yl]amino}-2-hydroxy-*N*-(prop-2-yn-1-yl)benzamide (21k)**

2-Hydroxy-3-[(2-methoxy-3,4-dioxocyclobut-1-en-1-yl)amino]-*N*-(prop-2-yn-1-yl)benzamide (**19**, 40 mg, 0.10 mmol, 1.0 *eq.*) and (*R*)-1-[5-(4-chlorophenyl)isoxazol-3-yl]-2,2-dimethylpropan-1-amine hydrochloride (**15g**, 40 mg, 0.10 mmol, 1.0 *eq.*) were dissolved in methanol (10 mL). After the addition of *N,N*-diisopropylethylamine (71 µL, 0.30 mmol, 3.0 *eq.*), the reaction was stirred for 3 days. Extraction between water and ethyl acetate (3 × 50 mL), drying over sodium sulfate, filtration, and evaporation of the solvent gave the crude product which was purified by preparative HPLC (acetonitrile/water (0.1% TFA): gradient 5−95%) to obtain the title compound as a white-brown amorphous solid (40 mg, 56%). $^1$H NMR (500 MHz, DMSO-$d_6$, δ [ppm]): 13.70 (s, 1H), 9.62 (s, 1H), 9.48 (t, *J* = 5.6 Hz, 1H), 8.91 (d, *J* = 10.1 Hz, 1H), 7.97 (dd, *J* = 8.0, 1.3 Hz, 1H), 7.95 - 7.89 (m, 2H), 7.65 - 7.59 (m, 2H), 7.56 (dd, *J* = 8.2, 1.4 Hz, 1H), 7.12 (s, 1H), 6.90 (t, *J* = 8.0 Hz, 1H), 5.35 (d, *J* = 10.1 Hz, 1H), 4.12 (dd, *J* = 5.5, 2.5 Hz, 2H), 3.20 (t, *J* = 2.5 Hz, 1H), 1.05 (s, 9H); $^{13}$C NMR (126 MHz, DMSO-$d_6$, δ [ppm]): 184.0, 180.4, 169.8, 168.7, 167.7, 163.3, 163.1, 151.0, 135.2, 129.3, 127.9, 127.5, 125.4, 123.7, 120.8, 118.3, 113.5, 101.0, 80.3, 73.4, 59.5, 35.4, 28.4, 25.9; HRMS m/z (ESI$^+$) [found: 533.1601, $C_{28}H_{26}ClN_4O_5^+$ requires [M + H]$^+$ 533.1586]; HPLC retention time: 13.65 min, purity: 95.6%.

**(*R*)-3-[(2-{[2,2-Dimethyl-1-(5-methylthiophen-2-yl)propyl]amino}-3,4-dioxocyclobut-1-en-1-yl)amino]-2-hydroxy-*N*-(prop-2-yn-1-yl)benzamide (21l)**

2-Hydroxy-3-[(2-methoxy-3,4-dioxocyclobut-1-en-1-yl)amino]-*N*-(prop-2-yn-1-yl)benzamide (**19**, 40 mg, 0.10 mmol, 1.0 *eq.*) and (*R*)-2,2-dimethyl-1-(5-methylthiophen-2-yl)propan-1-amine hydrochloride (**15d**, 29 mg, 0.10 mmol, 1.0 *eq.*) were dissolved in methanol (10 mL). After the addition of *N,N*-diisopropylethylamine (71 µL, 0.30 mmol, 3.0 *eq.*), the reaction was stirred for 3 days. Extraction between water and ethyl acetate (3 × 50 mL), drying over sodium sulfate, filtration, and evaporation of the solvent gave the crude product which was purified by preparative HPLC (acetonitrile/water (0.1% TFA): gradient 5−95%) to obtain the title compound as a white-brown amorphous solid (44 mg, 73%). $^1$H NMR (500 MHz, DMSO-$d_6$, δ [ppm]): 13.72 (s, 1H), 9.52 - 9.45 (m, 2H), 8.62 (d, *J* = 10.0 Hz, 1H), 7.95 (d, *J* = 8.1 Hz, 1H), 7.55 (dd, *J* = 8.1, 1.4 Hz, 1H), 6.90 (t, *J* = 8.1 Hz, 1H), 6.78 (d, *J* = 3.5 Hz, 1H), 6.70 (dd, *J* = 3.4, 1.1 Hz, 1H), 5.32 (d, *J* = 10.0 Hz, 1H), 4.12 (dd, *J* = 5.5, 2.5 Hz, 2H), 3.20 (t, *J* = 2.5 Hz, 1H), 2.42 (d, *J* = 1.1 Hz, 3H), 1.01 (s, 9H); $^{13}$C NMR (126 MHz, DMSO-$d_6$, δ [ppm]): 184.1, 180.2, 169.8, 168.5, 163.2, 151.0, 139.9, 138.5, 127.9, 126.3, 124.8, 123.7, 120.8, 118.3, 113.4, 80.3, 73.4, 62.8, 35.4, 28.4, 26.2, 14.8; HRMS m/z (ESI$^+$) [found: 452.1648, $C_{24}H_{26}N_3O_4S^+$ requires [M + H]$^+$ 452.1639]; HPLC retention time: 13.22 min, purity: 97.9%.

**(*R*)-3-[(2-{[2,2-Dimethyl-1-(5-methylfuran-2-yl)propyl]amino}-3,4-dioxocyclobut-1-en-1-yl)amino]-2-hydroxy-*N*,*N*-dimethylbenzamide (SLW132, 21m)**

2-Hydroxy-3-[(2-methoxy-3,4-dioxocyclobut-1-en-1-yl)amino]-*N*,*N*-dimethylbenzamide (**19b**, 60 mg, 0.2 mmol, 1.0 *eq.*) and (*R*)-2,2-dimethyl-1-(5-methylfuran-2-yl)propan-1-amine hydrochloride (**15c**, 84 mg, 0.4 mmol, 2.0 *eq.*) were dissolved in methanol (10 mL). After the addition of *N*,*N*-diisopropylethylamine (73 μL, 0.4 mmol, 2.0 *eq.*), the reaction was stirred for 3 days. Extraction between water and ethyl acetate (3 × 50 mL), drying over sodium sulfate, filtration, and evaporation of the solvent gave the crude product that was purified by preparative HPLC (acetonitrile/water (0.1% TFA): gradient 5−95%) to obtain compound 19 as a white-brown amorphous solid (43 mg, 49%) $^{1}$H NMR (400 MHz, DMSO-$d_6$, δ [ppm]): 9.97 (bs, 1H), 9.48 (s, 1H), 8.75 (d, $J$ = 10.12 Hz, 1H), 7.76 – 7.72 (m, 1H), 6.91 – 6.85 (m, 2H), 6.19 (d, $J$ = 3.12 Hz, 1H), 6.06 – 6.03 (m, 1H), 5.12 (d, $J$ = 10.09 Hz, 1H), 2.95 (s, 6H), 2.28 (s, 3H), 0.98 (s, 9H); $^{13}$C NMR (151 MHz, DMSO-$d_6$, δ [ppm]): 184.1, 180.3, 168.6, 168.3, 163.4, 151.0, 150.8, 143.5, 128.5, 124.4, 122.3, 121.2, 119.8, 108.5, 106.3, 60.2, 40.1, 35.7, 26.2, 13.4; HRMS *m*/*z* (ESI$^+$) [found: 426.2028, $C_{23}H_{28}N_3O_5^+$ requires [M+H]$^+$ 426.1951]; LRMS *m/z* (ESI$^+$) [found: 426.4, $C_{23}H_{28}N_3O_5^+$ requires [M+H]$^+$ 426.5]; HPLC retention time: 17.92 min, 98.0%.

#### Molecular modeling studies

*Structure visualization* was done in Molecular Operating Environment (MOE, version 2022.02).[10] The molecular surface was calculated at 4.5 Å around the *tert*-butyl group of Cmp2105 (**1**).

*Protein preparation.* The protein structure (PDB ID: 6QZH) was prepared in MOE using default setting. This preparation included adding missing hydrogens and protonating titratable groups with Protonate3D. N- and C-termini, as well as sequence breaks, were capped, missing sidechains were rebuilt by default. No energy minimization was performed. The prepared structure was processed with MakeReceptor (version 4.2.1.1)[11] to make it compatible with other OpenEye tools. Hydrogens were left untouched, and the tautomer option was switched off. The box delimiting grids was created based on Cmp2105 (**1**) and default settings were applied to create the site shape potential.

*Ligand preparation.* Protonation states were generated with fixpKa and tautomers (QUACPAC, version 2.2.2.0),[12] and the most probable protonation state was chosen considering the proximity of Asp94 in the 6QZH crystal structure. Stereoisomers were generated with flipper (OMEGA, version 4.2.2.0)[13] and ligand conformations were generated with the conformer_generator from CCDC (version 2023.3.0 CSD Portfolio 2023 release).[14]

*Molecular docking calculations*. Molecular docking calculations were performed with FRED (version 4.2.1.0).[15] The top-scored pose for compound **3** (score -12.048) was minimized with SZYBKI (version 2.6.0.0)[16] using steepest-descent optimization. Residual steric clashes were removed with MOE's Energy Minimization module using only ligand atoms as input and the MMFF94x force field. Restraints were applied with a maximum allowed deviation of 1.5 Å from initial coordinates to discourage significant shifts from the docking-derived pose.

*SZMAP calculations*. Atom charges and radii were assigned to the structure of the protein (PDB ID: 6QZH) using MakeReceptor. Desmethylated ligand **10** was prepared by removing methyl from the crystal structure of Cmp2105 (**1**). AM1BCC charges and Zap9 radii were assigned to ligands **1** and **10**. Grids were calculated separately for the methylated and desmethylated ligand, respectively, using default settings of SZMAP (version 1.6.6.0)[17] and adding stabilization calculations. Ligand displacement grids were calculated separately using the grid_comp tool. Additionally, the difference between the grids for water - neutral probe free energies were calculated using grid_comp for both ligand **1** and **10**. Grids were visualized using VIDA (version 5.0.5).[18] Free energy values for the water molecules were visualized with the WaterColor VIDA extension. Orientations of the predicted water molecules were sampled with the Water Orientation VIDA extension.

#### Biological evaluation

##### NanoBRET assays

*Cell culture:* HEK293T cells were cultured as previously described.[19] In brief, HEK293T cells (gift from Chair of Physiology, Prof. Dr. Alzheimer, FAU Erlangen-Nürnberg) were grown on 10 cm culture dishes at 37 °C and 5% $CO_2$. As growth medium, Dulbecco's Modified Eagle Medium (DMEM)/F12 (Invitrogen) supplemented with 10% fetal bovine serum (FBS, Gibco™ Fetal

Bovine Serum, qualified, Brazil), L-glutamine (final concentration: 2 mM; from Gibco™ L-glutamine 200 mM, 100X), penicillin (final concentration: 100 units/mL), and streptomycin (final concentration: 100 µg/mL; from Gibco™ Penicillin-Streptomycin 10,000 U/mL) was used. Cells were split every three to four days and regularly confirmed to be free of mycoplasma contamination using the luminescence-based MycoAlert Plus Kit (Lonza).

*Transient transfection using polyethylenimine:* Transfection was performed as previously described.[19] For transfection, we used a previously published procedure.[19] In brief, HEK293T cells were plated onto culture dishes (Ø 10 cm or Ø 15 cm) and grown to a confluence of approximately 50% at 37 °C and 5% $CO_2$. The growth medium was renewed one-hour before transfection. The transfection mix was prepared, as described in the following. Solution A (2.1-2.2% total DNA in Gibco™ phosphate buffered saline, pH 7.4 [5.5 µg in 250 µL, 10.5 µg in 500 µL]) and solution B (3% PEI (linear, 25 kDa, from Polysciences) solution prepared from a PEI stock solution (1 µg/µL) in PBS without $MgCl_2$ and $CaCl_2$) were mixed one to one, the resulting mixture was vortexed for 5 s and incubated for 30 min at room temperature. The pre-incubated transfection mix was added dropwise to the cells and cell cultivation was continued at 37 °C and 5% $CO_2$.

*Membrane preparation:* Membranes were prepared as previously reported.[19] In brief, membranes from HEK293T cells transiently expressing the respective GPCR were prepared as follows. The medium of the transfected cells was refreshed after 24 h before cells were harvested 48 h post-transfection. The growth medium was removed, the cells were carefully washed with cold phosphate buffered saline (10 mL per Ø 15 cm dish). The cells were detached with 15 mL of ice-cold Tris-EDTA buffer (10 mM Tris, 0.5 mM EDTA, 5.4 mM KCl, 140 mM NaCl, pH 7.4) and subsequently centrifuged with 218 *g* for 8 minutes. The supernatant was removed, and the cells were resuspended in 10 mL Tris-EDTA buffer. The cells were lysed with an Ultraturrax (20,000 rpm) used five times for 5 seconds with a 25-second break on ice in between. The lysate was centrifuged at 50,830 *g* for 18 min at 4 °C. The supernatant was discarded, and the pellet was homogenized in membrane buffer (50 mM Tris, 1 mM EDTA, 5 mM $MgCl_2$, 100 µg/mL bacitracin, 5 µg/mL soybean trypsin inhibitor, pH 7.4) with a glass-teflon homogenizer. Aliquots of 250 µL were shock frozen in liquid nitrogen and directly stored at -80 °C. Finally, the protein concentration was determined using the Lowry method.[20]

*cDNA constructs:* The CCR7-Nluc (CCR7_Nluc and CCR7_GSSG_Nluc) fusion constructs in pcDNA3.1 were generated as previously described,[19] using the Gibson Assembly (New England Biolabs) method.[21] Therefore, the sequences of the Nluc enzyme (pNLF1-C, Promega), (3xHA)-tagged CCR7 (CCR7, cdna.org, #CCR070TN00) were amplified by polymerase chain reaction and were directly fused in frame with different linker sequences (no linker and GSSG). DNA sequencing was performed to verify sequence integrity (Eurofins Genomics). Plasmids were cloned into *E.coli* DH5-alpha (New England Biolabs) and purified using a Maxiprep DNA purification kit (Invitrogen). The CCR7_GSSG_Nluc construct was already published.[10]

*ELISA:* ELISA-based experiments were performed as previously reported.[19] For confirmation of CCR7_Nluc and CCR7_GSSG_Nluc expression, HEK293T cells were transfected with the plasmid encoding 3xHA-CCR7, 3xHA-tagged CCR7_Nluc or 3xHA-tagged CCR7_GSSG_Nluc construct using polyethylenimine in suspension. Therefore, HEK293T cells were detached from their culture plates and diluted to a density of 3 x $10^5$ cells/mL in growth medium. This cell suspension was mixed with the preformed transfection mix (PEI/DNA ratio 2.5:1) consisting of 1.2 µg of receptor cDNA plasmid and 1.2 µg of single stranded salmon sperm DNA (ssDNA, Sigma Aldrich) in phosphate buffered saline (PBS) per 2.4 mL of cell suspension. Subsequently, cells were transferred to a 48-well plate (7.5 x $10^4$ cells/well), which was pretreated with poly-D-lysine (0.1 mg/mL, dissolved in water). Cells were incubated for 48 h at 37 °C and 5% $CO_2$. On the day of the assay, the medium was removed, and cells were incubated with 200 µL/well of ROTI®Histofix 4% fixation solution (Carl Roth) for 10 min at room temperature. Cells were washed once with 300 µL washing buffer for two minutes (150 mM NaCl, 25 mM Tris, pH 7.5) and blocked for one hour using 800 µL blocking buffer (30 g/L skim milk powder in washing buffer). After removal of the blocking solution, 200 µL/well of anti-HA rabbit IgG antibody (Sigma Aldrich, catalog # H6908, 1:4,000 in blocking solution) were added. After 60 min of incubation, wells were washed twice for two minutes (300 µL/well) and blocked again for one hour at room temperature, before 200 µL/well anti-rabbit IgG-HRP antibody (Invitrogen by Thermo Fisher Scientific, catalog # G-21234, 1:1,000 in blocking solution) was added. After incubation for one hour, cells were washed three times for two minutes (300 µL/well), before the substrate reaction was initiated by the addition of substrate buffer (6 mM o-phenylenediamine in 35 mM citric acid, 66 mM $Na_2HPO_4$, pH 5.0). After 15 minutes incubation in the dark, the reactions were terminated by addition of 1 M $H_2SO_4$ (200 µL/well). For each well, 2 x 150 µL of the resulting mixture were transferred to a clear, flat bottom 96-well plate and absorption was measured at 492 nm in a microplate reader. The measured absorbance values were baseline-corrected using cells transfected with a non-tagged muscarinic receptor (M3R, cdna.org) as negative control. These baseline-corrected values were normalized to 3xHA-CCR7 expression.

*Emission and excitation spectra of the fluorescent ligand:* For the detection of the emission and excitation spectra of the fluorescent ligand, we referred to a previously published procedure.[7,8] For the excitation spectrum, the fluorescent ligand was diluted to 500 µM in aqueous solution and 25 µL of this solutions were pipetted into a 384-well plate. Then, the excitation spectrum was measured with a CLARIOstar microplate reader. The emission spectrum of the fluorescent ligand (1 mM in DMSO) was recorded using a CLARIOstar (BMG Labtech, Ortenberg, Germany) microplate reader and 480 nm as excitation wavelength.

*Emission spectra of Nluc-labeled CCR7 proteins:* Furimazine (Promega, Mannheim, Germany 1:2,000) was added to the membrane preparations (5 µg protein/well). After 5 minutes incubation in the dark, the emission spectra were measured ranging from 350 to 700 nm using a CLARIOstar (BMG Labtech, Ortenberg, Germany) microplate reader. The emission spectra of CCR7_GSSG_Nluc has already been reported by Huber *et al.*[10,11]

*NanoBRET binding assays: Membrane-based NanoBRET saturation assay:* For the establishment of our NanoBRET binding assay, we referred to recently published protocols.[10,11] The fluorescent ligand Mz437 (**4**) was dissolved in DMSO (1 mM) and further diluted to varying concentrations in assay buffer (50 mM $Na_2HPO_4$, 50 mM $KH_2PO_4$, pH 7.4, 1 mg/mL saponin, 5% FBS) and 5 µL of these dilutions were pipetted to a 384-well plate. To determine total binding, 5 µL of assay buffer were added to the corresponding wells, while 5 µL of a solution of the non-fluorescent SLW131 (**10**, final assay concentration: 15 µM) in assay buffer were used to determine non-specific binding. Then, 20 µL of the membrane preparation (CCR7_GSSG_Nluc) diluted in assay buffer (3 µg total protein/well) were added and the plates were incubated for 90 min at 37 °C.

Subsequently, 5 µL of a furimazine solution (Promega, Mannheim, Germany, final assay dilution: 1:5,000) were added to each well (final assay volume: 35 µL) before measuring luminescence with a CLARIOstar microplate reader using 620/10 nm and 475/30 nm emission filters after 5 min of incubation in the dark. Bioluminescence resonance energy transfer (BRET) was determined as the ratio of acceptor fluorescence and donor luminescence. The algorithms for one-site saturation binding from PRISM10.2.1 (GraphPad, USA) were utilized to analyze total, non-specific and specific binding. Specific binding signals were calculated as a difference of total and non-specific binding. If required, netBRET values were calculated as the difference between total BRET values and the values obtained in the absence of a fluorescent ligand.

*Membrane-based NanoBRET competition assay:* The fluorescent ligand Mz437 (**4**) was dissolved in assay buffer and 5 µL of this solution (final assay concentration: 500 nM) were pipetted to a 384-well plate, followed by the addition of 5 µL of varying dilutions of the competing ligand dissolved in assay buffer. Then, 20 µL of the membrane preparations (CCR7_GSSG_Nluc diluted in assay buffer, 2.5 µg total protein/well) were added and the plates were incubated for 90 min at 37 °C. Subsequently, 5 µL of a furimazine solution (final assay dilution: 1:5,000 in assay buffer) were added to each well (final assay volume: 35 µL). Plates were read on a CLARIOstar microplate reader using 620/10 nm and 475/30 nm emission filters after 5 min of incubation in the dark. To determine the inhibition constants ($K_i$) of the non-labeled ligands, data were analyzed using the one site-fit $K_i$ equation in PRISM10.2.1 (GraphPad, USA). For compounds that showed more than 50% competition at the highest concentration tested, we manually set a constraint for the curve fitting to approach the value detected for non-specific binding (0% specific BRET). For compounds that showed less than 50% competition at the highest competitor concentration tested, only the values for percentual inhibition of tracer binding at a given concentration are provided.

*Membrane-based NanoBRET association kinetic assay:* 5 µL of a solution of the fluorescent ligand Mz437 (**4**) diluted to varying concentrations (final assay concentrations: 100–1000 nM) in assay buffer and 5 µL of assay buffer were transferred to a 384-well plate. For the determination of non-specific binding, we added a solution of a non-labeled competitor SLW131 (**10**) dissolved in assay buffer (final assay concentration: 10 µM) instead. After the addition of 5 µL of a furimazine solution (final assay dilution: 1:630 in assay buffer), plates were incubated for 3 minutes in the dark at ambient temperature. Subsequently, 20 µL of the membrane preparations (CCR7_GSSG_Nluc, 4 µg total protein/well) were added (final assay volume: 35 µL). BRET ratios were measured with a CLARIOstar microplate reader using 620/10 nm and 475/30 nm emission filters over time at ambient temperature. The obtained data were analyzed using the association kinetics (one ligand concentration) algorithm in PRISM10.2.1 (GraphPad, USA) to determine association kinetics using a pre-determined $k_{off}$ as a constraint.

*Membrane-based NanoBRET dissociation kinetic assay:* 5 µL of a solution of the fluorescent ligand Mz437 (**4**) diluted to varying concentrations (final assay concentrations: 250–1000 nM) in assay buffer and 5 µL of assay buffer were transferred to a 384-well plate. In order to determine non-specific binding, we added a solution of a non-labeled competitor SLW131 (**10**) dissolved in assay buffer (final assay concentration: 10 µM) instead of the 5 µL of assay buffer. Then, 20 µL of the membrane preparation (4 µg total protein/well) were added, and plates were incubated for 1.5-2 h at ambient temperature in the dark. Subsequently, 5 µL of a furimazine solution (final assay dilution: 1:630 in assay buffer) were added. Plates were incubated for further 5 minutes in the dark at ambient temperature. Thereafter, 1 µL of a solution of the unlabeled competitor SLW131 (**10**) dissolved in assay buffer (final assay concentration: 10µM) was added. For control experiments, we added 1 µL of assay buffer instead of the competitor solution. BRET ratios were measured with a CLARIOstar microplate reader using 620/10 nm and 475/30 nm emission filters over time at ambient temperature. Specific BRET ratios were calculated as a difference of total and non-specific binding. The obtained data were analyzed using the dissociation - one phase exponential decay algorithm in PRISM10.2.1 (GraphPad, USA) to determine dissociation kinetics.

*Live cell NanoBRET:* HEK293T cells were transfected with 5.5 µg of the plasmid (CCR7_GSSG_Nluc) using polyethylenimine (PEI; 7.5 µg) as transfection reagent. After 24 h at 37 °C and 5% $CO_2$, the cells were detached with DMEM and transferred to a white F-bottom assay 384-well plate [10,000 cells/well], which was coated with poly-D-lysine (0.1 mg/mL, dissolved in water), and incubated for further 24 h at 37 °C and 5% $CO_2$. Subsequently, cells were washed with phosphate-buffered saline (Gibco™ DPBS, with $CaCl_2$ and $MgCl_2$). Assay medium (Gibco™ DMEM/F-12, 15 mM HEPES, no phenol red supplemented with 5% FBS) was added and cells were incubated at 37 °C for 30 minutes. Then, 5 µL of a solution containing the fluorescent ligand Mz437 (**4**), diluted in assay medium at varying concentrations, were added in case of saturation binding experiments. To determine non-specific binding, 5 µL of a solution of the unlabeled competitor SLW131 (**10**) dissolved in assay medium (final assay concentration: 10 µM) were added. For competition binding experiments, 5 µL of a solution of the fluorescent ligand (**4**) diluted in assay medium (final assay concentration: 500 nM) and 5 µL of a solution of the potential competitor (diluted from 10 mM DMSO-stock solutions with assay medium) at varying concentrations were added to the corresponding wells. After 90 min of incubation at 37 °C, 5 µL of a furimazine solution (final assay dilution: 1:2,500 – 1:2,700, diluted with assay medium) were added. At the 96-well plates the double volume was added. After a further incubation of 5 min in the dark at 37 °C, BRET ratios were measured with a CLARIOstar microplate reader using 620/10 nm and 475/30 nm emission filters. Total, non-specific and specific binding, which was calculated as a difference of total and non-specific binding, were analyzed using the algorithms for one-site saturation binding in PRISM10.2.1 (GraphPad, USA). To determine the inhibition constants ($K_i$) of the potential competitors, data were normalized to total and non-specific binding and analyzed using the one site-fit $K_i$ equation in PRISM10.2.1 (GraphPad, USA).

**Biology**

**Cell culture.** HEK293T cells (obtained from the American Type Culture Collection) were grown in Dulbecco's Modified Eagle Medium (DMEM, Thermo Fisher Scientific) supplemented with 10% fetal bovine serum (FBS, PANbiotech), 100 units/ml penicillin and 100 µg/ml streptomycin at 37 °C and 5% $CO_2$. For culture of stable hCCR7-HEK293 DMEM was supplemented with G418 (500 mg/ml) (InvivoGen). For generation of single clones, cells were detached and seeded into a 96-well plate at a density of 1 cell per well in complete culture media. Cells that grew to confluency were detached and expanded stepwise.

**β-arrestin2 NanoBiT assay.** HEK293T cells were transfected with 500 ng hCCR7-smBiT or SS-HA-mCCR7-smBiT, 50 ng LgBiT-β-arrestin2 mutant (R393E, R395E) (kindly provided by Asuka Inoue) and 4450 ng empty pcDNA3.1 per 3 x $10^6$ cells. At 28 h post-transfection arrestin recruitment was measured using NanoBiT technology. Briefly, cells were counted, resuspended in

HBSS with 20 mM HEPES and transferred to a white 96-well plate with 60,000 cells per well. After preincubation of cells with antagonists or vehicle (final DMSO concentration 0.1%) for 30 min, cells were loaded with Nano-Glo Nano luciferase substrate (Promega, final dilution 1:1000) for 3 min, baseline luminescence was recorded and cells were stimulated with the indicated CCL19 concentration. Luminescence signals were measured using a PHERAstar FSX microplate reader (BMG Labtech). Luminescence changes were baseline- and buffer-corrected; concentration-inhibition-curves were derived from the maximum luminescence responses and expressed as percentage of the indicated concentration of CCL19.

**Real-time BRET-based G protein activation assay.** HEK293T cells were transfected in suspension with 1500 ng hCCR7 or 1500 ng SS-HA-mCCR7, 800 ng Gαo, 400 ng Venus(156-239)-Gβ1, 400 ng Venus(1-155)-Gγ2 and 400 ng masGRK3ct-Nluc (BRET sensor plasmids were kindly provided by Kirill A. Martemyanov) using polyethylenimine (PEI, 25kDa, linear, Polysciences, 1 mg/ml, DNA to PEI solution ratio 1:3). Empty pcDNA3.1 vector was used to adjust the total amount of DNA to 5000 ng per transfection of 3 x $10^6$ cells in a 10 cm dish. At 28 h post-transfection, BRET was measured according to previously published protocol.[22] Briefly, cells were counted, resuspended in Hank`s Balanced Salt Solution (HBSS) supplemented with 20 mM HEPES and transferred to a white 96-well plate with 80,000 cells per well. HEK293T cells were preincubated with antagonists or vehicle (final DMSO concentration 0.1%) for 30 min. Afterwards Nano-Glo nanoluciferase substrate (Promega, final dilution 1:1000) was added for 3 min before recording baseline BRET and stimulating cells with the indicated CCL19 concentration. BRET was recorded with a PHERAstar FSX microplate reader (BMG Labtech) and calculated as ratio of the fluorescence emitted by Venus-Gβγ (535 ± 30 nm) and the luminescence of GRK3-Nluc (475 ± 30 nm). The difference between the BRET ratio before and after agonist stimulation was calculated and buffer-corrected. Concentration-inhibition-curves were derived from the maximum BRET signals and expressed as percentage of the indicated concentration of CCL19.

**Label-free whole cell biosensing based on detection of dynamic mass redistribution (DMR).** For DMR experiments, stable hCCR7-HEK293 cells were seeded at a density of 18,000 cells per well into fibronectin-coated 384-well biosensor plates. Murine BMDCs were plated on 100 mm Petri dishes (non-adhesive plastic; Fisher; cat.nr: 10470613) at a density of 2 × $10^6$ cells/plate in complete medium (RPMI 1640 supplemented with 10% fetal calf serum, 2 mM L-glutamine, 100 U/ml penicillin, 100 µg/ml streptomycin, 50 µM ß-mercaptoethanol; all purchased from Thermo Fisher Scientific) containing 10% GM-CSF (supernatant from hybridoma culture). DC differentiation was induced with 200 ng/ml LPS from E.coli 0127:B8 (Sigma-Aldrich) for 24 – 48 h at 37 °C and 5% $CO_2$. The supernatant was then collected to harvest the mature non-adherent BMDCs, cells were washed with assay buffer (HBSS with 20 mM HEPES) and seeded at a density of 80,000 cells per well on 384-well fibronectin-coated biosensor plates. Cells were incubated for 60 min at 37 °C in the EPIC reader (EnSight Multimode Plate Reader, Perkin Elmer, MA, US), followed by a 3 min baseline read. Thereafter, SLW131 (**10**) and SLW132 (**21m**) were added using the Selma semiautomatic liquid handling system (Analytik Jena AG, Jena, DE) for another 60 min. After a second 3 min baseline read, CCL19 was added and wavelength shifts over time were recorded for 3600 seconds. Real-time DMR recordings were buffer-corrected and are presented as wavelength shift over time; concentration-effect-curves were derived from the peak wavelength shifts, concentration-inhibition-relationships are expressed as percentage of the indicated concentration of CCL19.

**Fluorescence microscopy**. Stably transfected hCCR7-HEK293 cells were seeded onto poly-D-lysine (PDL) - coated 8-well μ-slides (Ibidi) at a density of $1 \times 10^5$ cells per well and cultured overnight at 37 °C and 5% $CO_2$. For immunostaining, the cells were first fixed with 4% paraformaldehyde solution, washed with PBS, and then blocked for 60 minutes with 10% goat serum and 1% fatty acid-free bovine serum albumin (BSA) in PBS at 37 °C. Cells were then incubated with the primary antibody, anti-CCR7 mouse IgG (R&D Systems, catalog # 150503, 1:100 in blocking solution) for 60 minutes at 37 °C. Following intense washing with PBS, the secondary antibody, goat anti-mouse IgG (H+L) conjugated to FITC (Sigma-Aldrich, catalog # AP124F, 1:500 in blocking solution) was added for 60 minutes at 37 °C. After a 2nd round of intense washing with PBS, cells were counterstained with 4′,6-diamidin-2-phenylindole (DAPI) solution (0.1 μg/mL) for 15 min in the dark at room temperature. The following day, cells were treated with either cmp2105 (**1**), SLW131 (**10**), or SLW132 (**21m**) (at a final concentration of 10 μM each, or vehicle solution (OptiMEM reduced serum medium (Gibco) with 0.1% DMSO) for 60 min at 37 °C. Thereafter, the fluorescent CCR7 probe Mz437 (at a final concentration of 500 nM) was added for 60 min at 37 °C and fluorescence images were acquired. Microscopy was performed using an AxioObserver.Z1 microscope (Carl Zeiss, Jena, Germany) equipped with an ApoTome imaging system and a Heating unit XL S, utilizing a Plan-Apochromat ×63/1.40 Oil objective (filter set 43 (red), 38 (green), 49 (blue)). Image processing and line scan analysis were conducted using Zen blue Imaging software. Fluorescence intensity values were normalized to the vehicle control.

**$CD4^+$ T cell transwell migration assay.** Four wild-type C57BL/6 mice (Charles River Laboratories) were used to isolate naive $CD4^+$ T cells from their skin-draining lymph nodes and spleens using the EasySep™ Mouse Naive $CD4^+$ T Cell Isolation Kit (Stemcell). The T cells ($1 \times 10^6$ cells/ml) were incubated with 10 µM of SWL131, SWL132, and a DMSO control in chemotaxis buffer (RPMI + 0,5% BSA + 10 mM HEPES + 100 U/ml penicillin/streptomycin + 1 mM sodium pyruvate + 1x non-essential amino acid solution + 50 µM ß-mercaptoethanol) for 30 minutes. In the lower compartment of the transwell plate (Corning 3421), 600 µl of chemotaxis buffer were added, with and without 100 ng/ml CCL19 or 100 ng/ml CXCL12. A total of 100 µl of the cell suspension was then transferred directly into the upper compartment of a transwell insert. After a 2.5-hour incubation period, the migrated cells were quantified using flow cytometry on a BD LSRFortessa with the aid of counting beads.[23] The percentage of net migration was calculated by subtracting the number of cells that migrated to CCL19 or CXCL12 from the number of cells that migrated in the absence of chemokines, and then dividing the result by the number of cells in the input control. The input control consisted of 100 µl of cells added directly to the lower compartment.

**Dendritic cell (DC) culture**

Cultures were started from freshly isolated bone marrow of 8-12-week-old mice with C57BL/6J background. DC differentiation was induced by plating $2x10^6$ cells in 10 ml complete medium (Roswell Park Memorial Institute (RPMI) 1640 supplemented with 10% Fetal Calf Serum, 2 mM L-Glutamine, 100 U/ml Penicillin, 100 μg/ml Streptomycin, 50 μM ß-Mercaptoethanol) (all purchased from Thermo Fischer) containing 10% Granulocyte-Monocyte Colony Stimulating

Factor (GM-CSF, supernatant from hybridoma culture). Cells were fed on day 3 and 6 with complete medium supplemented with 20% GM-CSF. To induce maturation, cells were stimulated overnight with 200 ng/ml lipopolysaccharide (LPS) from *E.coli* 0127:B8 (Sigma) and used for experiments on day 8-9 (mature DCs). DC differentiation and maturation was confirmed by flow cytometric assessment of surface markers such as MHCII, CD11c, CD86 and CCR7. For 3D collagen migration experiments, mature cells were treated with 10 µM SLW131, SLW132 or DMSO (ctrl) for 1 h, casted into migration chambers and recorded by video-microscopy.

**Flow cytometry**
Before staining, 1-2x$10^6$ cells were incubated for 15 min at 4 °C with blocking buffer (1 x PBS, 1% BSA, 2 mM EDTA) containing 5 mg/ml anti-CD16/CD32 antibody (2.4G2; BD Biosciences). For cell surface staining, cells were incubated for 30 min at 4 °C with conjugated monoclonal antibodies diluted in blocking buffer. The following antibodies were used: mouse anti-mouse CCR7-PE (4B12, 1:300), rat anti-mouse I-A/I-E-eFluor450 (M5/114.15.2, 1:800), hamster anti-mouse CD11c-APC (N418, 1:300), anti-mouse CD86 PE (GL1, 1:500). For live dead staining the Live/Dead Fixable Dead Cell Stain Kit (1:1000, Invitrogen) or DRAQ7 (1:1000, Biolegend) was used. Flow cytometry was performed on a LSR flow cytometer (BD Biosciences). Data analysis was carried out using FlowJo X 10.0.7r2.

***In vitro* 3D collagen migration assay**
For 3D *in vitro* migration, 2x$10^5$ bone marrow-derived DCs were suspended in a medium-collagen I mixture (PureCol bovine collagen, (INAMED) in 1x minimum essential medium eagle (MEM, Invitrogen) and 0.4% sodium bicarbonate (Sigma) at a volume ratio of 1:2 yielding in a final collagen concentration of 1.73 mg/ml. Collagen gel mixtures were casted into custom-made migration chambers as previously described[19,24] and incubated for 45 min at 37 °C to allow polymerization of the gel. CCL19 was suspended in full medium to a final concentration 100 nM and placed on top of the gel. To prevent drying-out of the gels, migration chambers were sealed with Paraffin wax (Sigma-Aldrich). Gels that failed to polymerize were excluded from the analysis.
Image acquisition was performed with a Nikon Eclipse widefield microscope and a C-Apochromat 10×/0.3 PH1 air objective. Images were acquired in 120 s intervals for 2 hrs at 37 °C, 5% $CO_2$. 50 cells were tracked manually, using the ‘Manual tracking Plug-in’ for ImageJ. The ImageJ Chemotaxis tool was used to determine average (frame-to-frame) speed and persistence distance in gradient direction/total distance). Additionally, automized y-displacement over time was measured using an in-house generated MATLAB script as recently described.[19,24]

NMR SPECTRA OF SYNTHESIZED COMPOUNDS

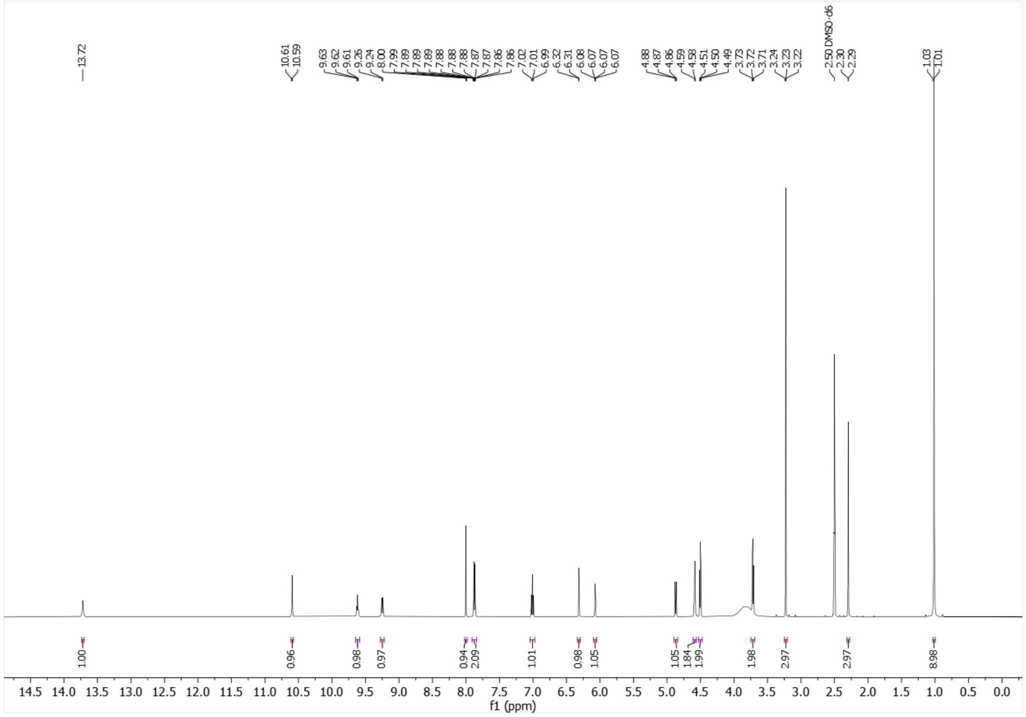


[1]H-NMR (500 MHz, DMSO-$d_6$) of **3**.

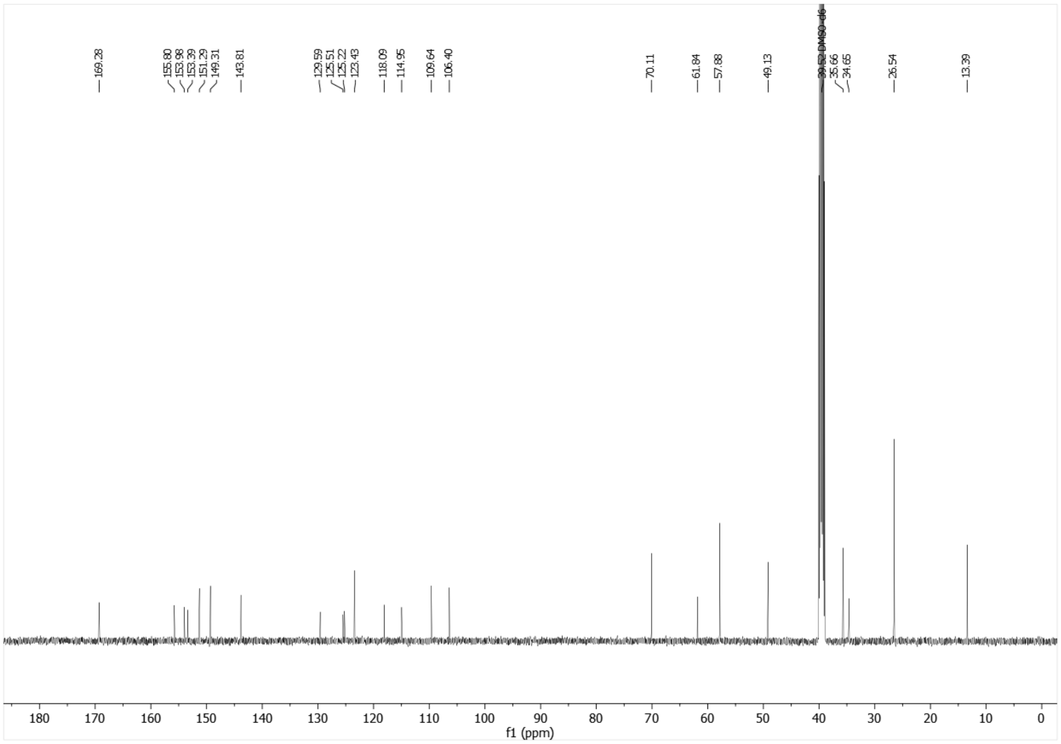


[13]C-NMR (126 MHz, DMSO-$d_6$) of **3**.

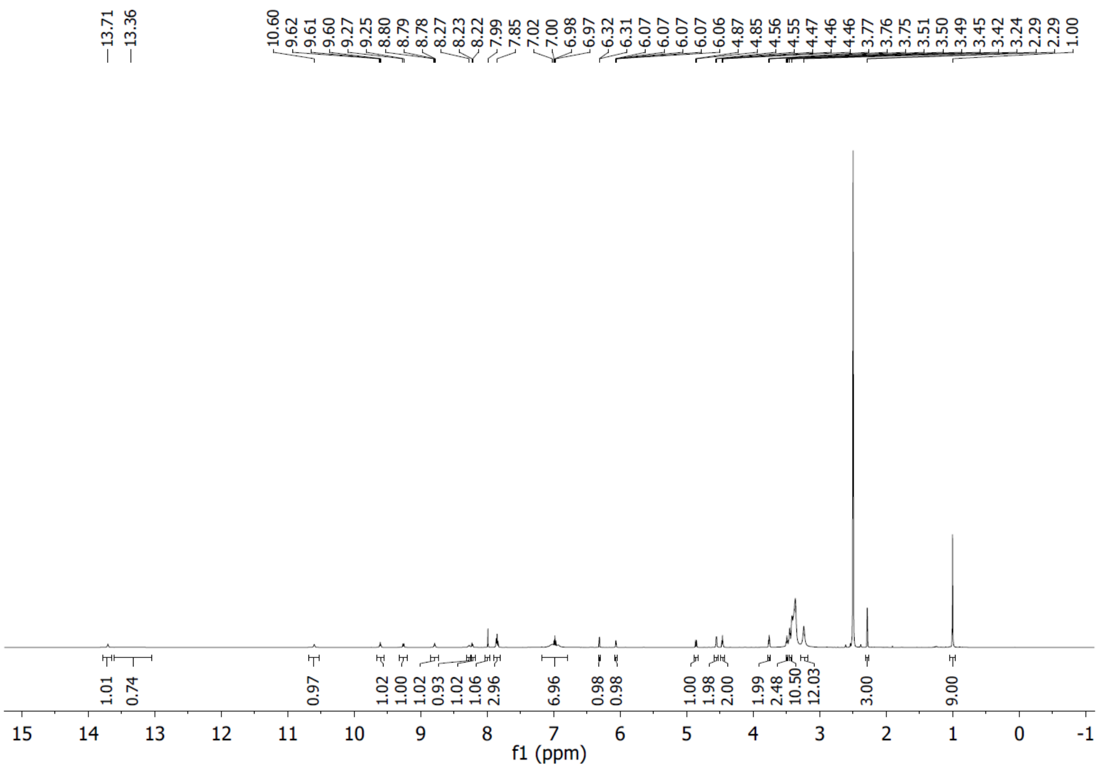

[1]H-NMR (600 MHz, DMSO-$d_6$) of Mz437 (**4**).

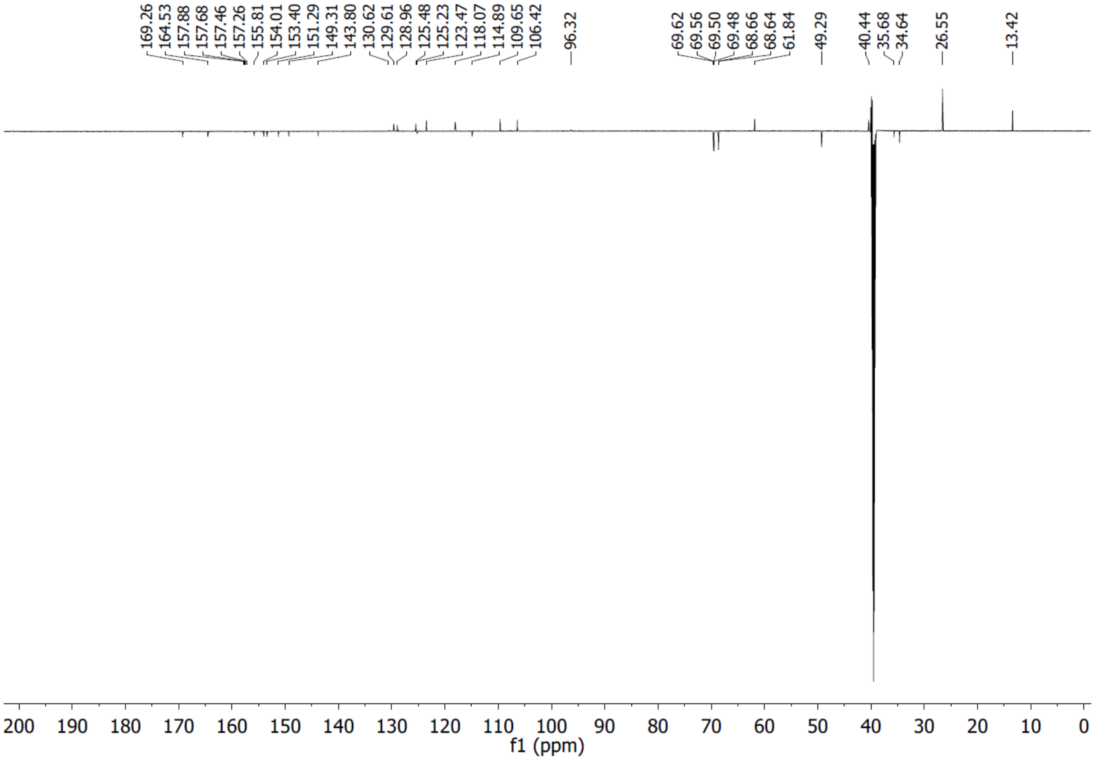

[13]C-NMR (151 MHz, DMSO-$d_6$) of Mz437 (**4**).

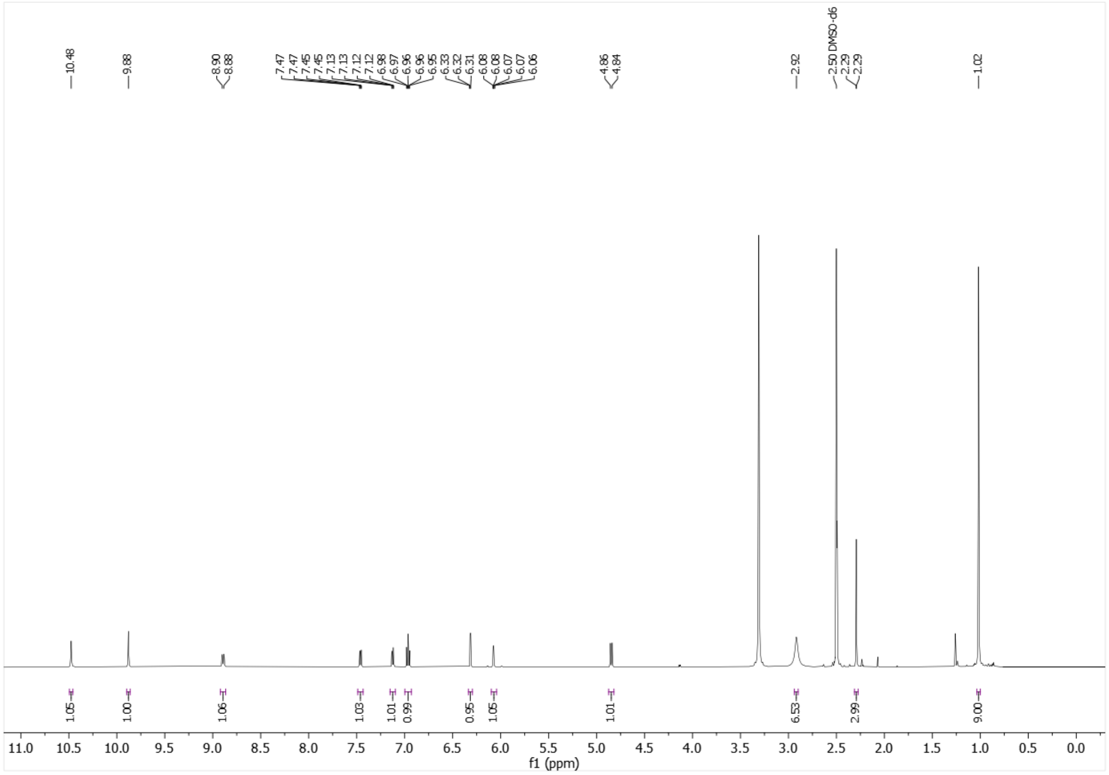


[1]H-NMR (500 MHz, DMSO-$d_6$) of SLW131 (**10**).

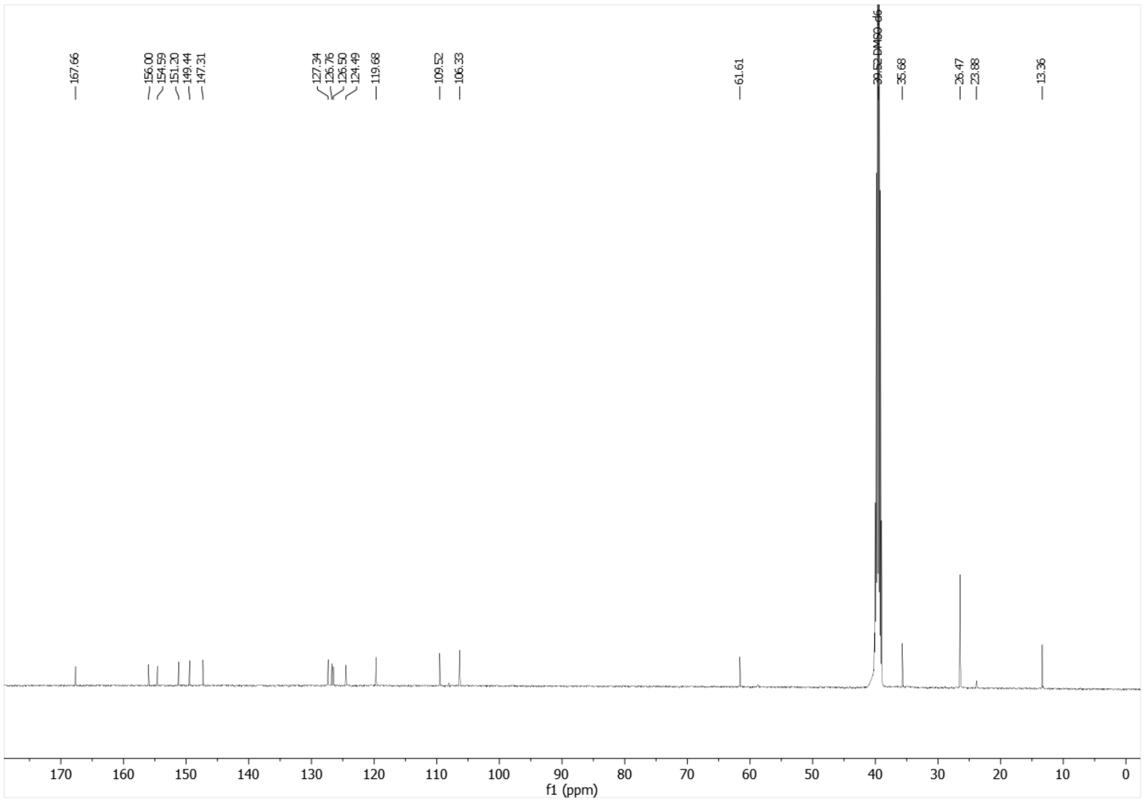


[13]C-NMR (126 MHz, DMSO-$d_6$) of SLW 131 (**10**).

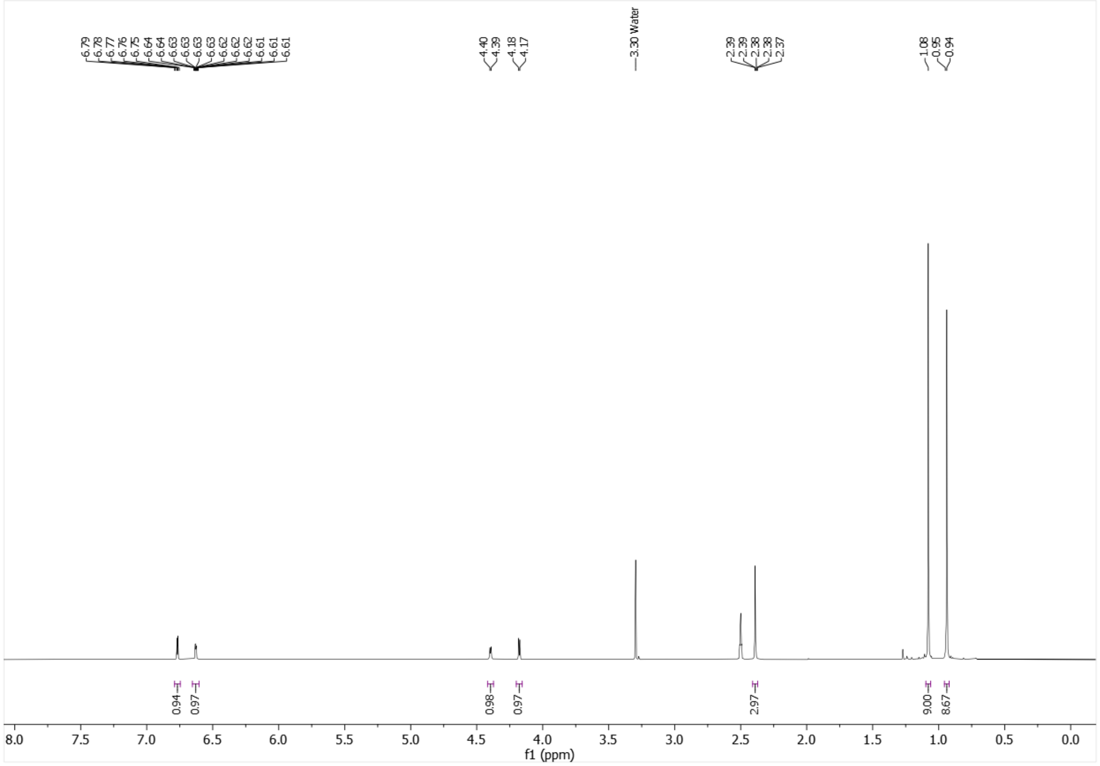


[1]H-NMR (500 MHz, DMSO-$d_6$) of **14d**.

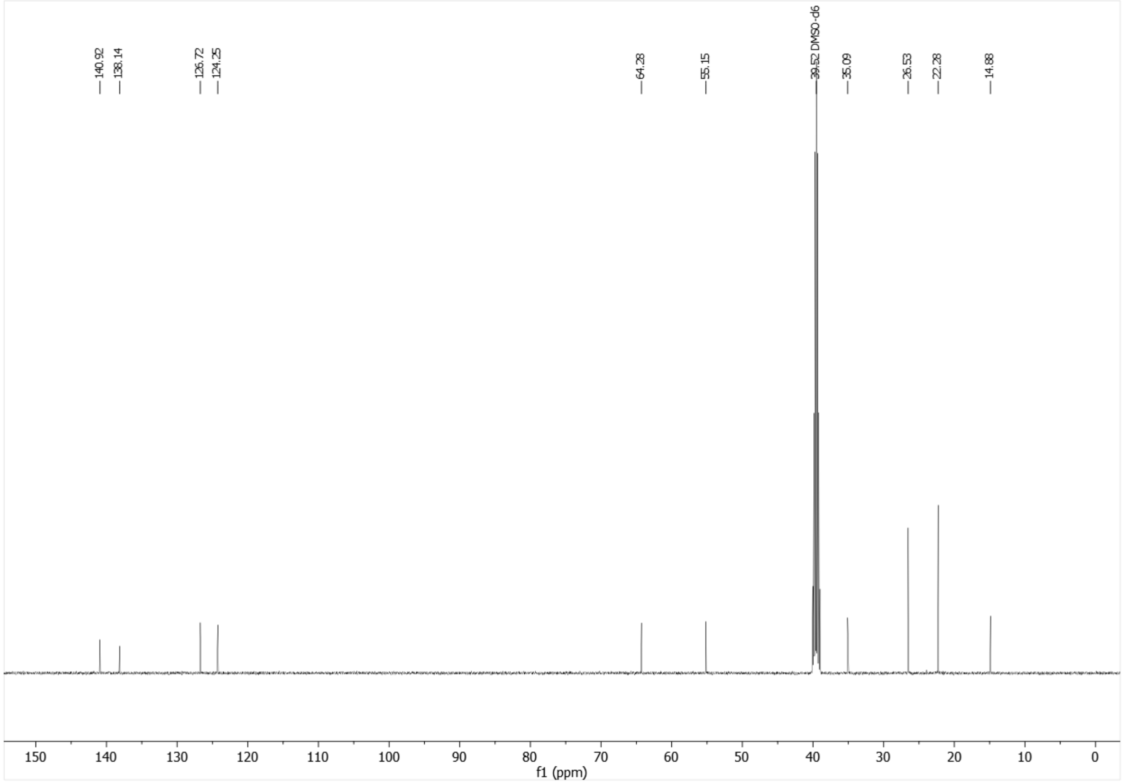


[13]C-NMR (126 MHz, DMSO-$d_6$) of **14d**.

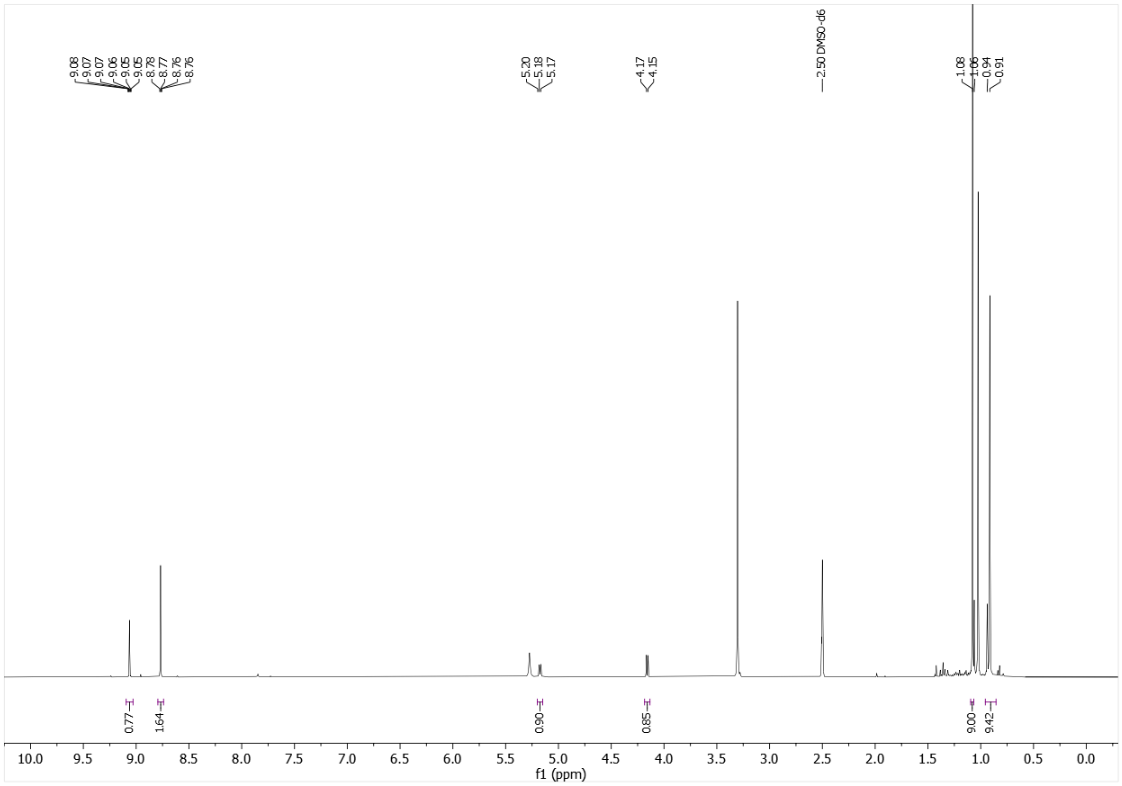

$^{1}$H-NMR (500 MHz, DMSO-$d_6$) of **14e**.

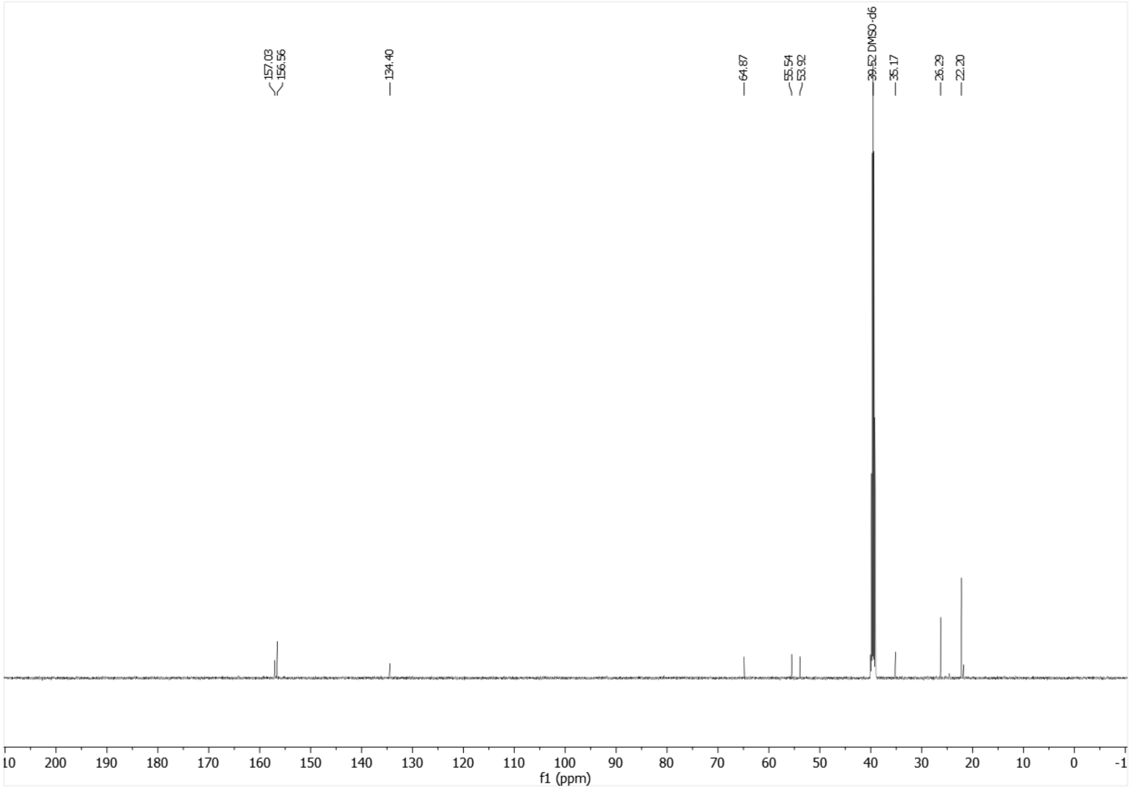

$^{13}$C-NMR (126 MHz, DMSO-$d_6$) of **14e**.

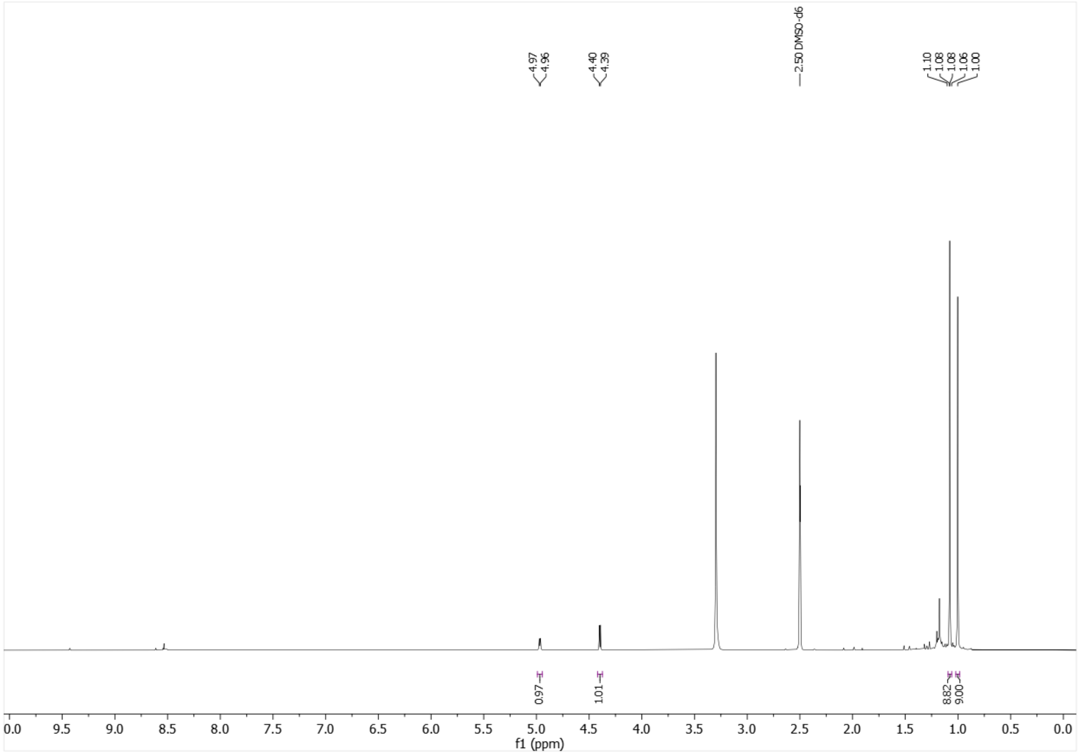


$^{1}$H-NMR (500 MHz, DMSO-$d_6$) of **14f**.

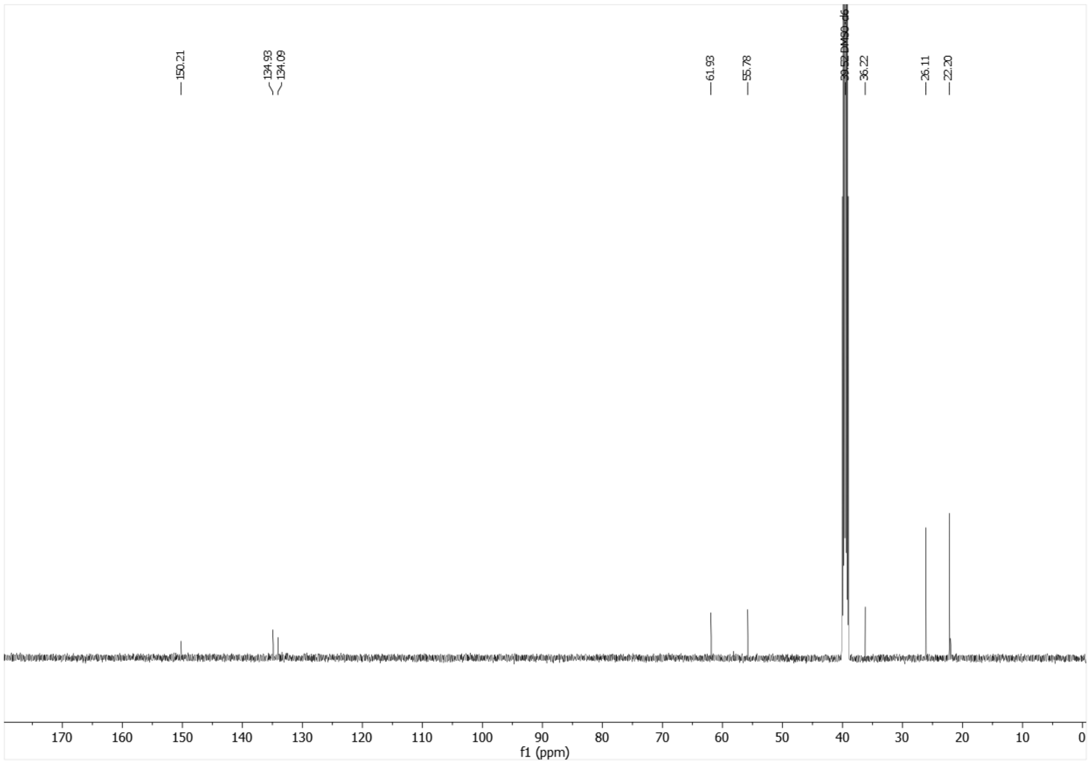


$^{13}$C-NMR (126 MHz, DMSO-$d_6$) of **14f**.

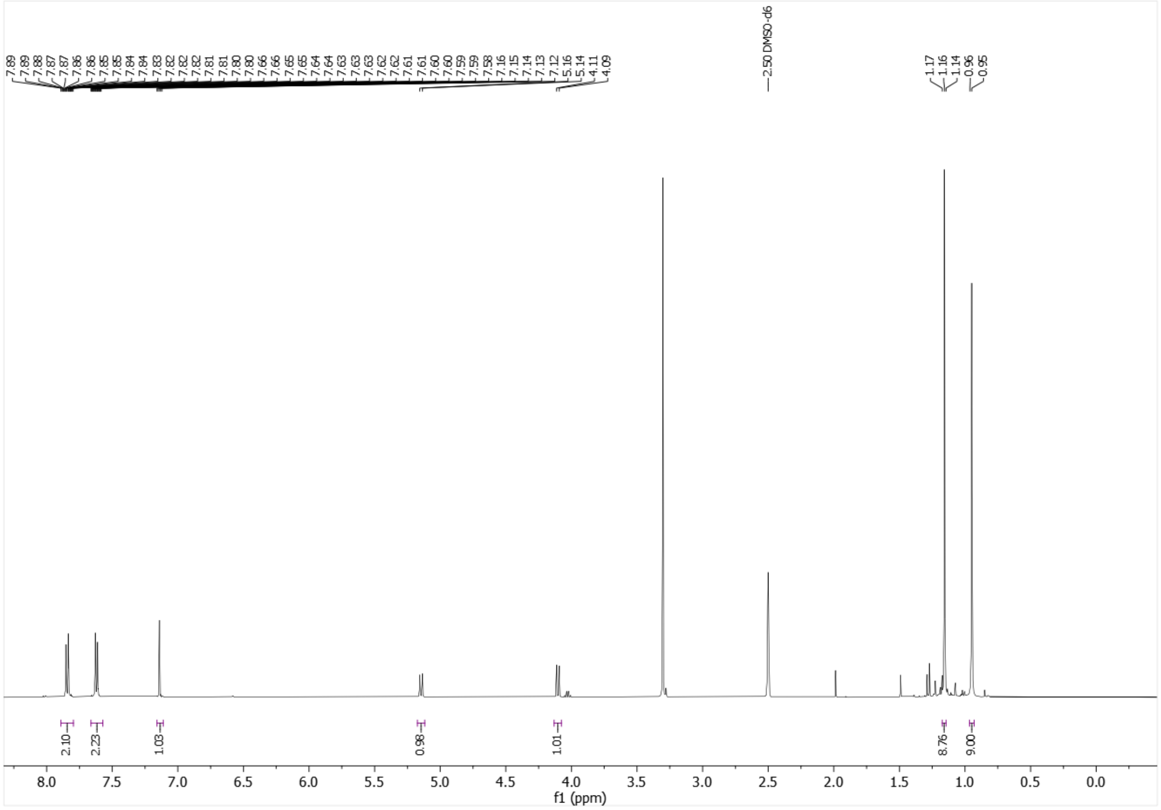

$^{1}$H-NMR (500 MHz, DMSO-$d_6$) of **14g**.

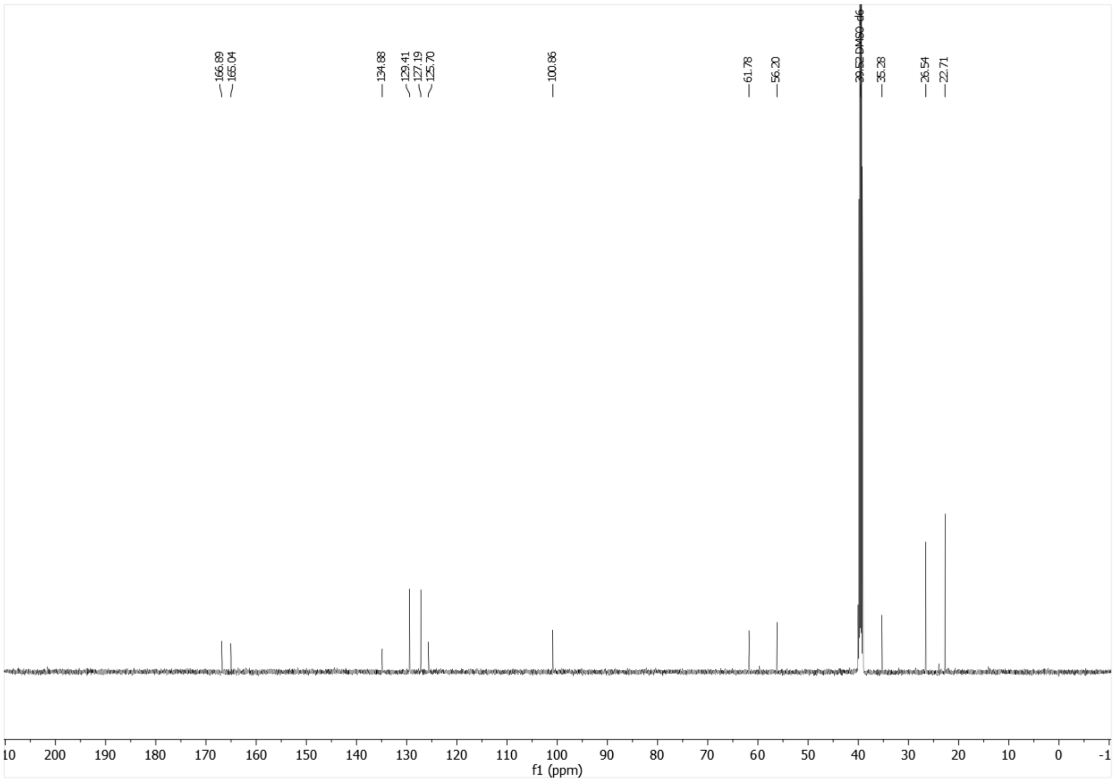

$^{13}$C-NMR (126 MHz, DMSO-$d_6$) of **14g**.

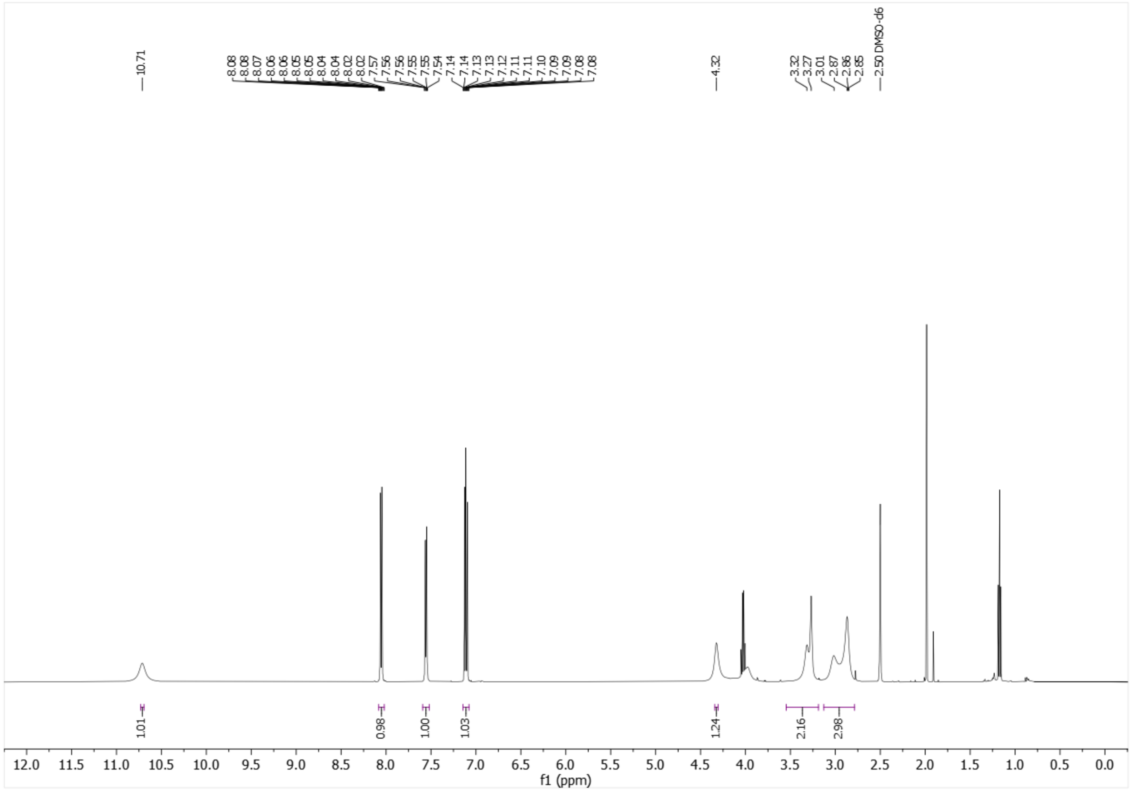

[1]H-NMR (500 MHz, DMSO-$d_6$) of **16c**.

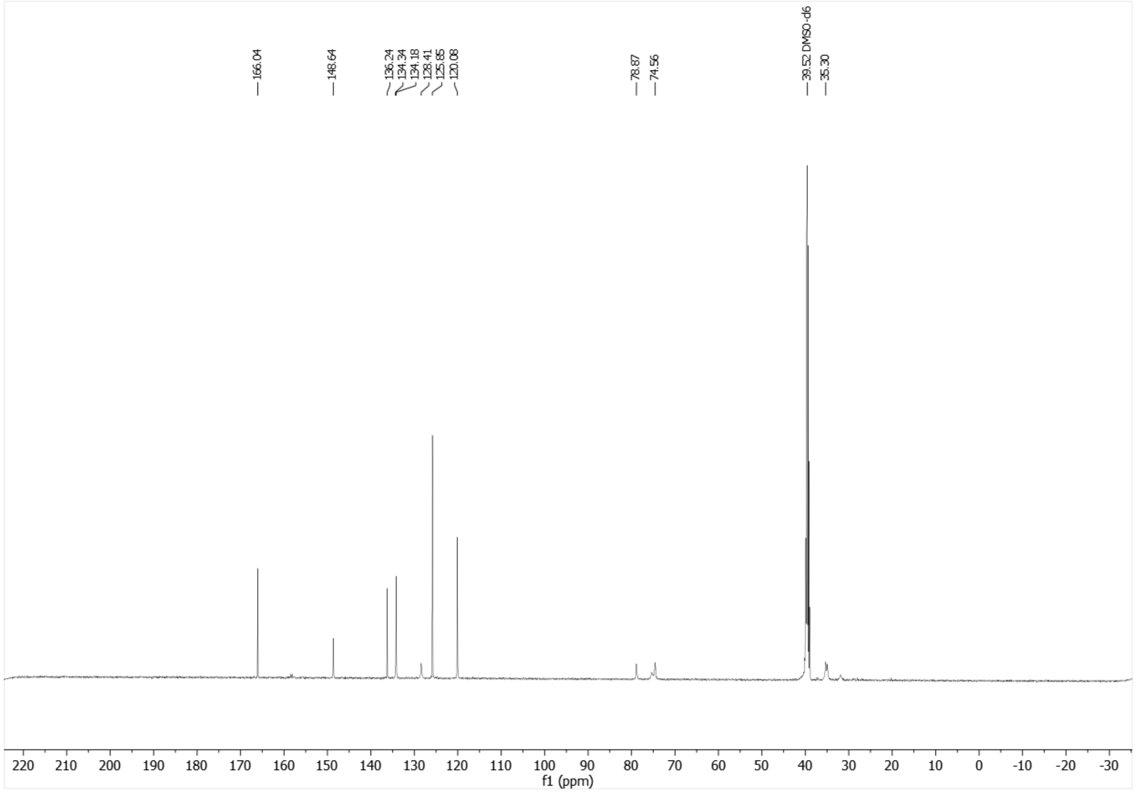

[13]C-NMR (126 MHz, DMSO-$d_6$) of **16c**.

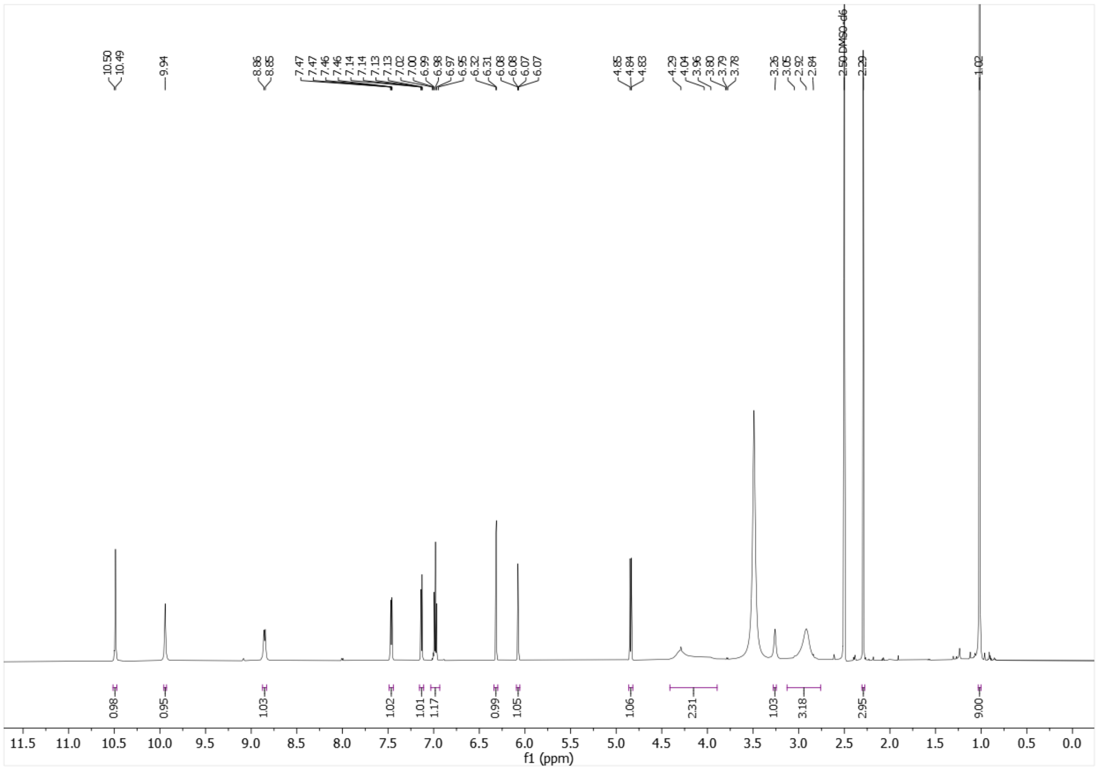

$^{1}$H-NMR (500 MHz, DMSO-$d_6$) of **20a**.

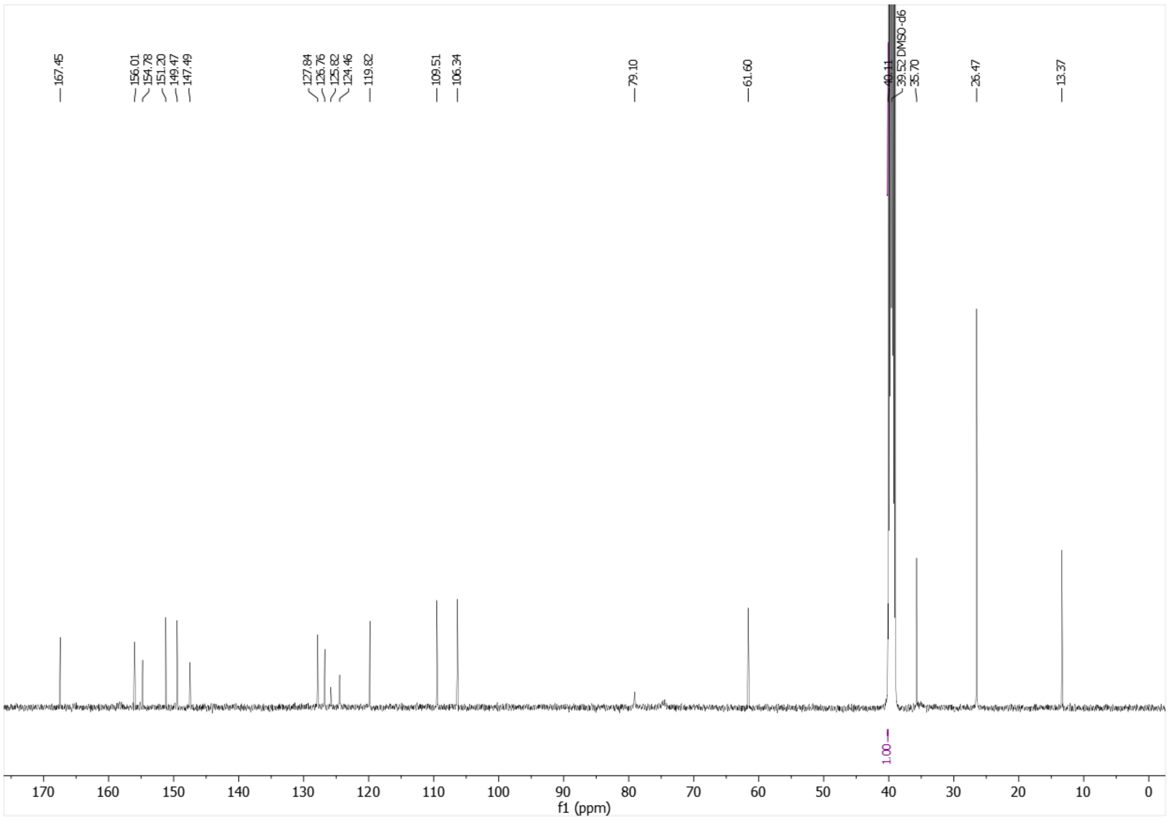

$^{13}$C-NMR (126 MHz, DMSO-$d_6$) of **20a**.

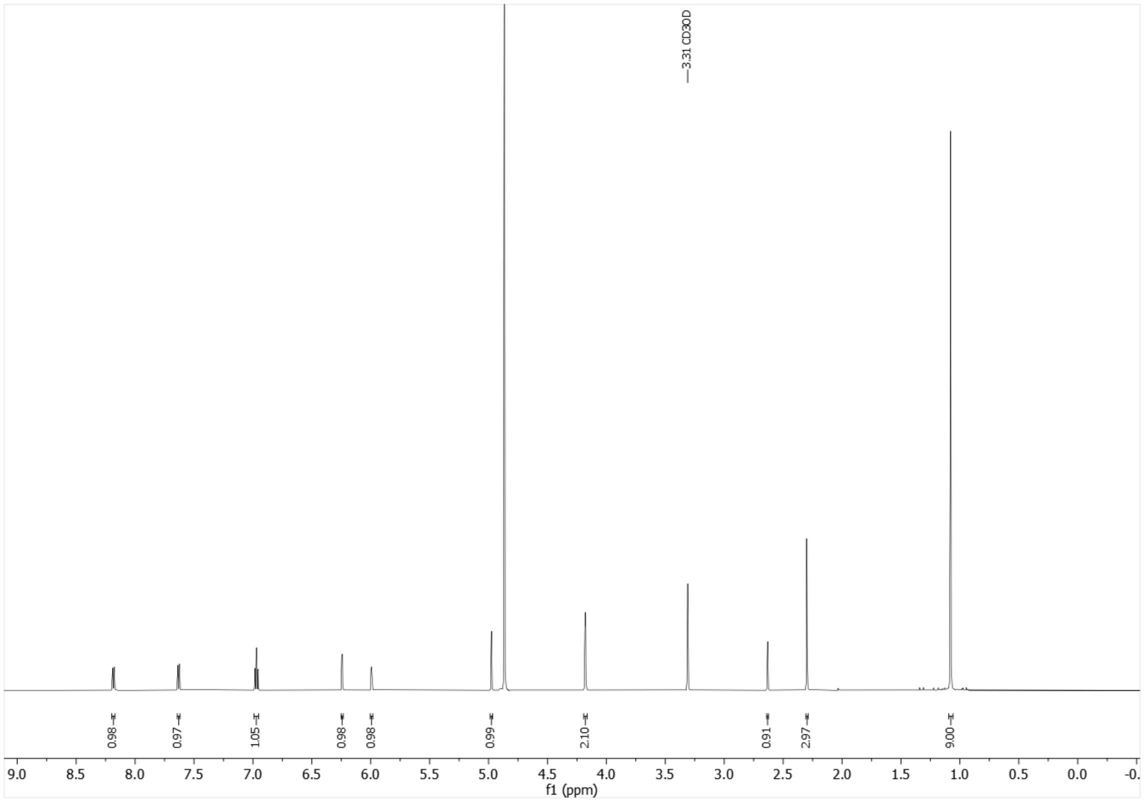


$^{1}$H-NMR (600 MHz, MeOD-$d_4$) of **20b**.

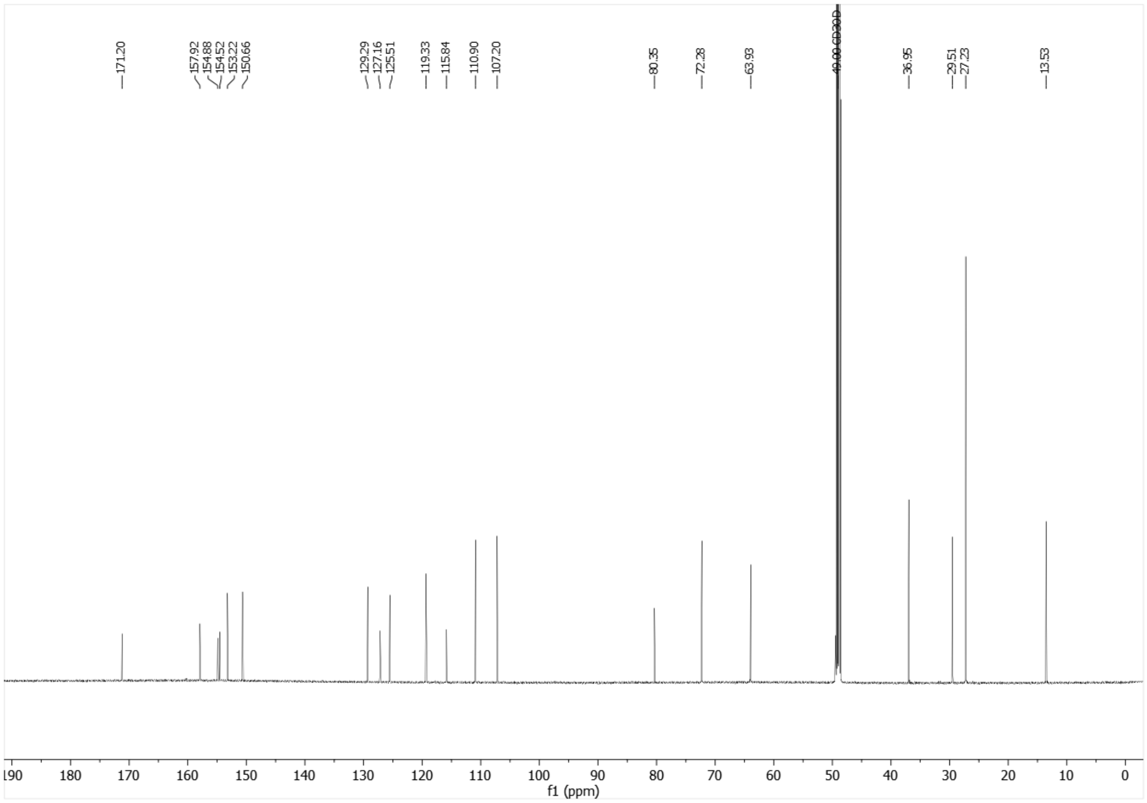


$^{13}$C-NMR (151 MHz, MeOD-$d$) of **20b**.

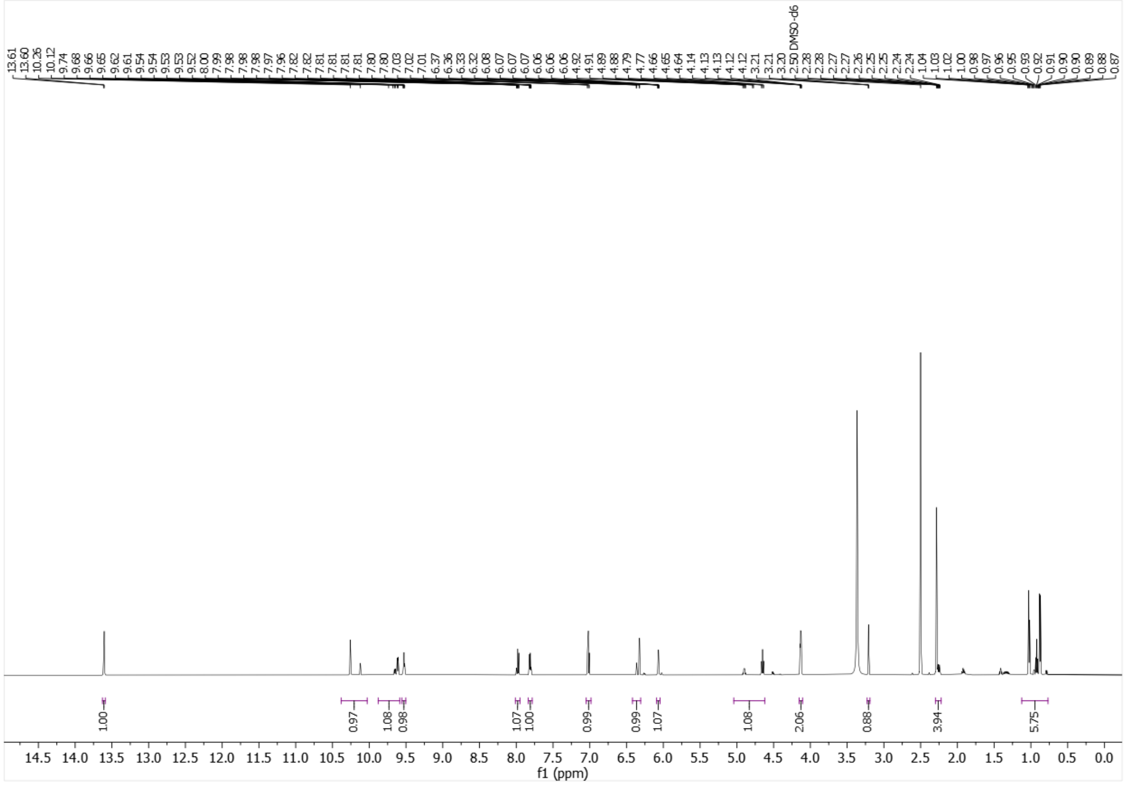


[1]H-NMR (600 MHz, DMSO-$d_6$) of **20c**.

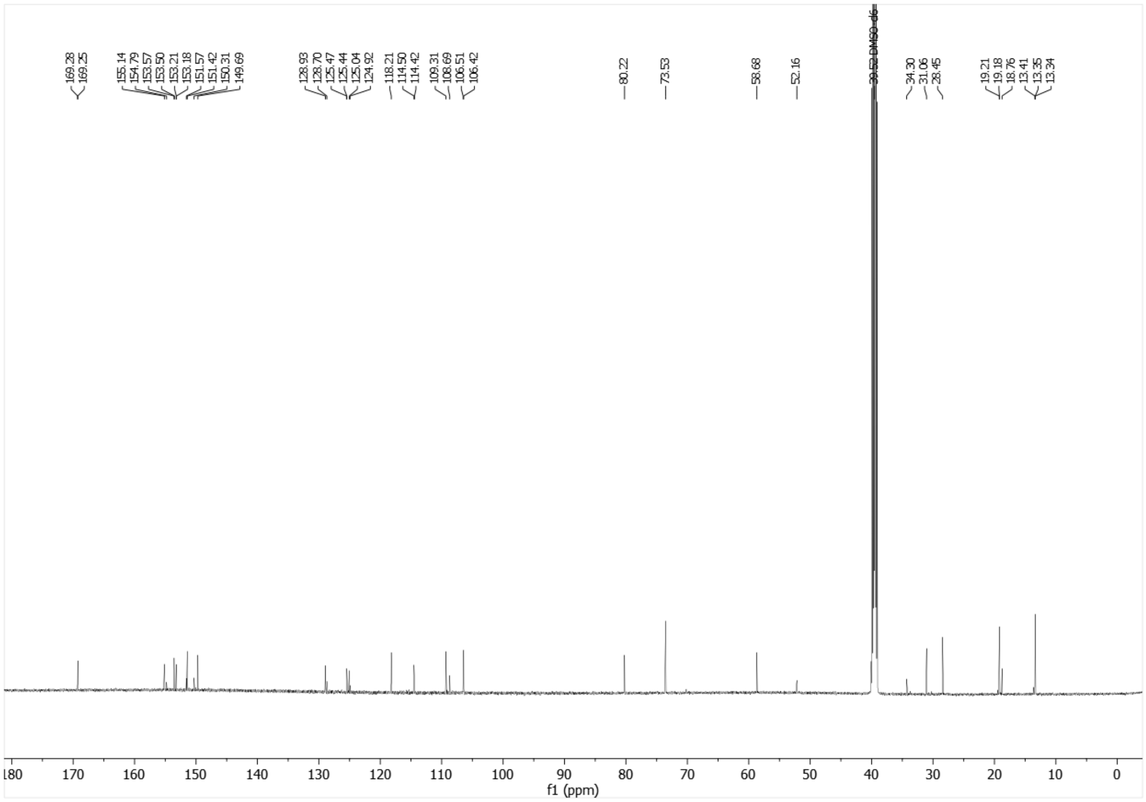


[13]C-NMR (151 MHz, DMSO-$d_6$) of **20c**.

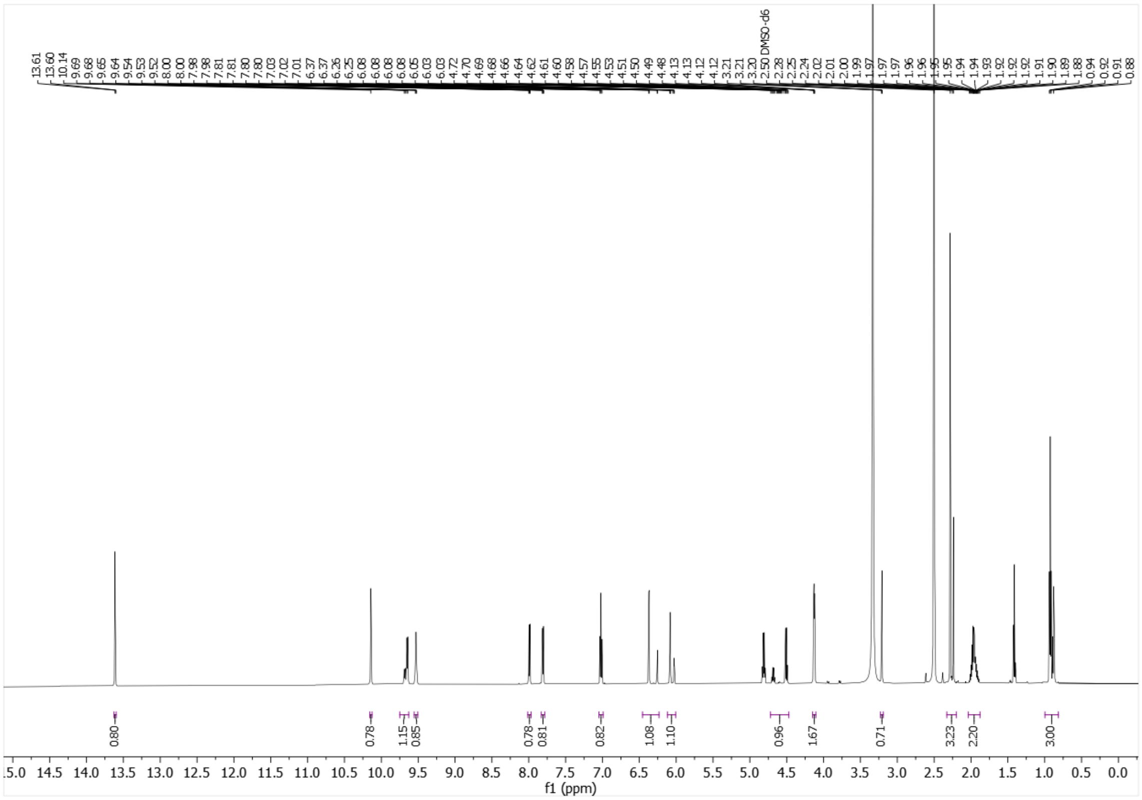

$^{1}$H-NMR (600 MHz, DMSO-$d_6$) of **20d**.

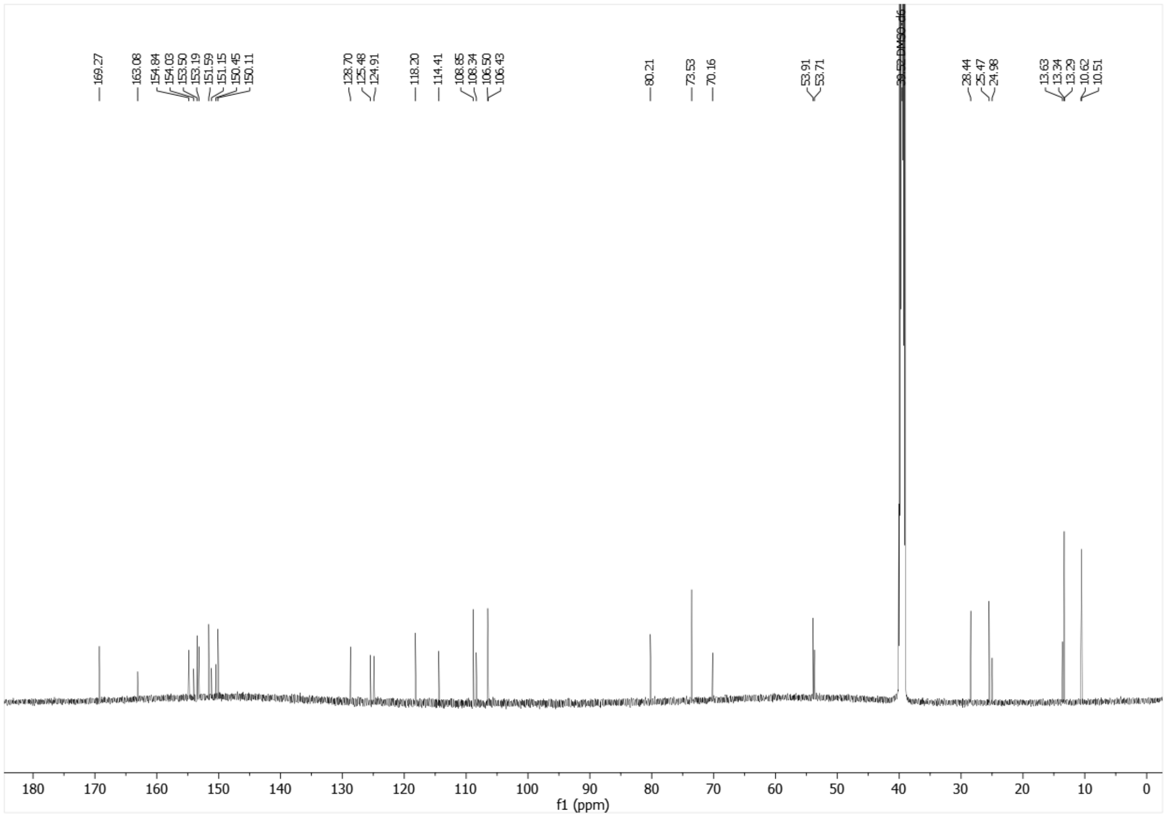

$^{13}$C-NMR (151 MHz, DMSO-$d_6$) of **20d**.

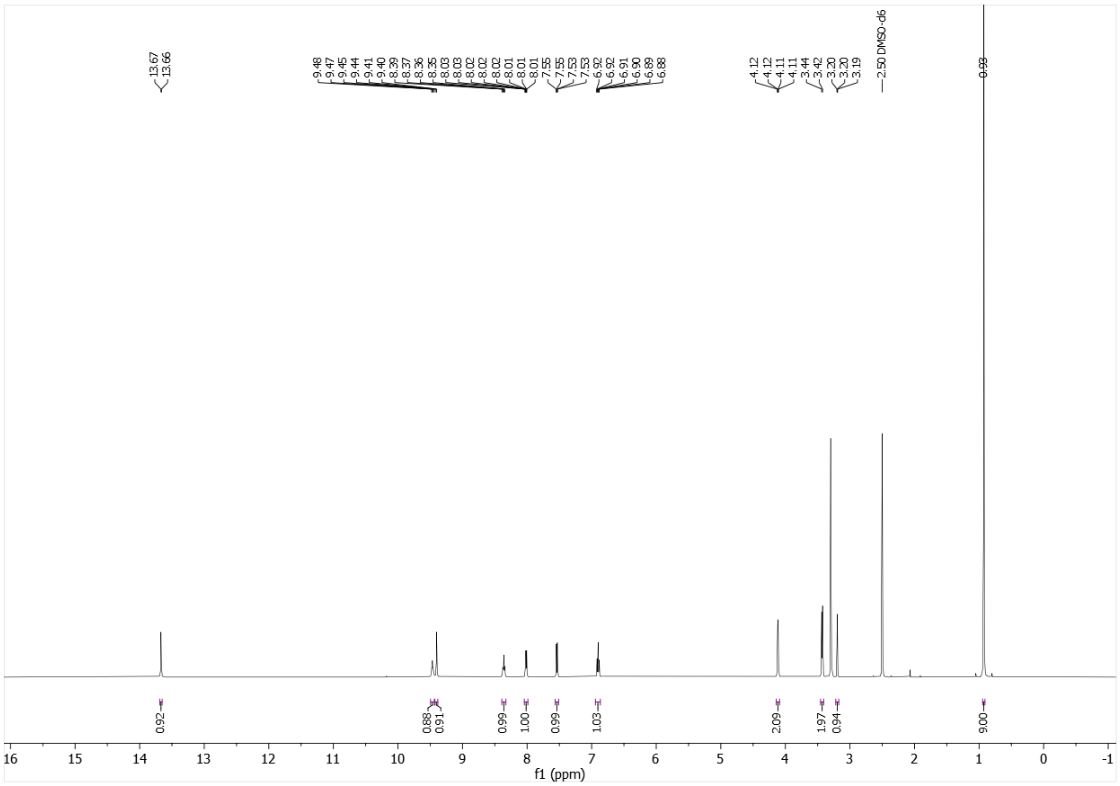

$^{1}$H-NMR (500 MHz, DMSO-$d_6$) of **21a**.

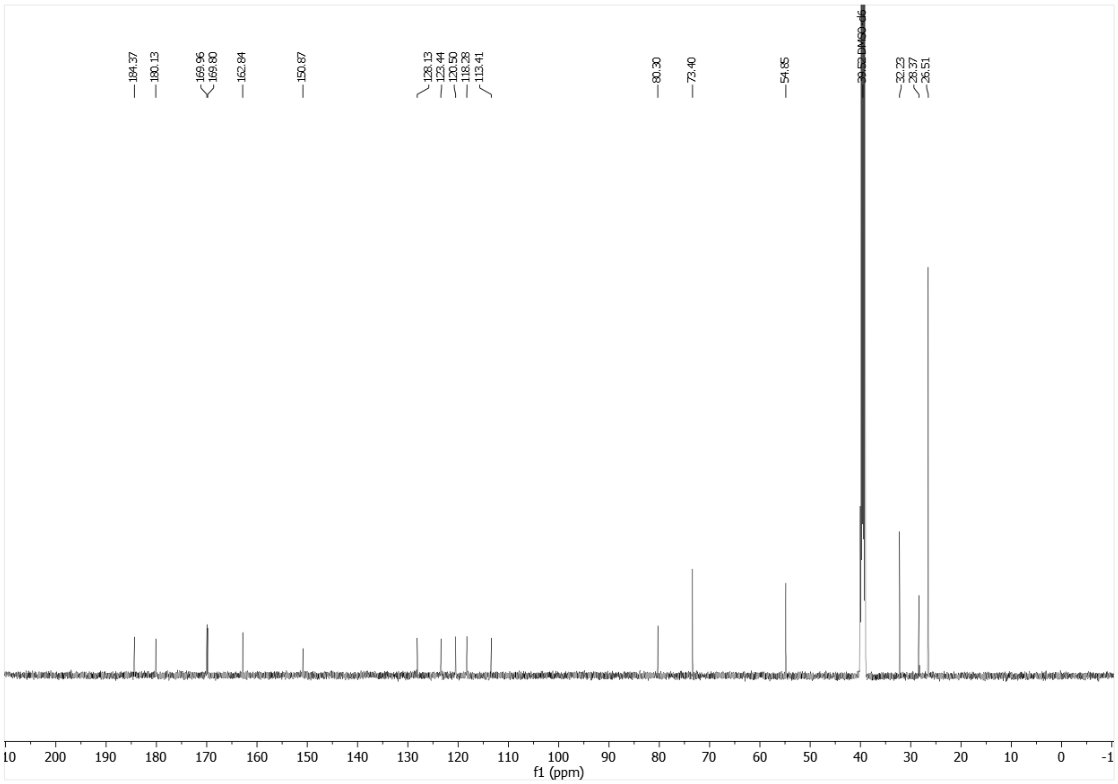

$^{13}$C-NMR (126 MHz, DMSO-$d_6$) of **21a**.

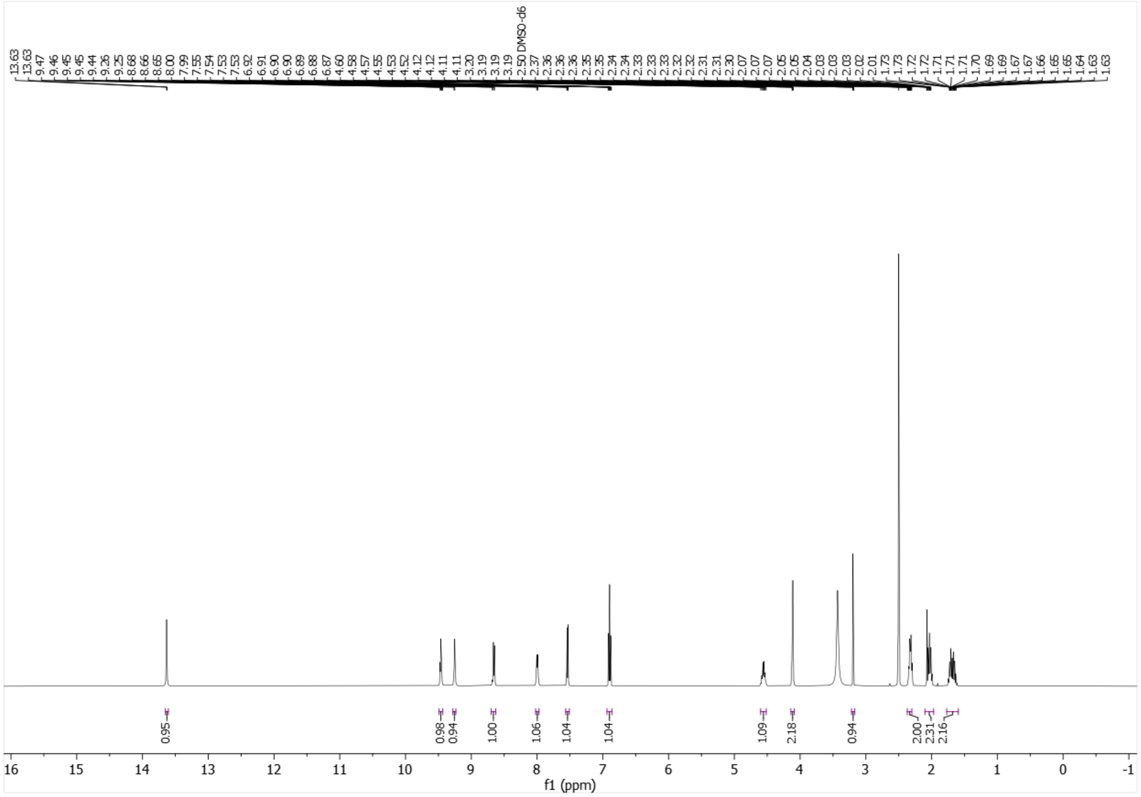

$^{1}$H-NMR (500 MHz, DMSO-$d_6$) of **21b**.

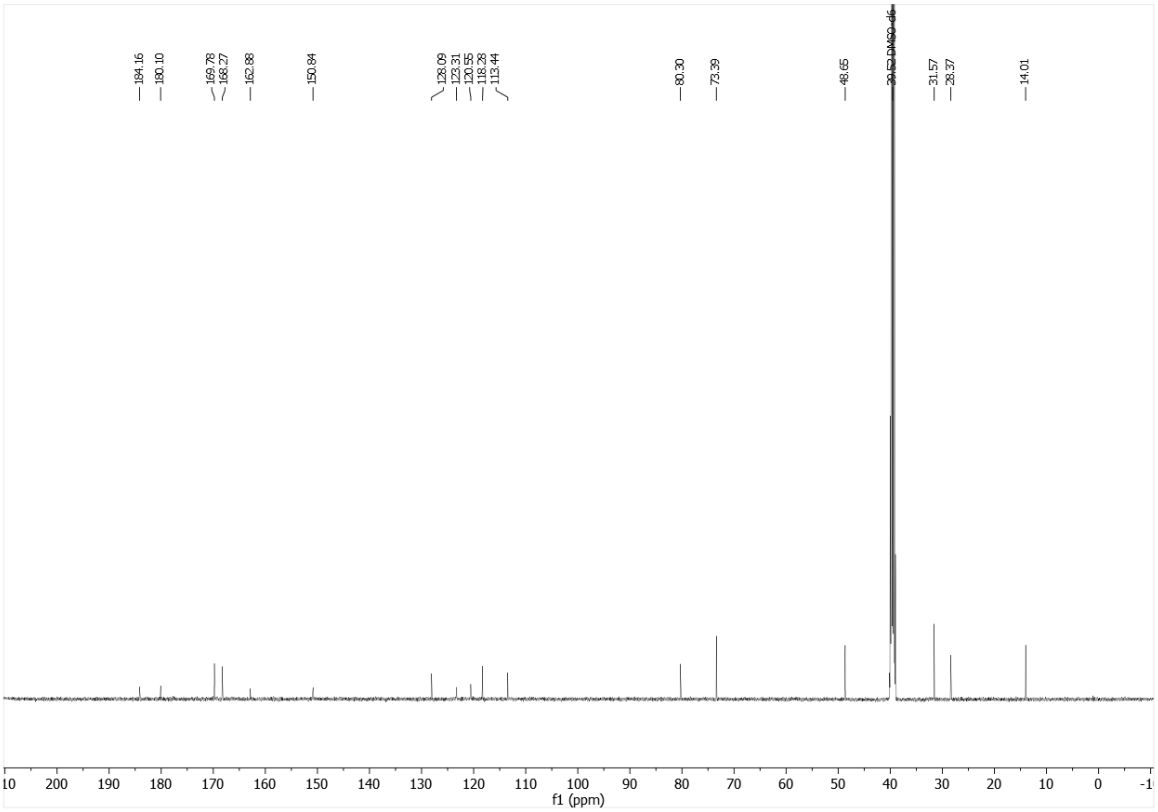

$^{13}$C-NMR (126 MHz, DMSO-$d_6$) of **21b**.

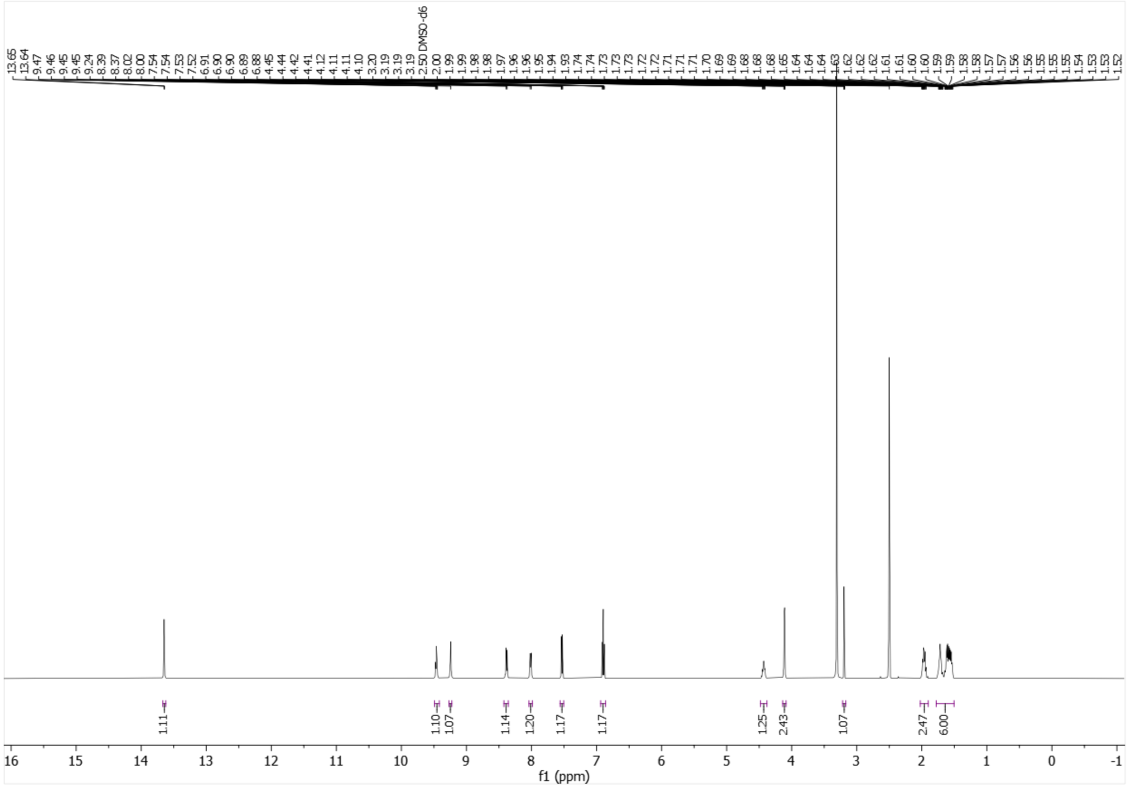


[1]H-NMR (500 MHz, DMSO-$d_6$) of **21c**.

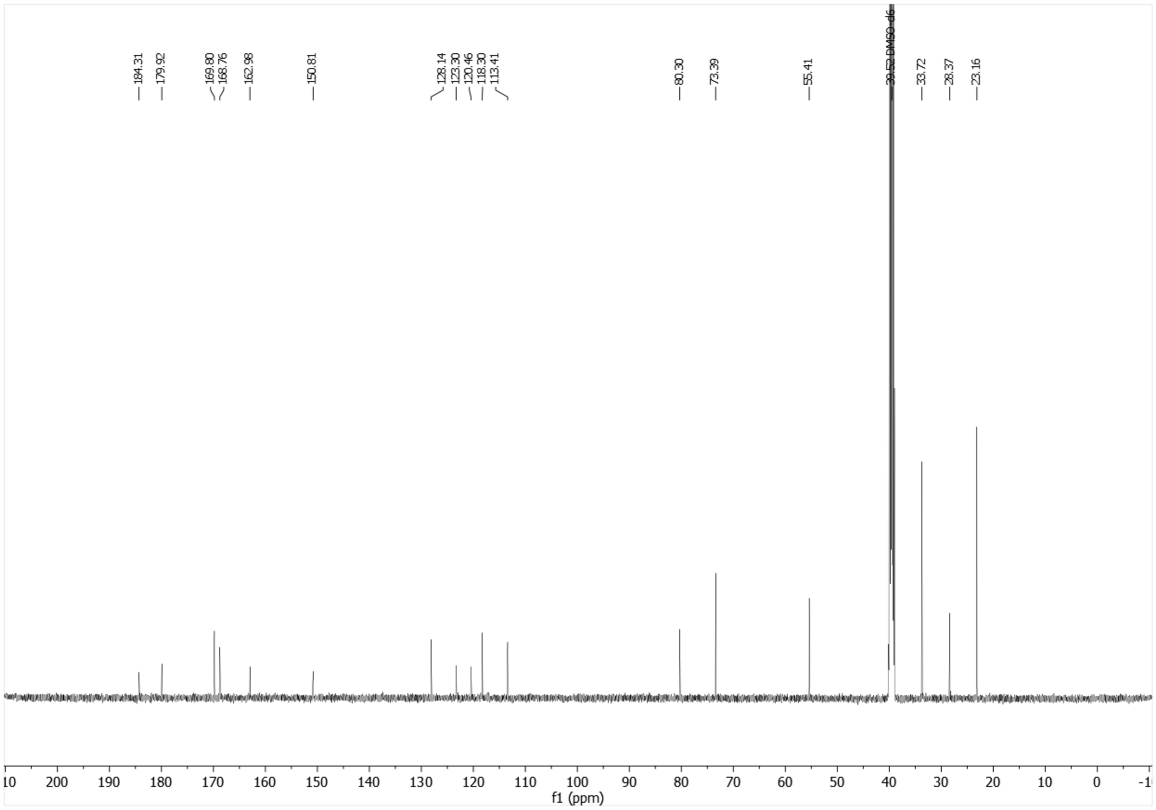


[13]C-NMR (126 MHz, DMSO-$d_6$) of **21c**.

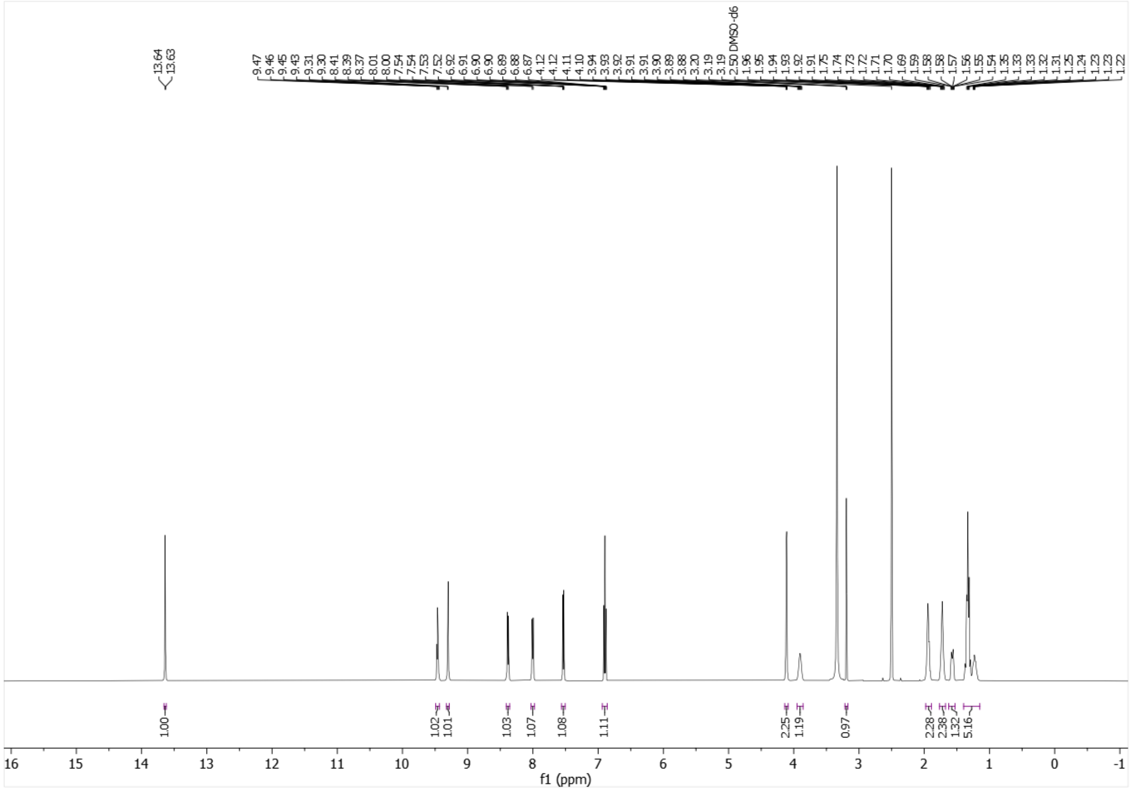


$^{1}$H-NMR (500 MHz, DMSO-$d_6$) of **21d**.

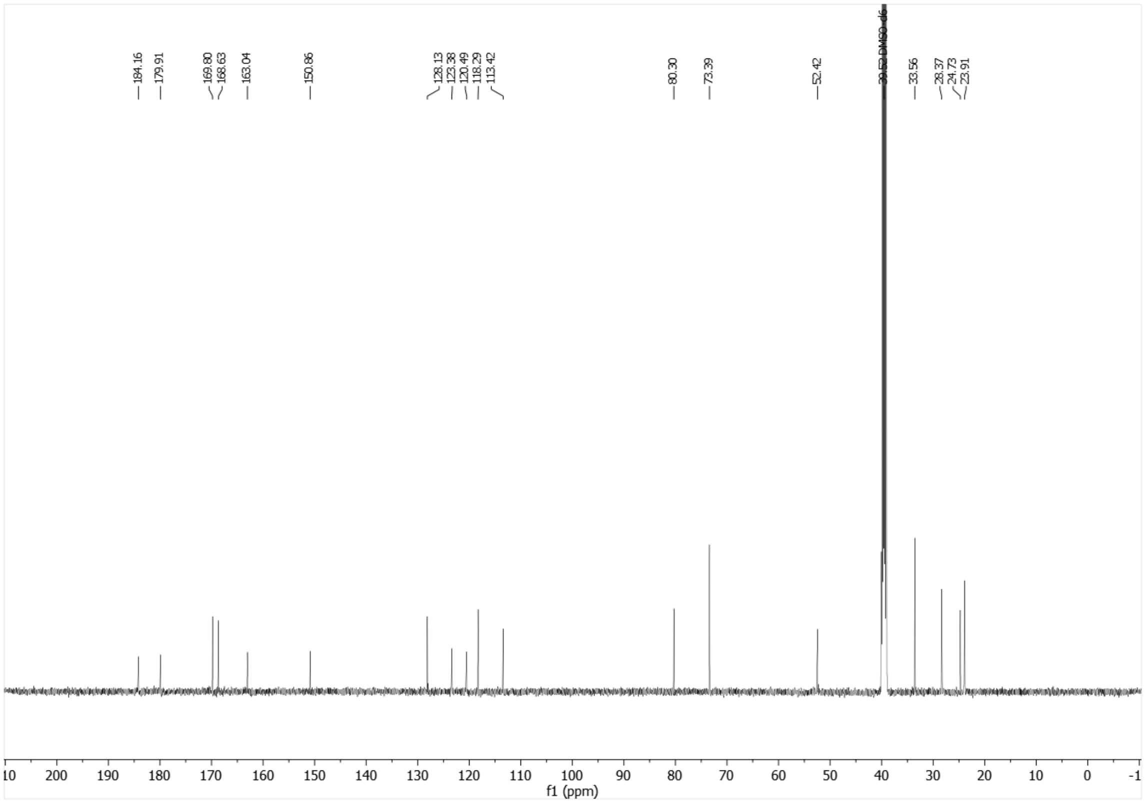


$^{13}$C-NMR (126 MHz, DMSO-$d_6$) of **21d**.

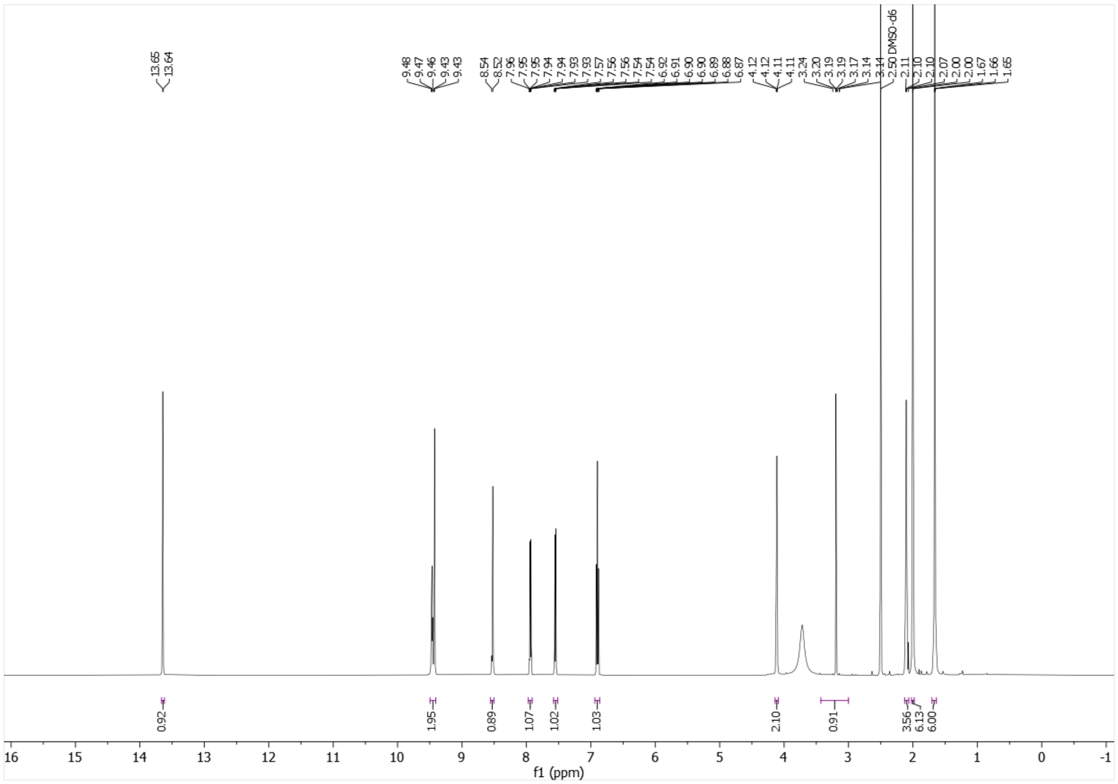


[1]H-NMR (500 MHz, DMSO-$d_6$) of **21e**.

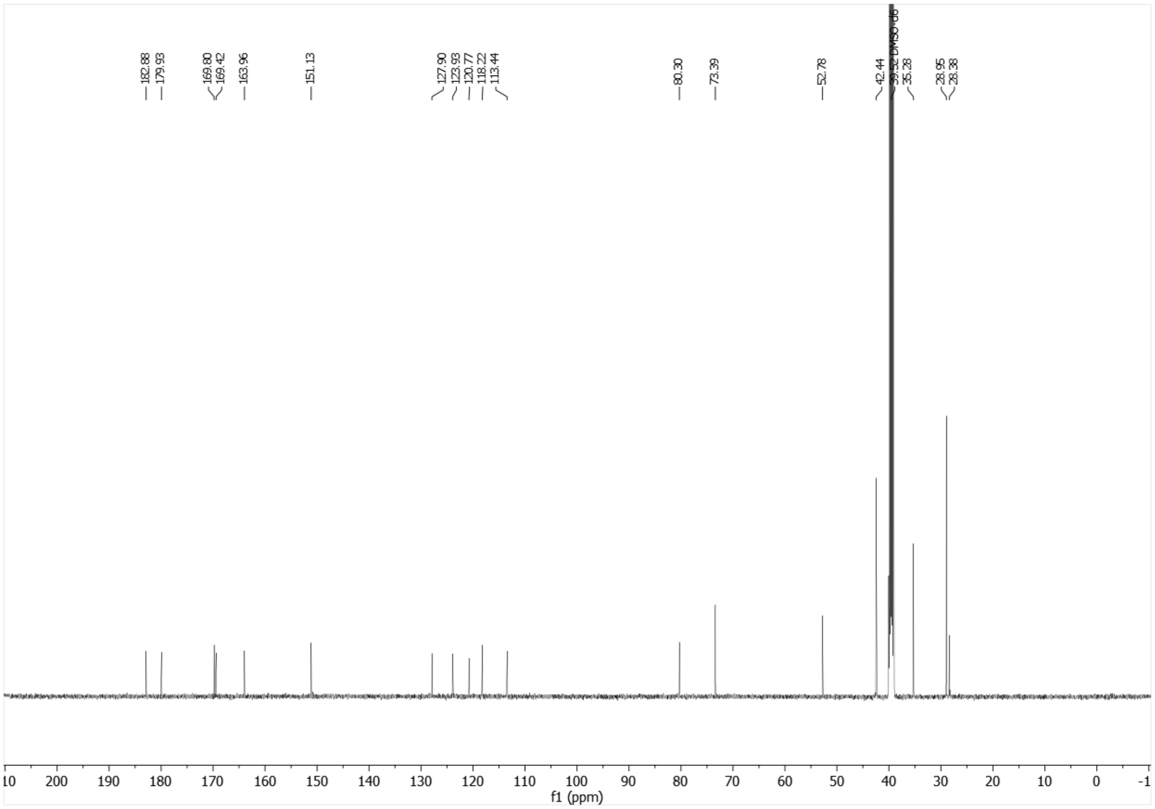


[13]C-NMR (126 MHz, DMSO-$d_6$) of **21e**.

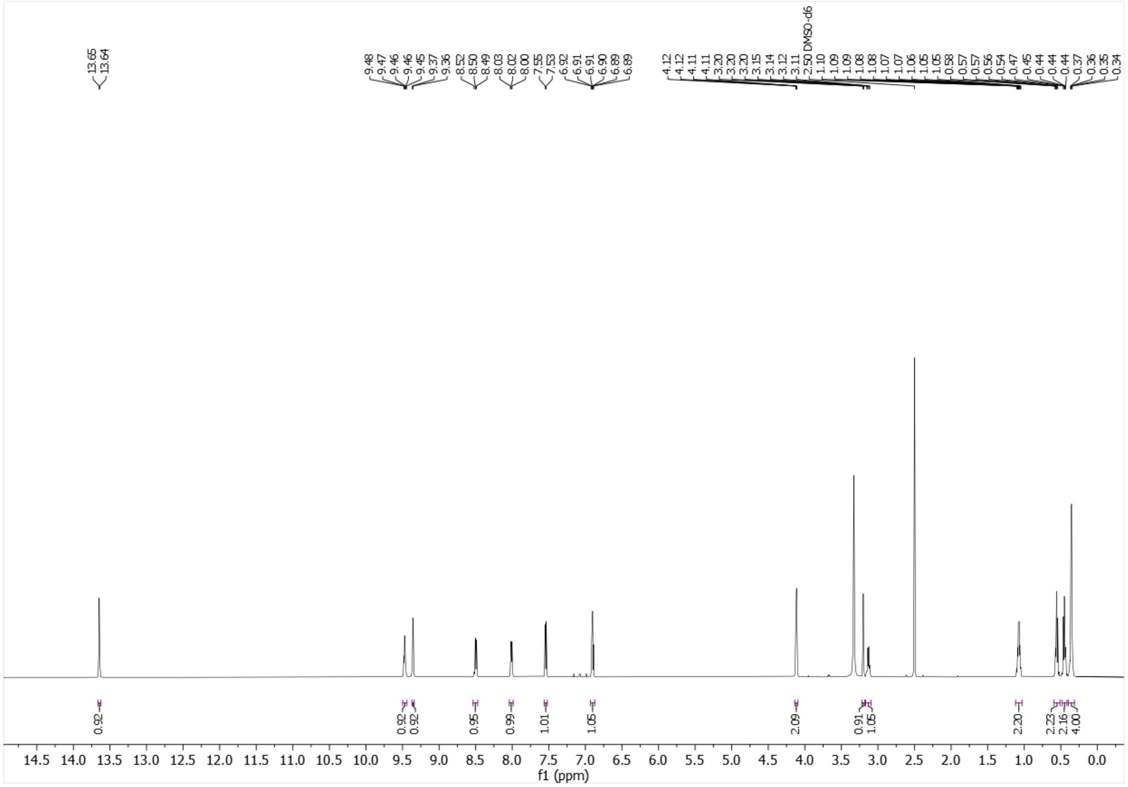

[1]H-NMR (500 MHz, DMSO-$d_6$) of **21f**.

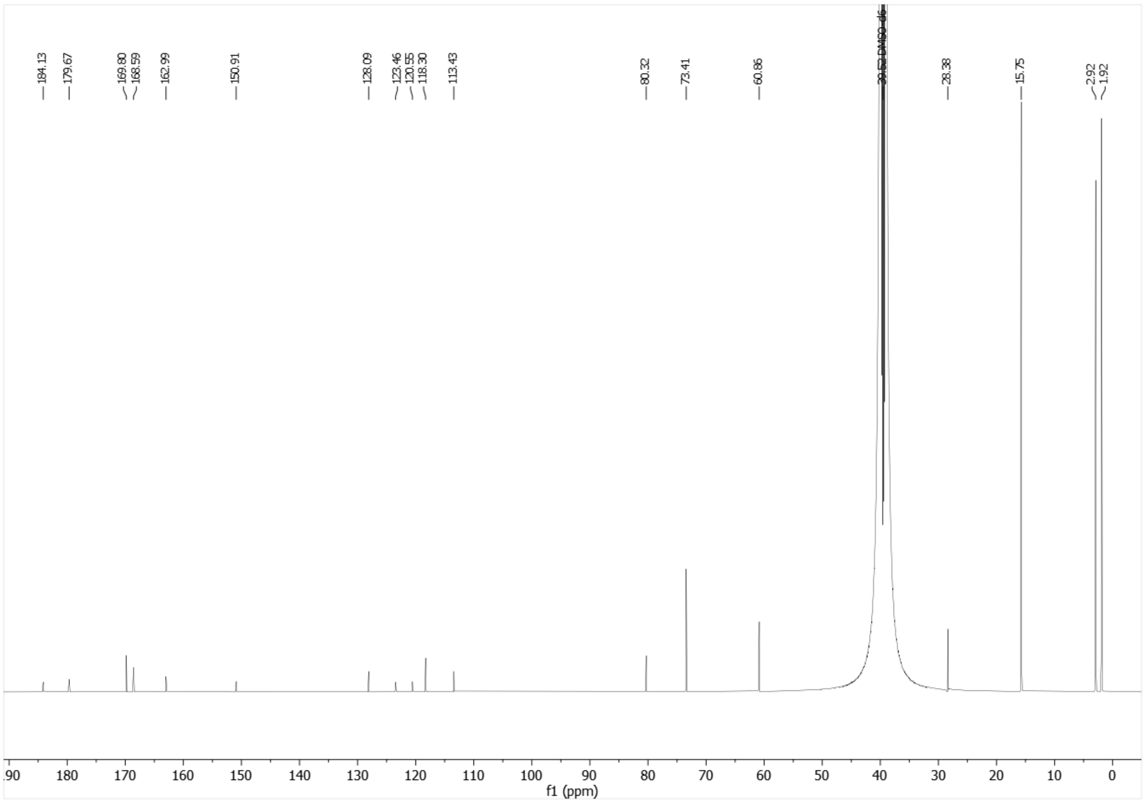

[13]C-NMR (126 MHz, DMSO-$d_6$) of **21f**.

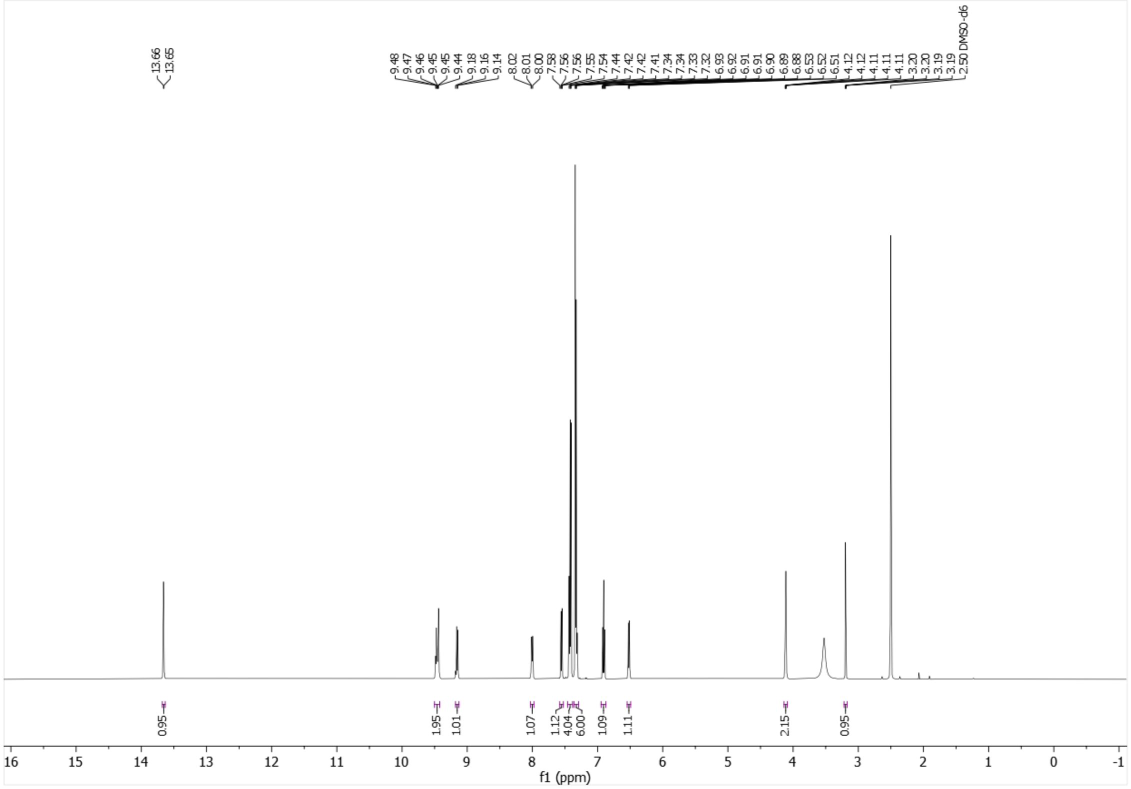

[1]H-NMR (500 MHz, DMSO-$d_6$) of **21g**.

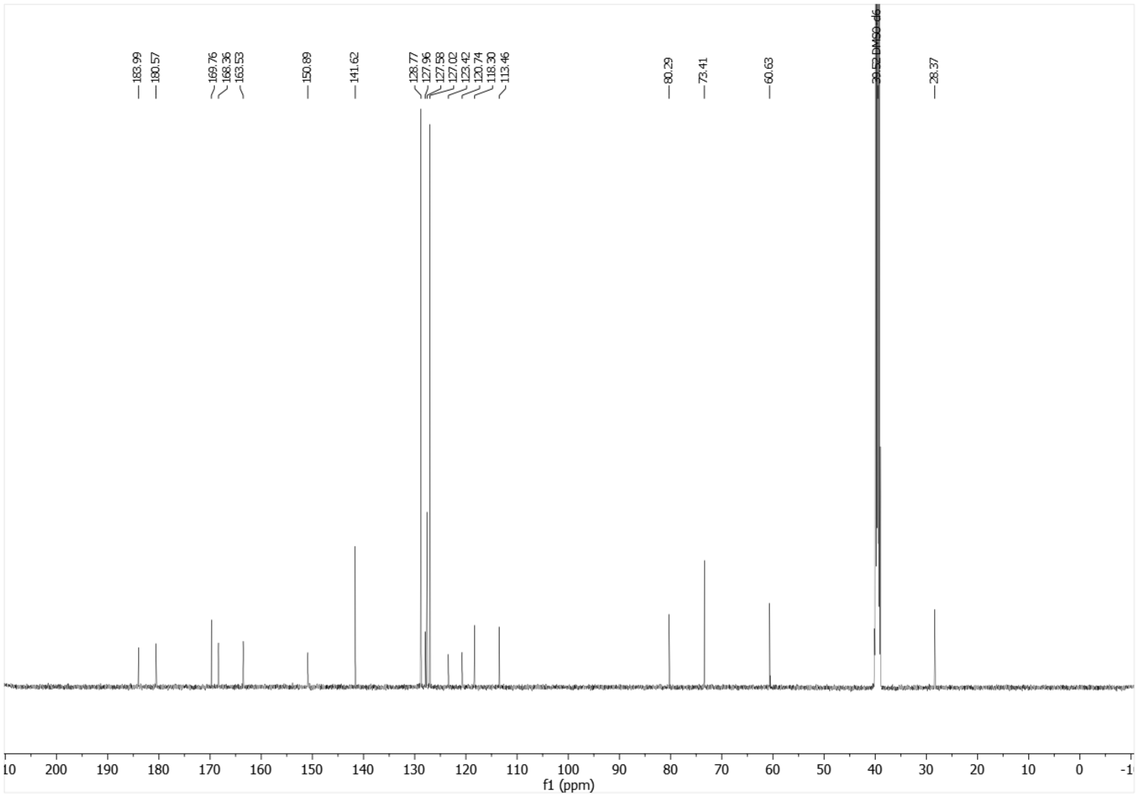

[13]C-NMR (126 MHz, DMSO-$d_6$) of **21g**.

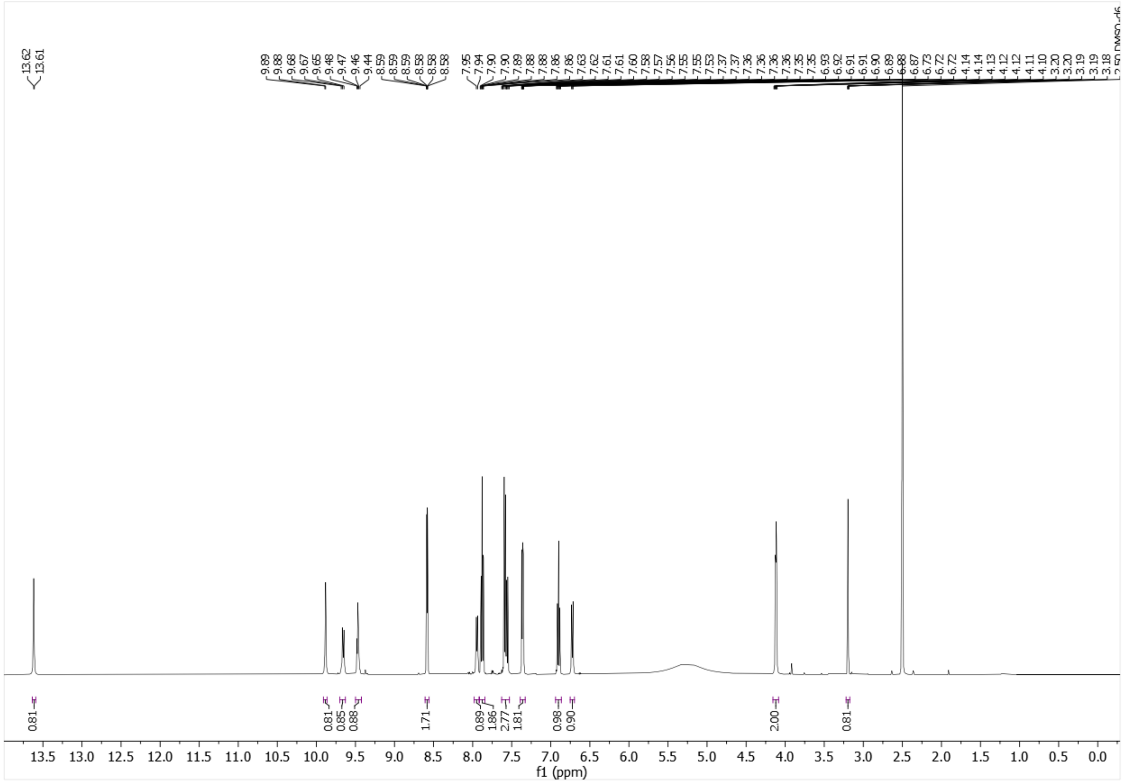

[1]H-NMR (500 MHz, DMSO-$d_6$) of **21h**.

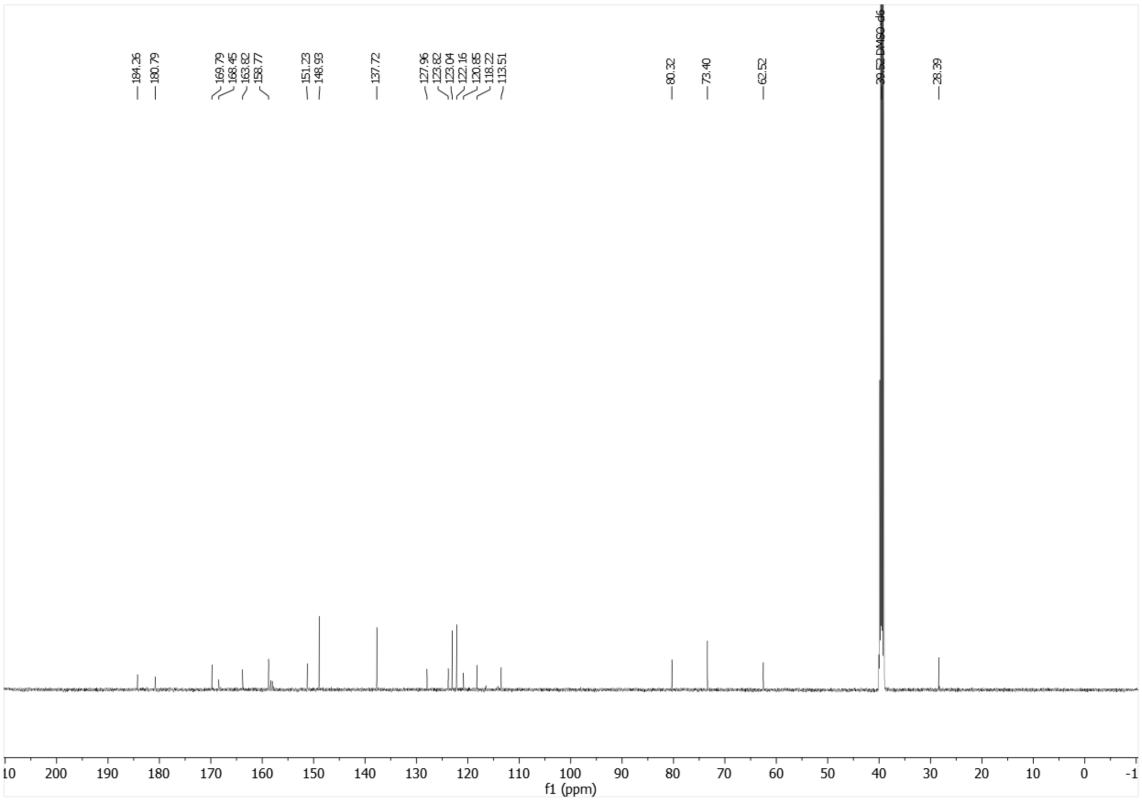

[13]C-NMR (126 MHz, DMSO-$d_6$) of **21h**.

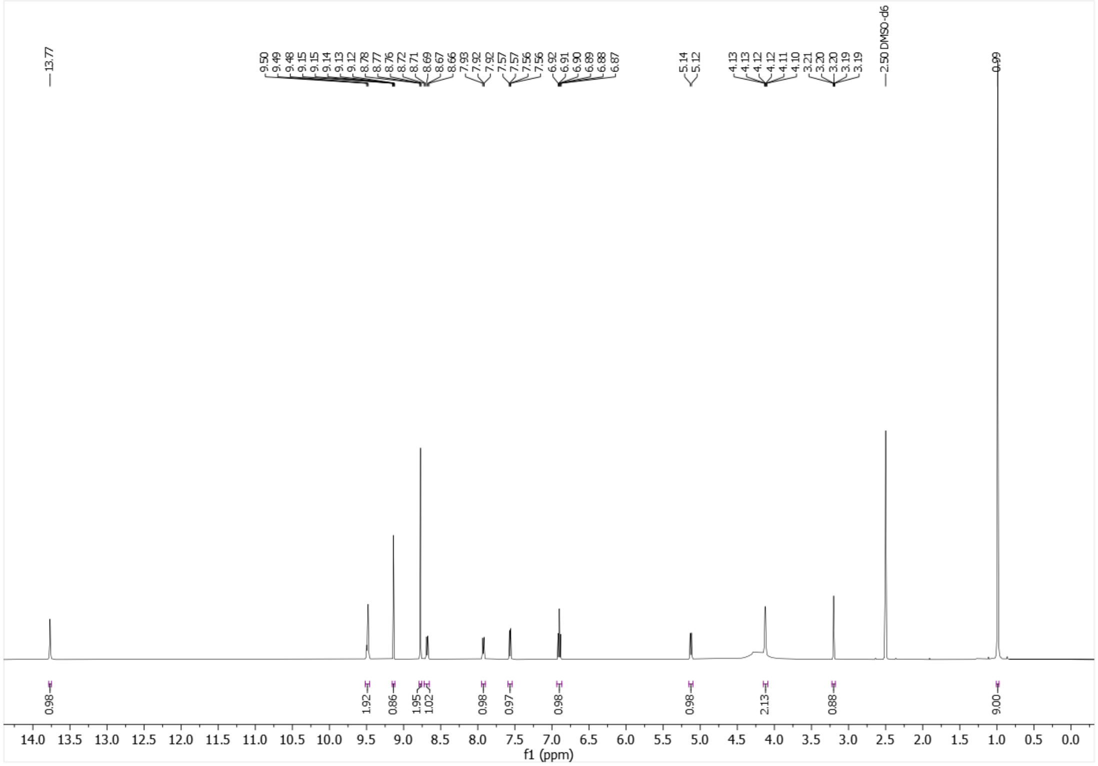

[1]H-NMR (500 MHz, DMSO-$d_6$) of **21i**.

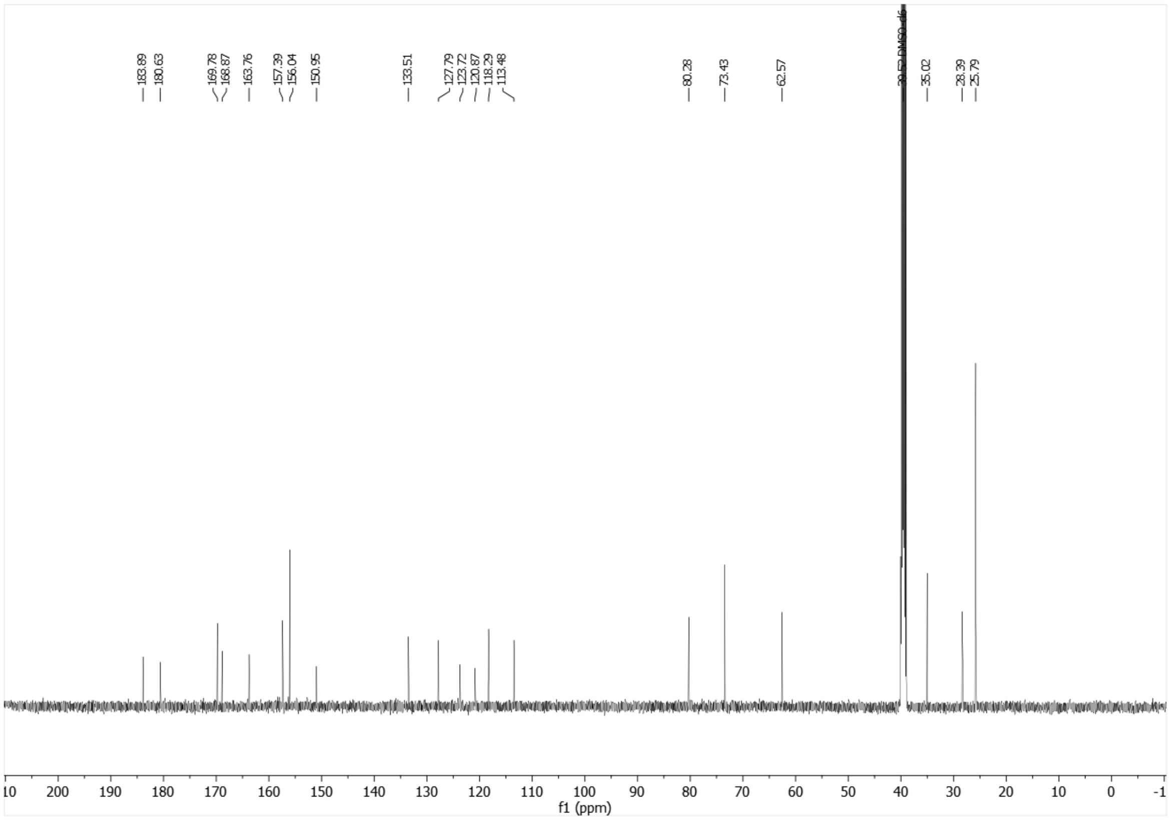

[13]C-NMR (126 MHz, DMSO-$d_6$) of **21i**.

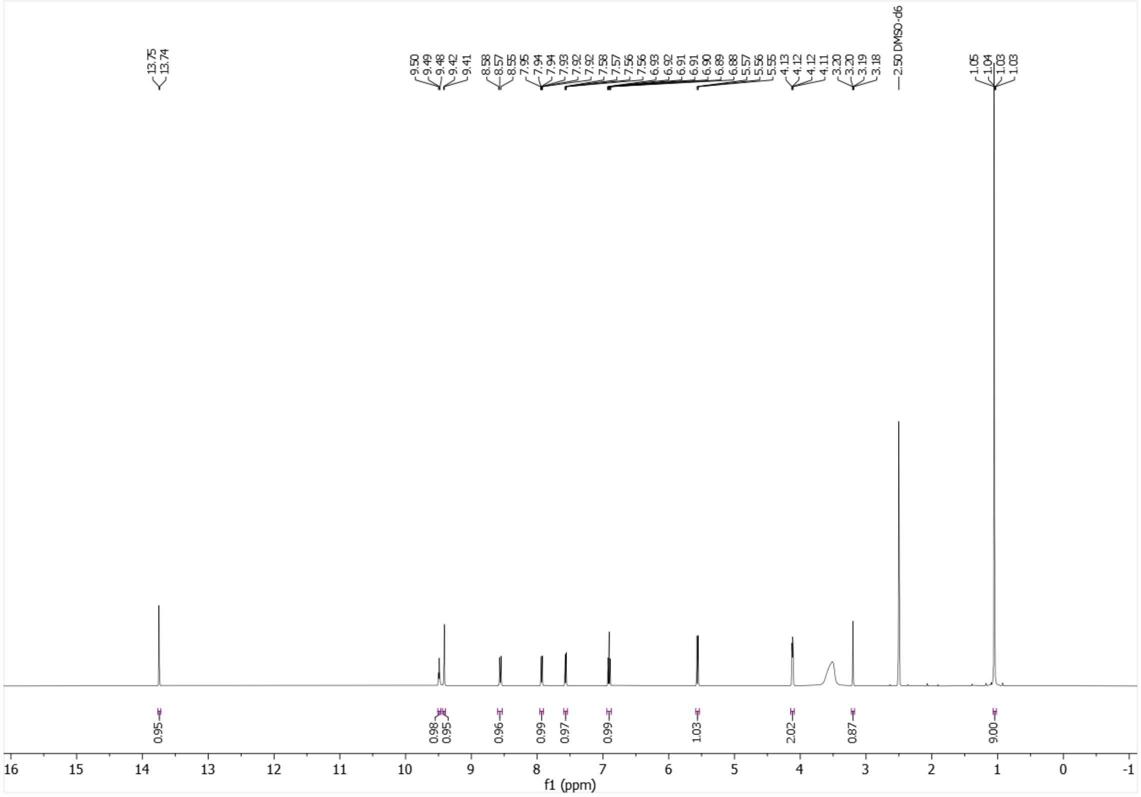


[1]H-NMR (500 MHz, DMSO-$d_6$) of **21j**.

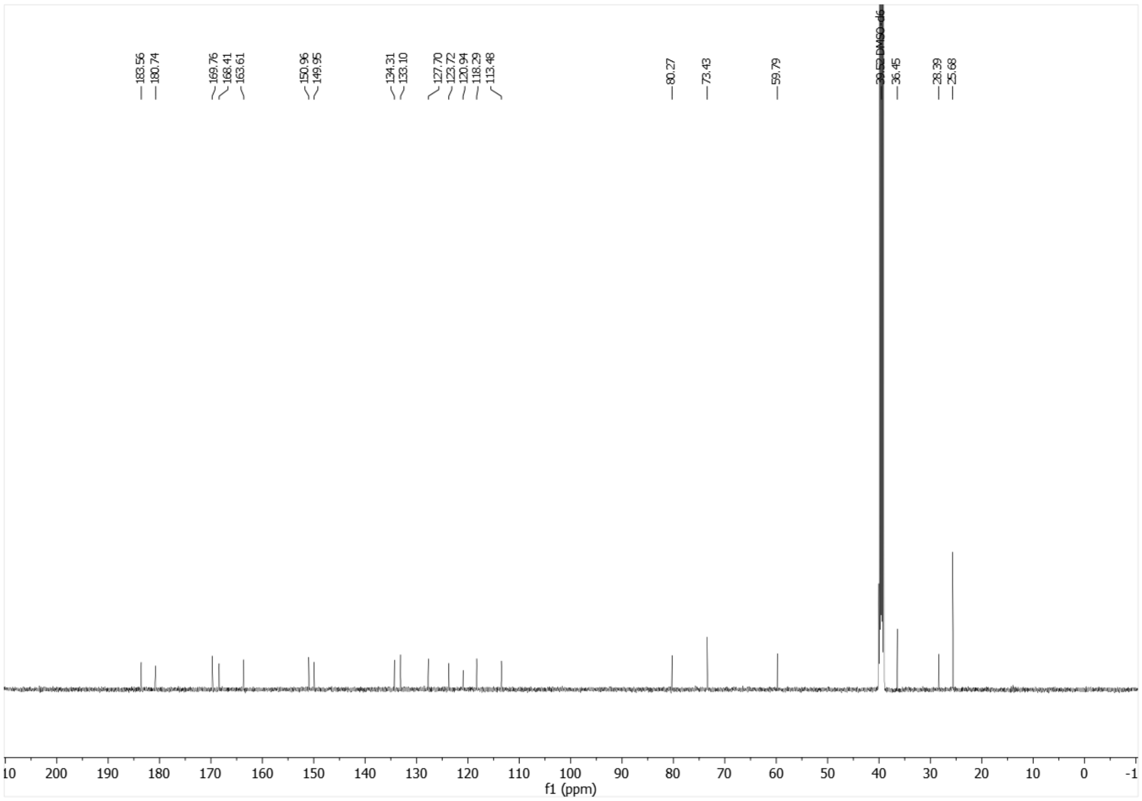


[13]C-NMR (126 MHz, DMSO-$d_6$) of **21j**.

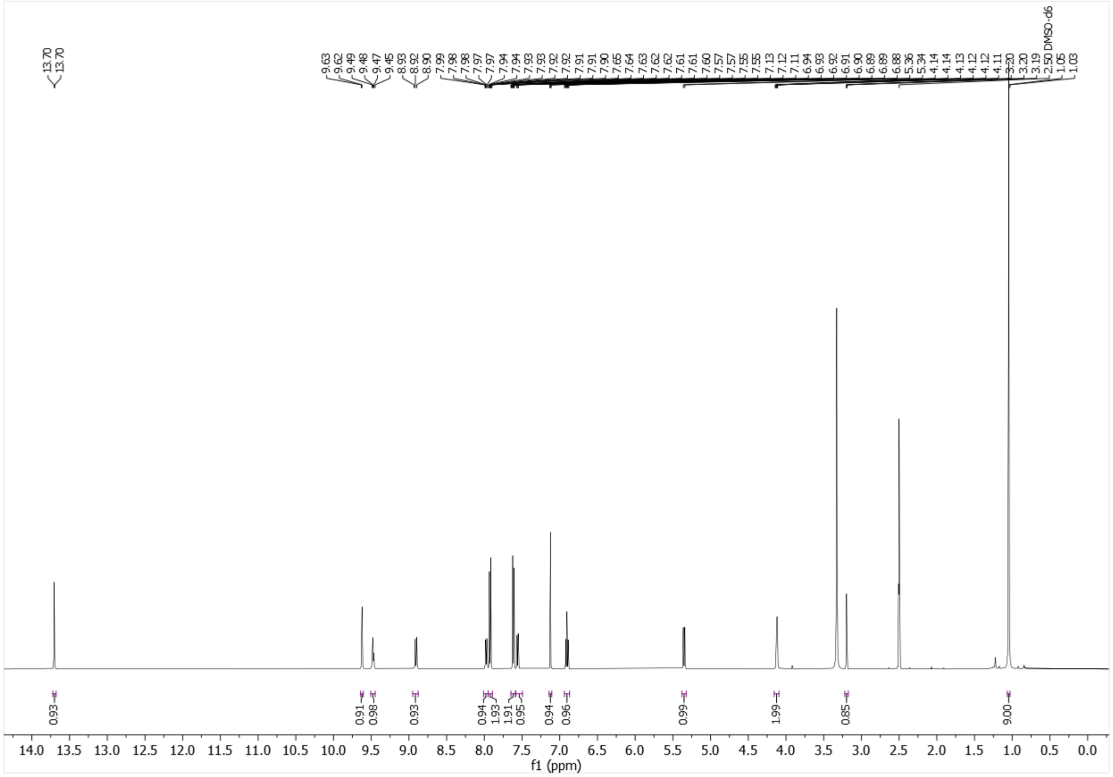


$^{1}$H-NMR (500 MHz, DMSO-$d_6$) of **21k**.

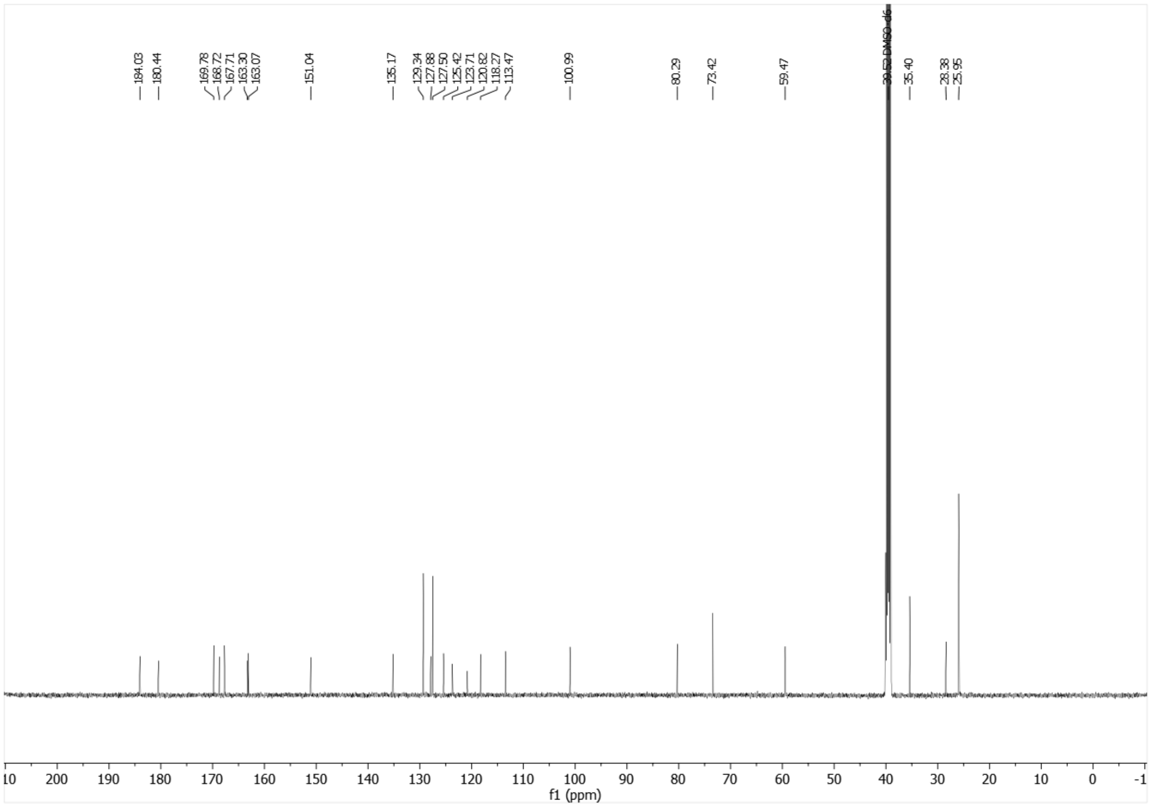


$^{13}$C-NMR (126 MHz, DMSO-$d_6$) of **21k**.

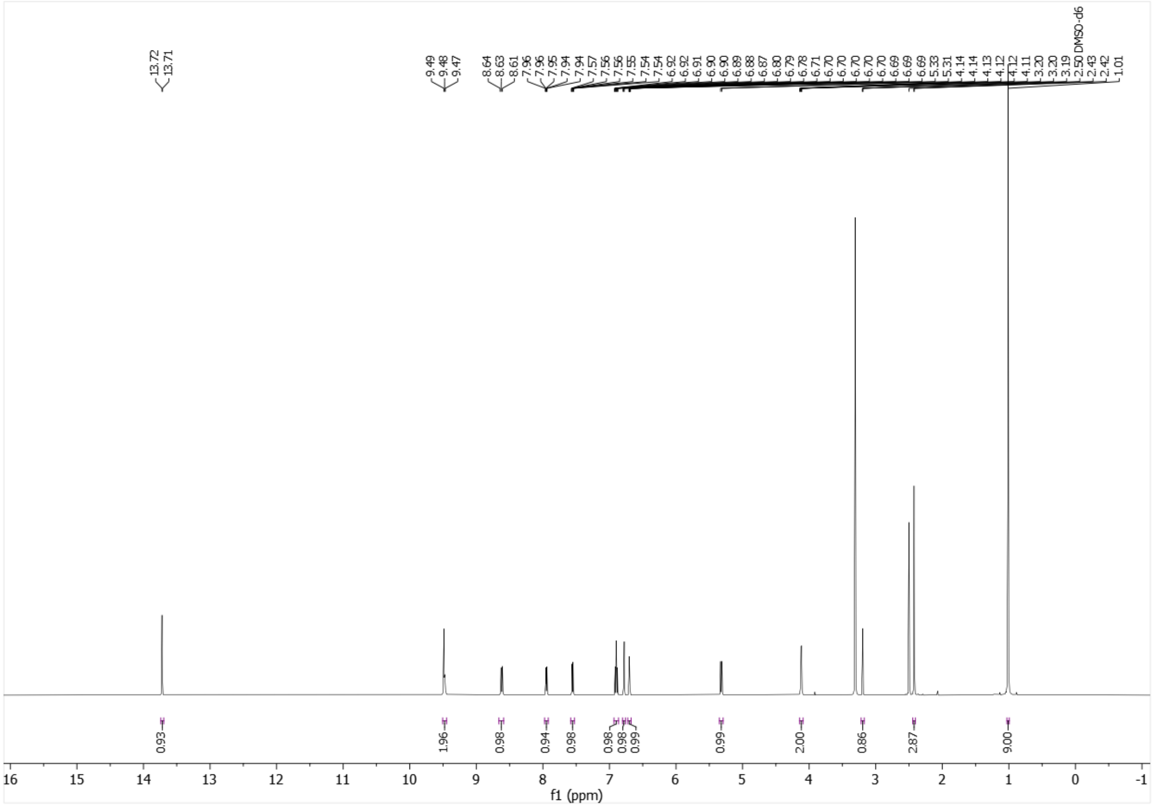


[1]H-NMR (500 MHz, DMSO-$d_6$) of **21l**.

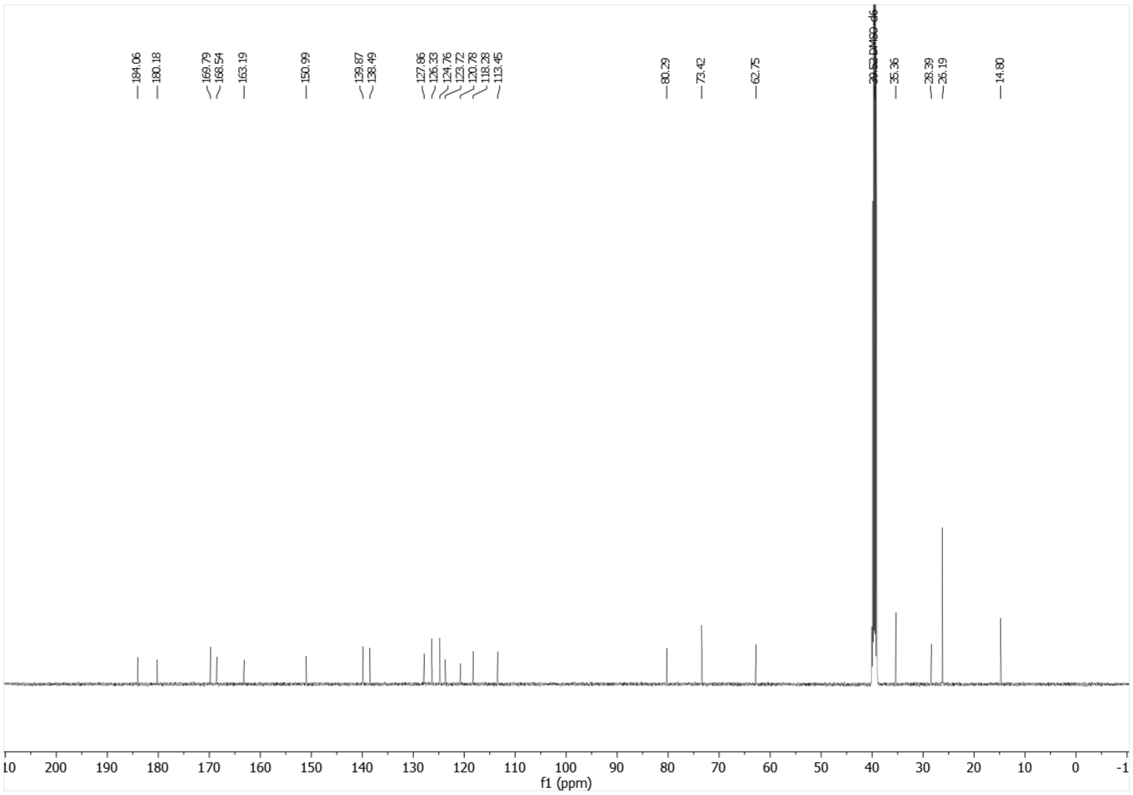


[13]C-NMR (126 MHz, DMSO-$d_6$) of **21l**.

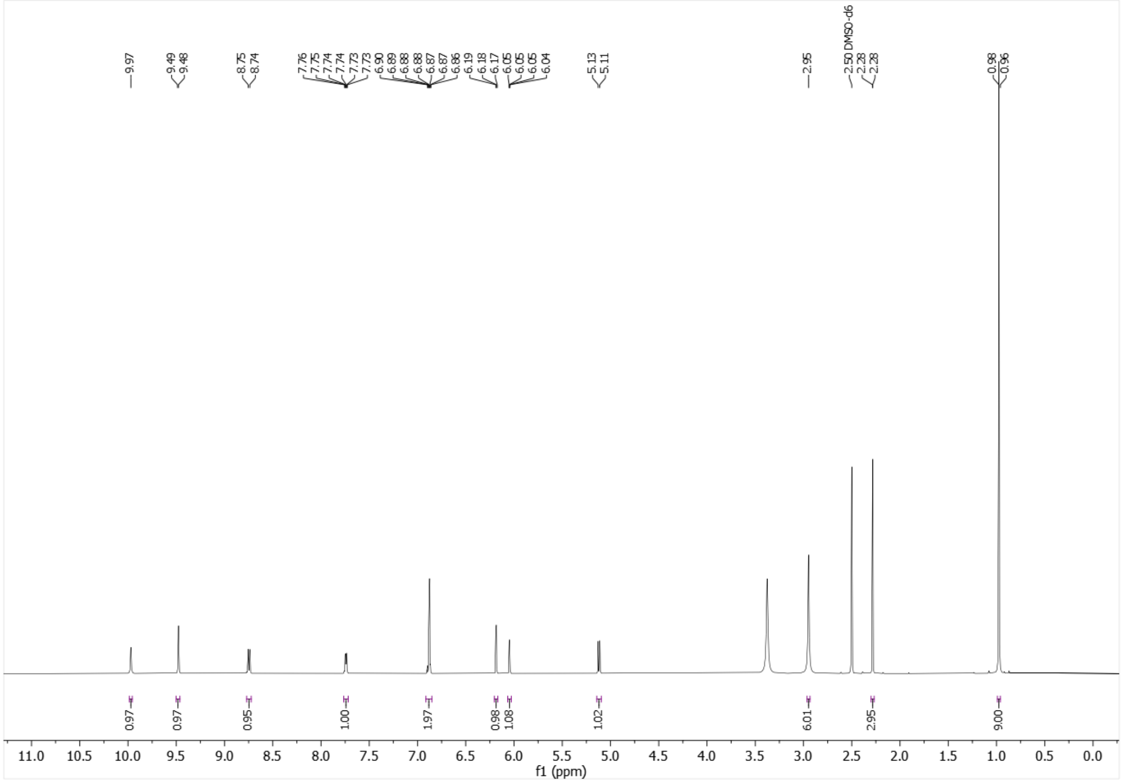


[1]H-NMR (400 MHz, DMSO-$d_6$) of SLW132 (**21m**).

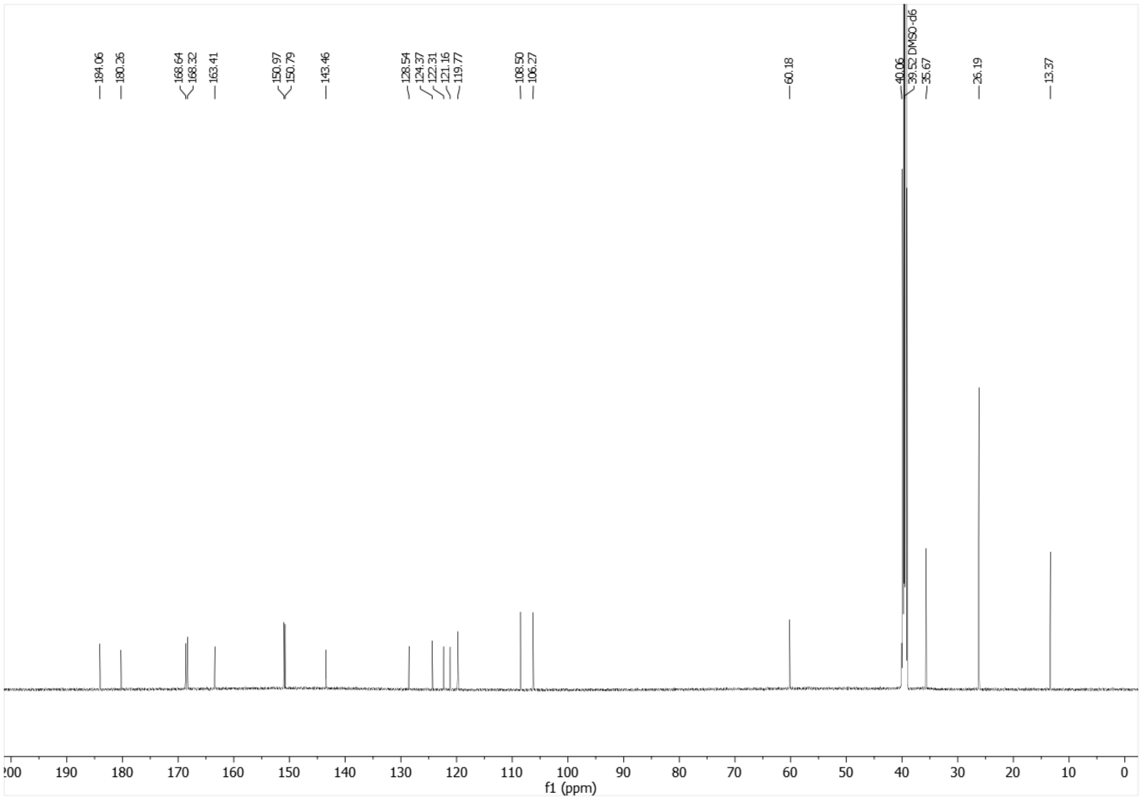


[13]C-NMR (151 MHz, DMSO-$d_6$) of SLW132 (**21m**).

### HPLC CHROMATOGRAMS OF TARGET COMPOUNDS

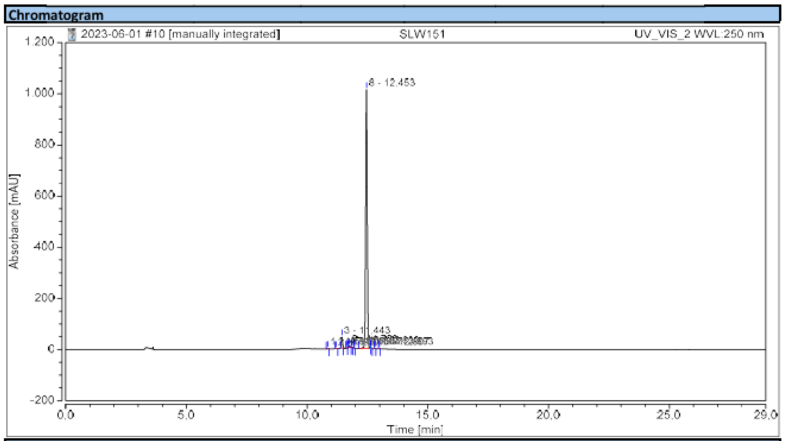


**Integration Results**

| No. | Peak Name | Retention Time min | Area mAU*min | Height mAU | Relative Area % | Relative Height % | Amount n.a. |
|---|---|---|---|---|---|---|---|
| 1 | | 10,843 | 0,143 | 2,275 | 0,19 | 0,21 | n.a. |
| 2 | | 11,207 | 0,047 | 0,733 | 0,06 | 0,07 | n.a. |
| 3 | | 11,443 | 2,436 | 43,598 | 3,20 | 4,03 | n.a. |
| 4 | | 11,653 | 0,071 | 1,572 | 0,09 | 0,15 | n.a. |
| 5 | | 11,750 | 0,483 | 8,162 | 0,63 | 0,75 | n.a. |
| 6 | | 11,853 | 0,091 | 2,132 | 0,12 | 0,20 | n.a. |
| 7 | | 11,957 | 0,025 | 0,810 | 0,03 | 0,07 | n.a. |
| 8 | | 12,453 | 72,358 | 1013,740 | 95,08 | 93,62 | n.a. |
| 9 | | 12,630 | 0,296 | 6,521 | 0,39 | 0,60 | n.a. |
| 10 | | 12,807 | 0,065 | 1,487 | 0,08 | 0,14 | n.a. |
| 11 | | 12,993 | 0,083 | 1,754 | 0,11 | 0,16 | n.a. |
| **Total:** | | | **76,099** | **1082,785** | **100,00** | **100,00** | |

HPLC chromatogram of **3**.

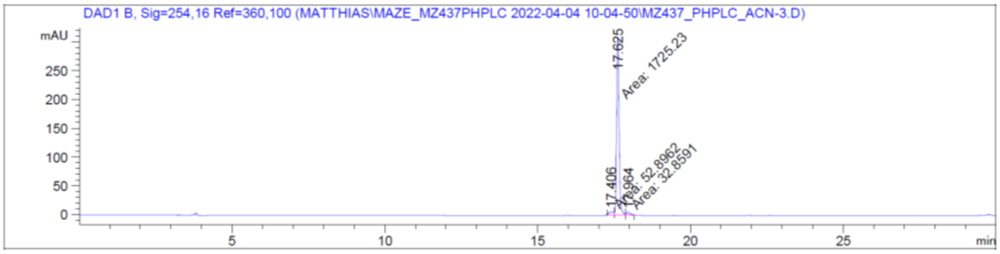


```
==================================================================
                       Area Percent Report
==================================================================

Sorted By              :      Signal
Multiplier             :      1.0000
Dilution               :      1.0000
Use Multiplier & Dilution Factor with ISTDs

        Signal 2: DAD1 B, Sig=254,16 Ref=360,100

        Peak RetTime Type  Width     Area      Height     Area
          #   [min]        [min]   [mAU*s]     [mAU]       %
        ----|-------|----|-------|----------|----------|--------|
           1  17.406 MF    0.1777   52.89622    4.96240   2.9209
           2  17.625 MF    0.0929 1725.22974  309.50720  95.2647
           3  17.964 FM    0.1302   32.85914    4.20531   1.8144

        Totals :                  1810.98510  318.67491
```

HPLC chromatogram of Mz437 (**4**).

**Chromatogram**

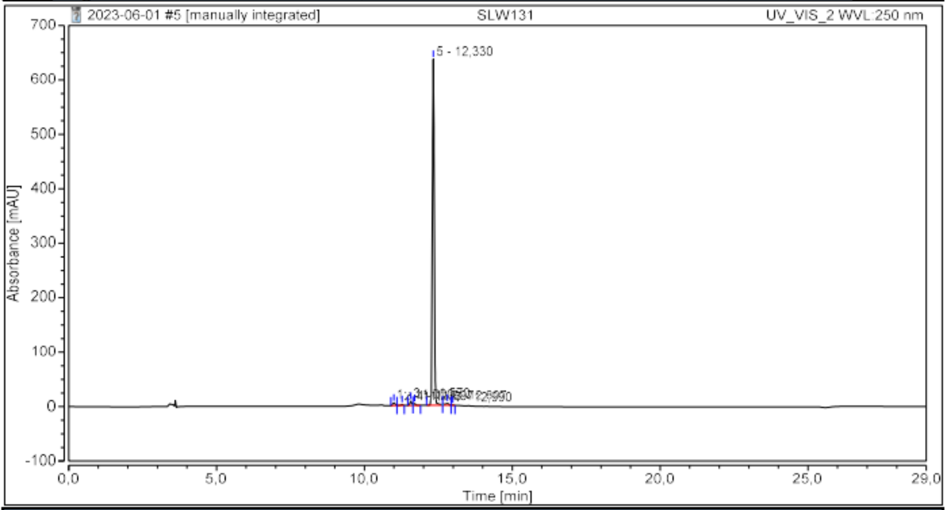


**Integration Results**

| No. | Peak Name | Retention Time min | Area mAU*min | Height mAU | Relative Area % | Relative Height % | Amount n.a. |
|---|---|---|---|---|---|---|---|
| 1 | | 11,000 | 0,479 | 5,130 | 1,01 | 0,78 | n.a. |
| 2 | | 11,273 | 0,183 | 2,128 | 0,39 | 0,32 | n.a. |
| 3 | | 11,570 | 0,561 | 7,728 | 1,19 | 1,17 | n.a. |
| 4 | | 11,697 | 0,274 | 3,466 | 0,58 | 0,53 | n.a. |
| 5 | | 12,330 | 45,131 | 635,638 | 95,55 | 96,46 | n.a. |
| 6 | | 12,807 | 0,552 | 4,103 | 1,17 | 0,62 | n.a. |
| 7 | | 12,990 | 0,054 | 0,785 | 0,11 | 0,12 | n.a. |
| **Total:** | | | **47,234** | **658,978** | **100,00** | **100,00** | |

HPLC chromatogram of SLW131 (**10**).

**Chromatogram**

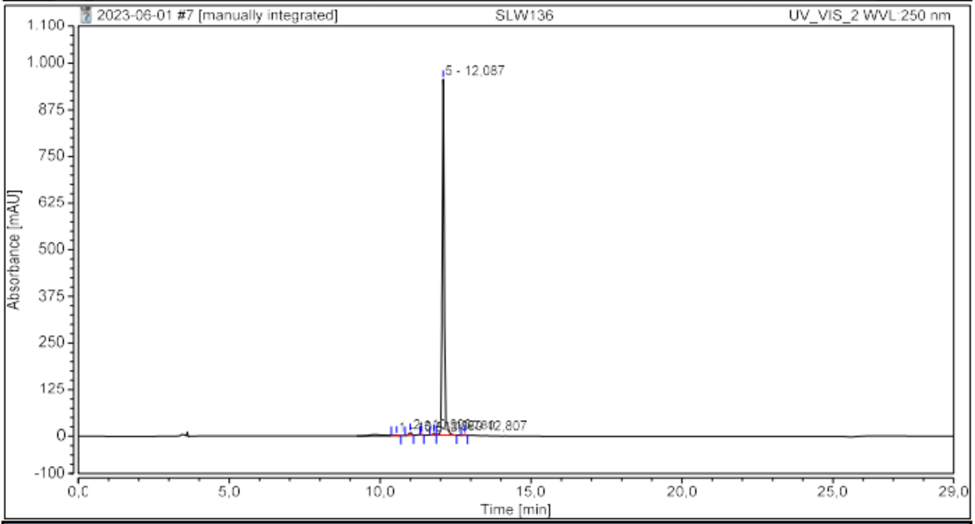


**Integration Results**

| No. | Peak Name | Retention Time min | Area mAU*min | Height mAU | Relative Area % | Relative Height % | Amount n.a. |
|---|---|---|---|---|---|---|---|
| 1 | | 10,543 | 0,250 | 1,240 | 0,33 | 0,13 | n.a. |
| 2 | | 10,993 | 0,720 | 7,566 | 0,95 | 0,78 | n.a. |
| 3 | | 11,370 | 0,065 | 1,023 | 0,09 | 0,11 | n.a. |
| 4 | | 11,780 | 0,365 | 4,639 | 0,48 | 0,48 | n.a. |
| 5 | | 12,087 | 74,497 | 954,262 | 97,90 | 98,26 | n.a. |
| 6 | | 12,807 | 0,199 | 2,448 | 0,26 | 0,25 | n.a. |
| **Total:** | | | **76,096** | **971,177** | **100,00** | **100,00** | |

HPLC chromatogram of **20a**.

**Chromatogram**

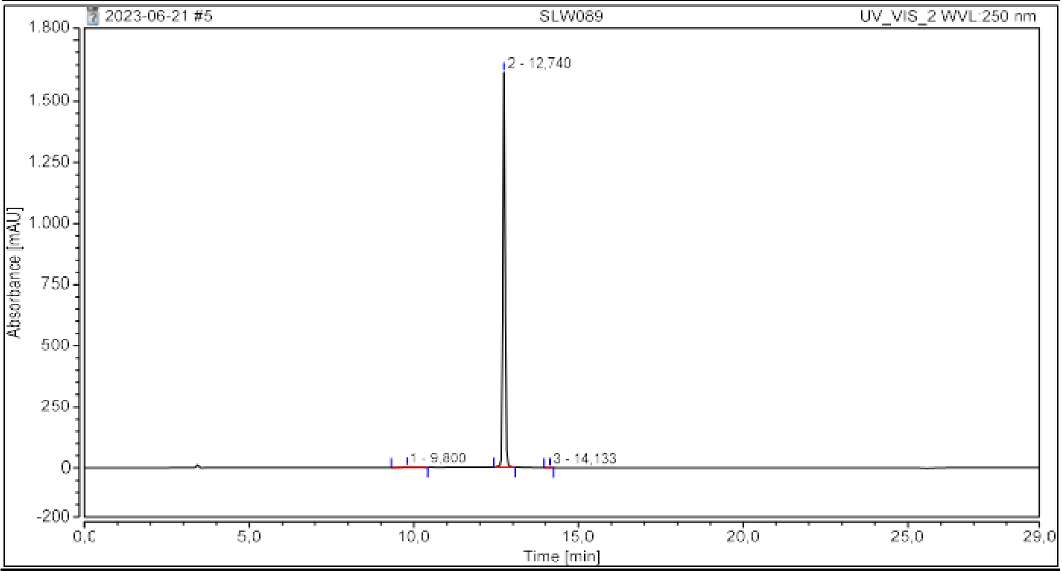


**Integration Results**

| No. | Peak Name | Retention Time min | Area mAU*min | Height mAU | Relative Area % | Relative Height % | Amount n.a. |
|---|---|---|---|---|---|---|---|
| 1 | | 9,800 | 1,139 | 2,700 | 0,86 | 0,17 | n.a. |
| 2 | | 12,740 | 130,727 | 1615,520 | 99,09 | 99,80 | n.a. |
| 3 | | 14,133 | 0,062 | 0,604 | 0,05 | 0,04 | n.a. |
| **Total:** | | | **131,928** | **1618,825** | **100,00** | **100,00** | |

HPLC chromatogram of **20b**.

**Chromatogram**

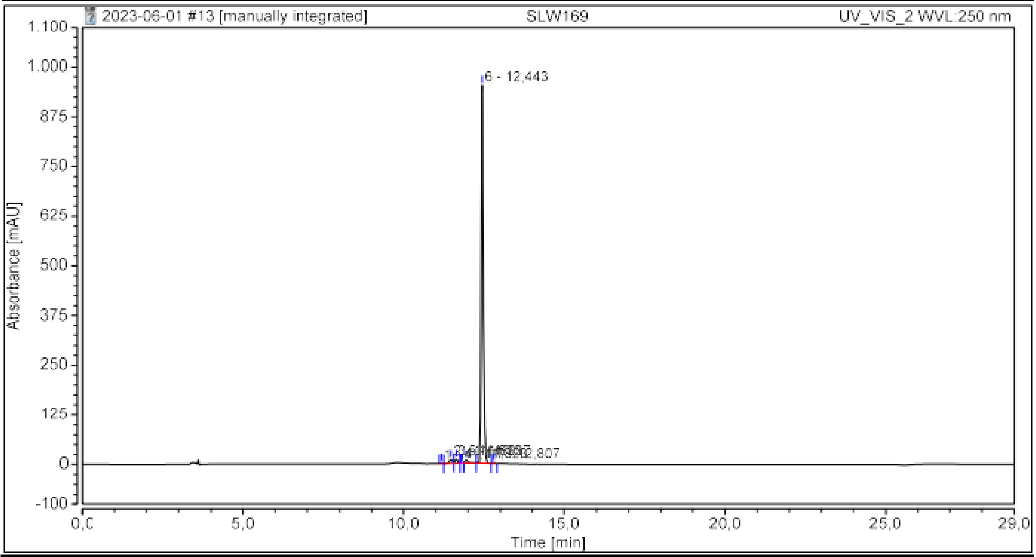


**Integration Results**

| No. | Peak Name | Retention Time min | Area mAU*min | Height mAU | Relative Area % | Relative Height % | Amount n.a. |
|---|---|---|---|---|---|---|---|
| 1 | | 11,177 | 0,103 | 1,574 | 0,14 | 0,16 | n.a. |
| 2 | | 11,470 | 1,088 | 10,113 | 1,50 | 1,03 | n.a. |
| 3 | | 11,633 | 0,974 | 10,974 | 1,34 | 1,11 | n.a. |
| 4 | | 11,823 | 0,050 | 1,003 | 0,07 | 0,10 | n.a. |
| 5 | | 11,937 | 0,986 | 9,180 | 1,36 | 0,93 | n.a. |
| 6 | | 12,443 | 69,278 | 951,017 | 95,36 | 96,41 | n.a. |
| 7 | | 12,807 | 0,171 | 2,558 | 0,24 | 0,26 | n.a. |
| **Total:** | | | **72,651** | **986,418** | **100,00** | **100,00** | |

HPLC chromatogram of **20c**.

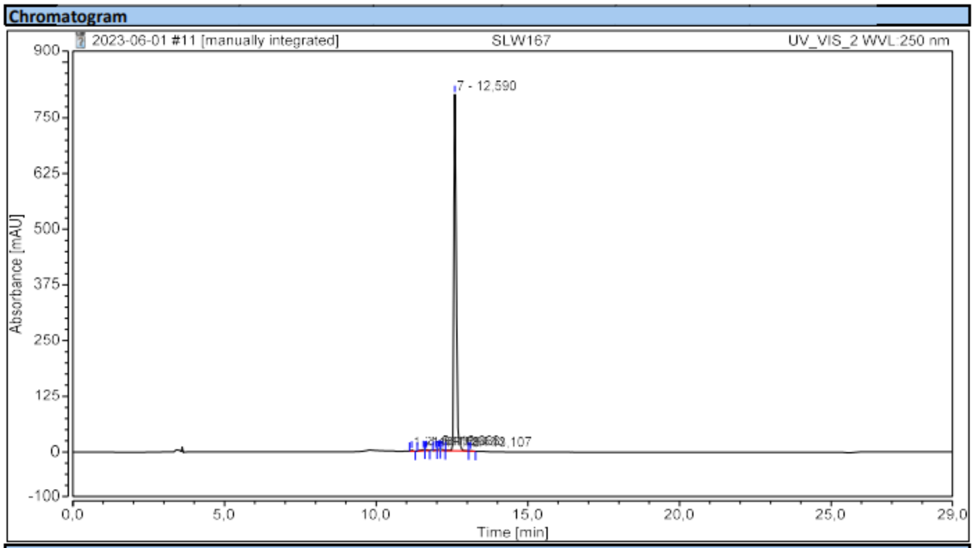


Integration Results

| No. | Peak Name | Retention Time min | Area mAU*min | Height mAU | Relative Area % | Relative Height % | Amount n.a. |
|---|---|---|---|---|---|---|---|
| 1 | | 11,177 | 0,174 | 2,530 | 0,24 | 0,31 | n.a. |
| 2 | | 11,553 | 0,363 | 2,774 | 0,50 | 0,34 | n.a. |
| 3 | | 11,657 | 0,227 | 2,368 | 0,31 | 0,29 | n.a. |
| 4 | | 11,963 | 0,146 | 2,035 | 0,20 | 0,25 | n.a. |
| 5 | | 12,063 | 0,263 | 3,172 | 0,36 | 0,39 | n.a. |
| 6 | | 12,173 | 0,270 | 2,497 | 0,37 | 0,31 | n.a. |
| 7 | | 12,590 | 71,118 | 799,944 | 97,92 | 98,06 | n.a. |
| 8 | | 13,107 | 0,066 | 0,436 | 0,09 | 0,05 | n.a. |
| **Total:** | | | **72,627** | **815,755** | **100,00** | **100,00** | |

HPLC chromatogram of **20d**.

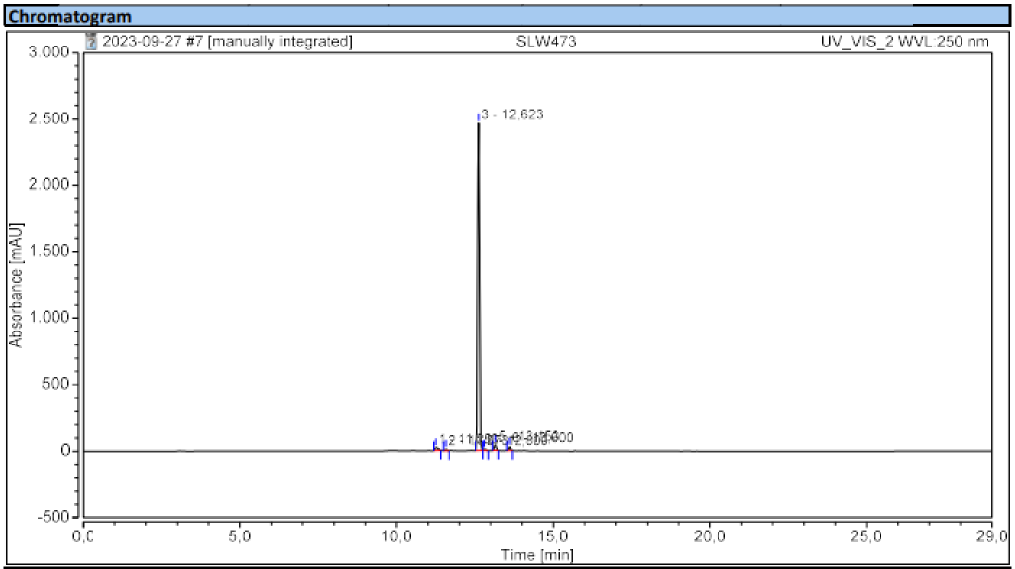


Integration Results

| No. | Peak Name | Retention Time min | Area mAU*min | Height mAU | Relative Area % | Relative Height % | Amount n.a. |
|---|---|---|---|---|---|---|---|
| 1 | | 11,253 | 2,777 | 23,428 | 1,53 | 0,91 | n.a. |
| 2 | | 11,573 | 0,743 | 10,712 | 0,41 | 0,41 | n.a. |
| 3 | | 12,623 | 172,521 | 2466,286 | 95,02 | 95,53 | n.a. |
| 4 | | 12,800 | 0,793 | 8,690 | 0,44 | 0,34 | n.a. |
| 5 | | 13,153 | 2,824 | 44,217 | 1,56 | 1,71 | n.a. |
| 6 | | 13,600 | 1,895 | 28,325 | 1,04 | 1,10 | n.a. |
| **Total:** | | | **181,554** | **2581,658** | **100,00** | **100,00** | |

HPLC chromatogram of **21a**.

**Chromatogram**

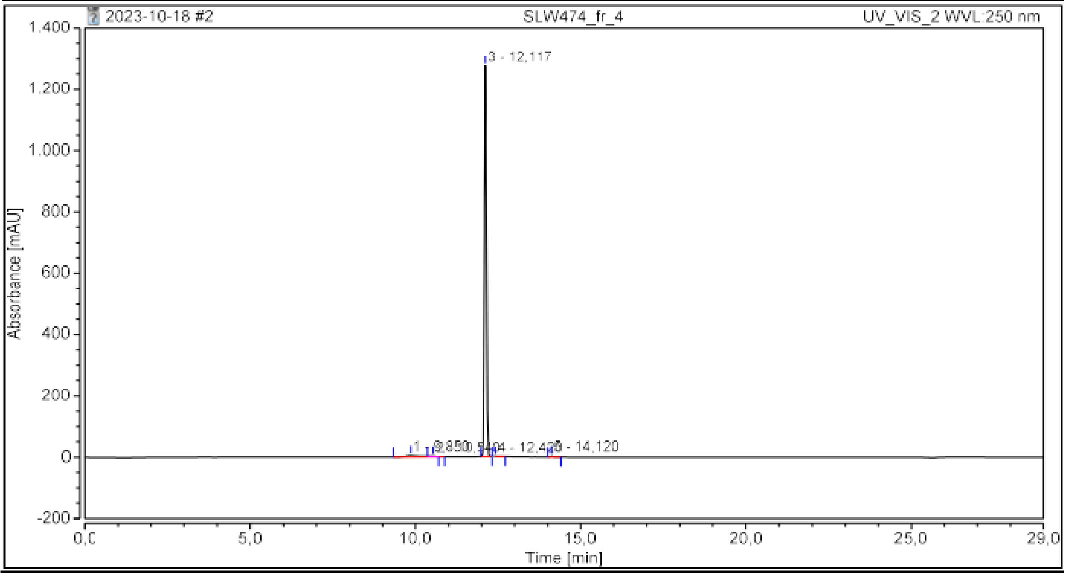


**Integration Results**

| No. | Peak Name | Retention Time min | Area mAU*min | Height mAU | Relative Area % | Relative Height % | Amount n.a. |
|---|---|---|---|---|---|---|---|
| 1 | | 9,850 | 2,472 | 4,404 | 2,65 | 0,34 | n.a. |
| 2 | | 10,540 | 0,127 | 0,831 | 0,14 | 0,06 | n.a. |
| 3 | | 12,117 | 90,371 | 1274,775 | 96,79 | 99,30 | n.a. |
| 4 | | 12,420 | 0,147 | 0,825 | 0,16 | 0,06 | n.a. |
| 5 | | 14,120 | 0,249 | 2,912 | 0,27 | 0,23 | n.a. |
| **Total:** | | | **93,367** | **1283,747** | **100,00** | **100,00** | |

HPLC chromatogram of **21b**.

**Chromatogram**

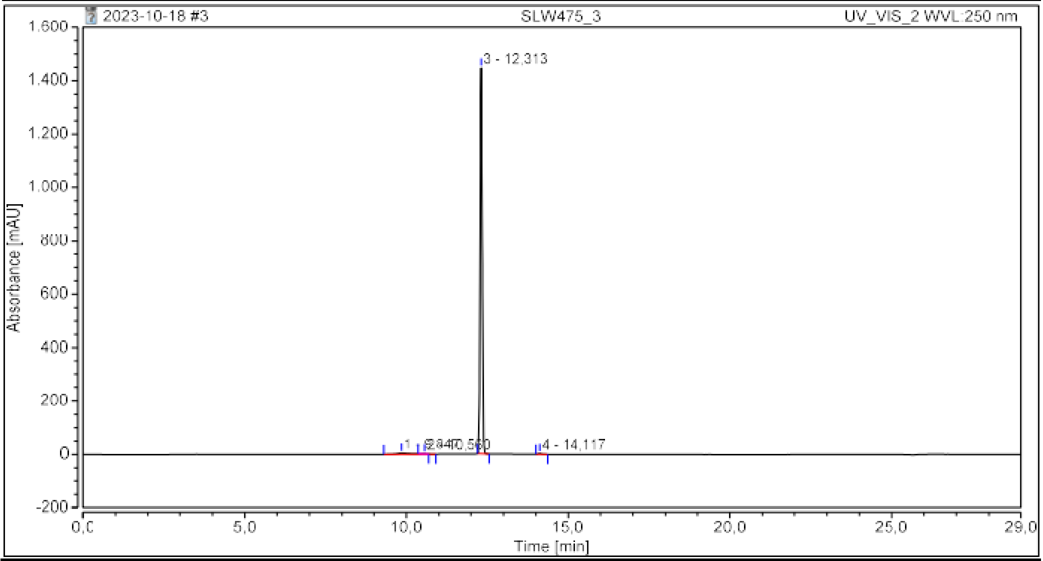


**Integration Results**

| No. | Peak Name | Retention Time min | Area mAU*min | Height mAU | Relative Area % | Relative Height % | Amount n.a. |
|---|---|---|---|---|---|---|---|
| 1 | | 9,847 | 2,464 | 4,405 | 2,32 | 0,30 | n.a. |
| 2 | | 10,560 | 0,147 | 0,915 | 0,14 | 0,06 | n.a. |
| 3 | | 12,313 | 103,387 | 1444,231 | 97,30 | 99,41 | n.a. |
| 4 | | 14,117 | 0,262 | 3,245 | 0,25 | 0,22 | n.a. |
| **Total:** | | | **106,260** | **1452,796** | **100,00** | **100,00** | |

HPLC chromatogram of **21c**.

**Chromatogram**

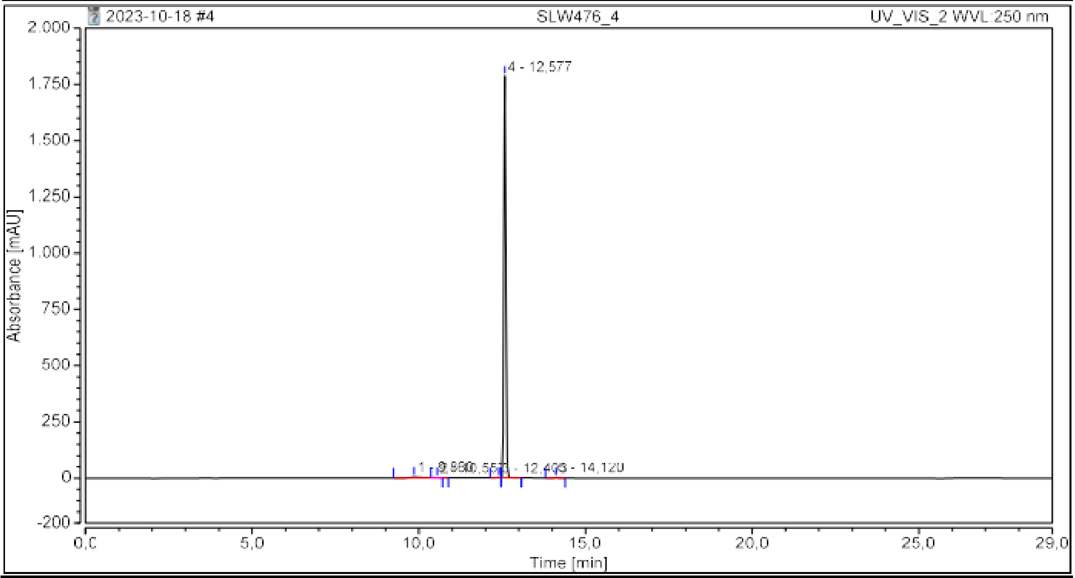


**Integration Results**

| No. | Peak Name | Retention Time min | Area mAU*min | Height mAU | Relative Area % | Relative Height % | Amount n.a. |
|---|---|---|---|---|---|---|---|
| 1 | | 9,860 | 2,497 | 4,405 | 1,89 | 0,25 | n.a. |
| 2 | | 10,557 | 0,132 | 0,826 | 0,10 | 0,05 | n.a. |
| 3 | | 12,403 | 0,196 | 1,881 | 0,15 | 0,10 | n.a. |
| 4 | | 12,577 | 129,293 | 1785,790 | 97,60 | 99,41 | n.a. |
| 5 | | 14,120 | 0,348 | 3,439 | 0,26 | 0,19 | n.a. |
| **Total:** | | | **132,466** | **1796,340** | **100,00** | **100,00** | |

HPLC chromatogram of **21d**.

**Chromatogram**

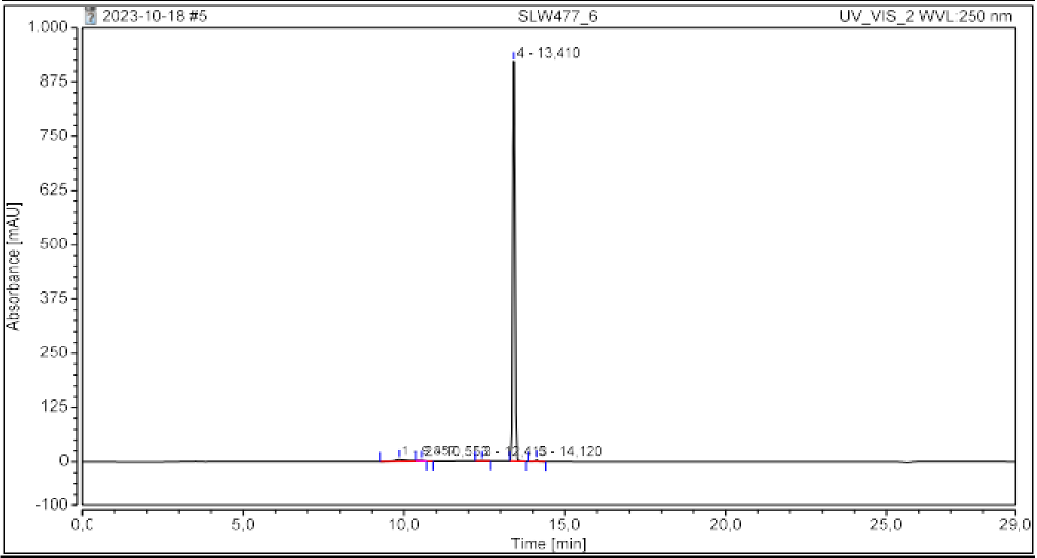


**Integration Results**

| No. | Peak Name | Retention Time min | Area mAU*min | Height mAU | Relative Area % | Relative Height % | Amount n.a. |
|---|---|---|---|---|---|---|---|
| 1 | | 9,857 | 2,494 | 4,483 | 3,35 | 0,48 | n.a. |
| 2 | | 10,553 | 0,141 | 0,914 | 0,19 | 0,10 | n.a. |
| 3 | | 12,413 | 0,100 | 0,565 | 0,13 | 0,06 | n.a. |
| 4 | | 13,410 | 71,404 | 919,660 | 95,88 | 98,99 | n.a. |
| 5 | | 14,120 | 0,334 | 3,415 | 0,45 | 0,37 | n.a. |
| **Total:** | | | **74,472** | **929,037** | **100,00** | **100,00** | |

HPLC chromatogram of **21e**.

**Chromatogram**

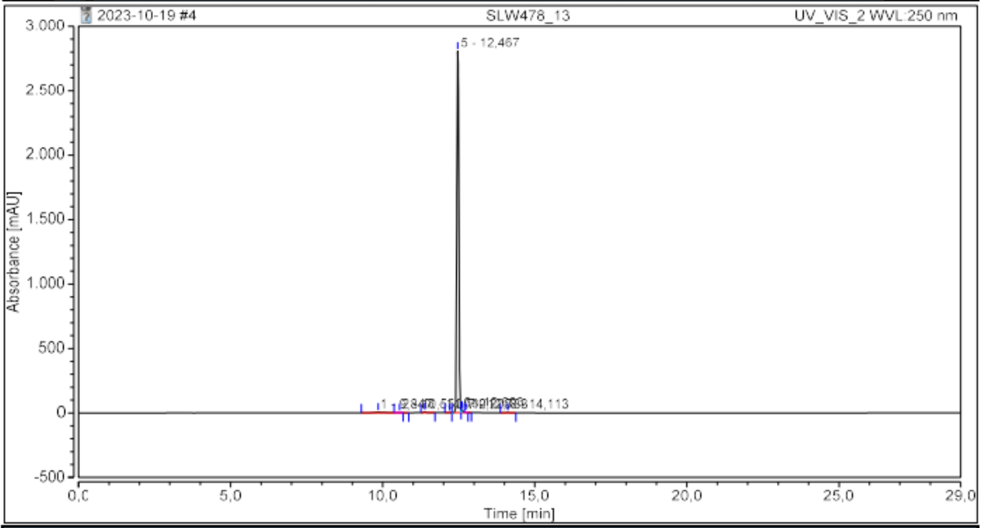


**Integration Results**

| No. | Peak Name | Retention Time<br>min | Area<br>mAU*min | Height<br>mAU | Relative Area<br>% | Relative Height<br>% | Amount<br>n.a. |
|---|---|---|---|---|---|---|---|
| 1 | | 9,847 | 2,447 | 4,516 | 1,15 | 0,16 | n.a. |
| 2 | | 10,550 | 0,109 | 0,746 | 0,05 | 0,03 | n.a. |
| 3 | | 11,380 | 0,808 | 7,159 | 0,38 | 0,25 | n.a. |
| 4 | | 12,200 | 0,110 | 1,136 | 0,05 | 0,04 | n.a. |
| 5 | | 12,467 | 206,955 | 2804,777 | 97,44 | 98,64 | n.a. |
| 6 | | 12,603 | 1,506 | 19,157 | 0,71 | 0,67 | n.a. |
| 7 | | 12,733 | 0,143 | 2,391 | 0,07 | 0,08 | n.a. |
| 8 | | 14,113 | 0,325 | 3,476 | 0,15 | 0,12 | n.a. |
| Total: | | | 212,402 | 2843,358 | 100,00 | 100,00 | |

HPLC chromatogram of **21f**.

**Chromatogram**

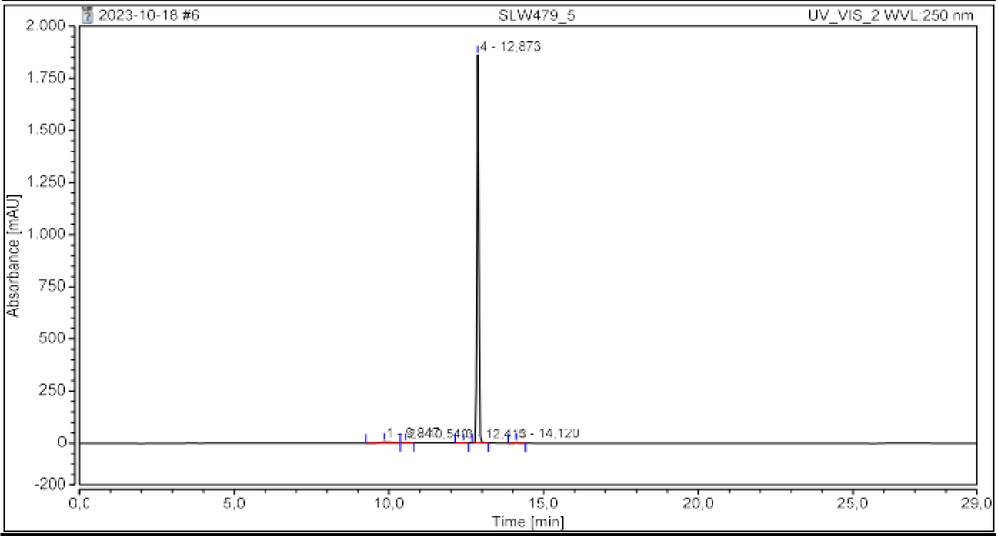


**Integration Results**

| No. | Peak Name | Retention Time<br>min | Area<br>mAU*min | Height<br>mAU | Relative Area<br>% | Relative Height<br>% | Amount<br>n.a. |
|---|---|---|---|---|---|---|---|
| 1 | | 9,847 | 2,084 | 4,270 | 1,59 | 0,23 | n.a. |
| 2 | | 10,540 | 0,443 | 1,781 | 0,34 | 0,10 | n.a. |
| 3 | | 12,413 | 0,074 | 0,494 | 0,06 | 0,03 | n.a. |
| 4 | | 12,873 | 128,498 | 1859,417 | 97,76 | 99,47 | n.a. |
| 5 | | 14,120 | 0,343 | 3,438 | 0,26 | 0,18 | n.a. |
| Total: | | | 131,442 | 1869,400 | 100,00 | 100,00 | |

HPLC chromatogram of **21g**.

**Chromatogram**

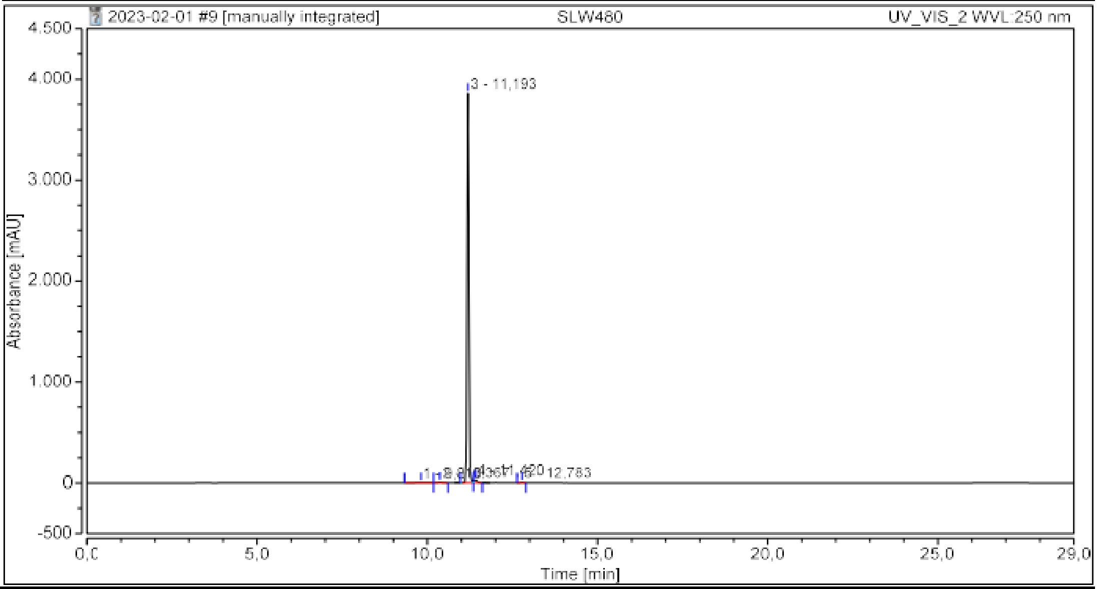


**Integration Results**

| No. | Peak Name | Retention Time<br>min | Area<br>mAU*min | Height<br>mAU | Relative Area<br>% | Relative Height<br>% | Amount<br>n.a. |
|---|---|---|---|---|---|---|---|
| 1 | | 9,810 | 1,562 | 3,797 | 0,57 | 0,10 | n.a. |
| 2 | | 10,367 | 0,994 | 6,179 | 0,36 | 0,16 | n.a. |
| 3 | | 11,193 | 269,811 | 3856,641 | 97,85 | 98,83 | n.a. |
| 4 | | 11,420 | 2,997 | 29,965 | 1,09 | 0,77 | n.a. |
| 5 | | 12,783 | 0,384 | 5,681 | 0,14 | 0,15 | n.a. |
| **Total:** | | | **275,748** | **3902,264** | **100,00** | **100,00** | |

HPLC chromatogram of **21h**.

**Chromatogram**

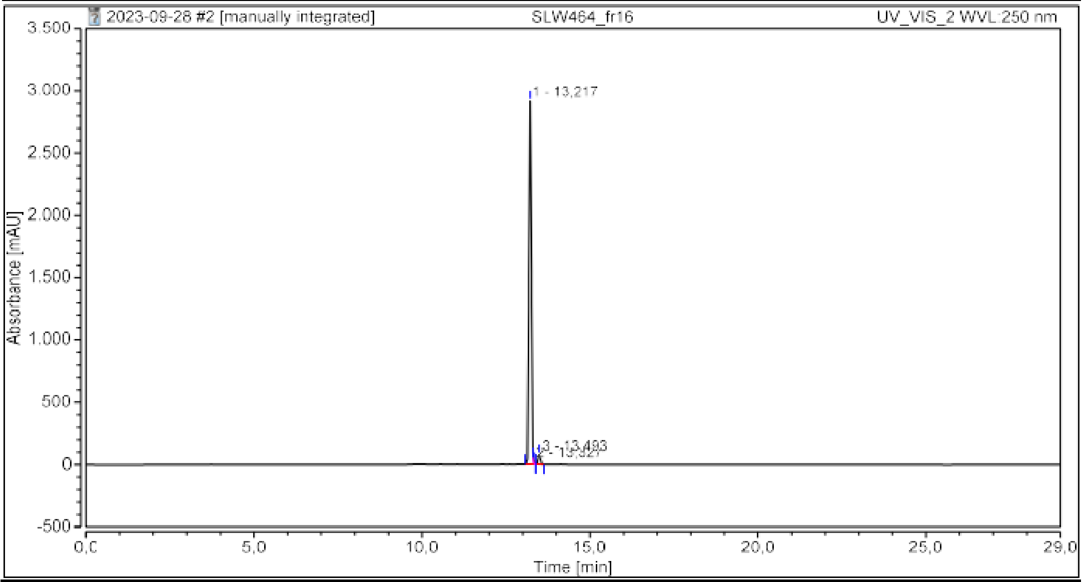


**Integration Results**

| No. | Peak Name | Retention Time<br>min | Area<br>mAU*min | Height<br>mAU | Relative Area<br>% | Relative Height<br>% | Amount<br>n.a. |
|---|---|---|---|---|---|---|---|
| 1 | | 13,217 | 268,050 | 2914,265 | 97,92 | 97,31 | n.a. |
| 2 | | 13,327 | 0,238 | 5,846 | 0,09 | 0,20 | n.a. |
| 3 | | 13,493 | 5,442 | 74,692 | 1,99 | 2,49 | n.a. |
| **Total:** | | | **273,730** | **2994,802** | **100,00** | **100,00** | |

HPLC chromatogram of **21i**.

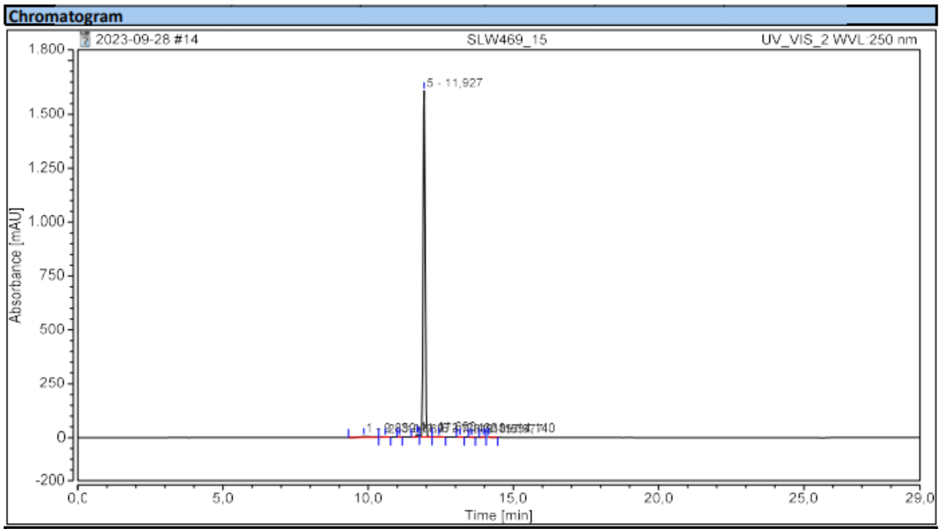


**Integration Results**

| No. | Peak Name | Retention Time min | Area mAU*min | Height mAU | Relative Area % | Relative Height % | Amount n.a. |
|---|---|---|---|---|---|---|---|
| 1 | | 9,850 | 1,680 | 3,750 | 1,24 | 0,23 | n.a. |
| 2 | | 10,587 | 0,387 | 1,551 | 0,29 | 0,09 | n.a. |
| 3 | | 11,073 | 0,140 | 1,641 | 0,10 | 0,10 | n.a. |
| 4 | | 11,690 | 1,092 | 12,272 | 0,81 | 0,75 | n.a. |
| 5 | | 11,927 | 130,701 | 1607,035 | 96,75 | 98,29 | n.a. |
| 6 | | 12,433 | 0,435 | 2,239 | 0,32 | 0,14 | n.a. |
| 7 | | 13,157 | 0,073 | 0,602 | 0,05 | 0,04 | n.a. |
| 8 | | 13,557 | 0,052 | 0,597 | 0,04 | 0,04 | n.a. |
| 9 | | 13,977 | 0,191 | 1,926 | 0,14 | 0,12 | n.a. |
| 10 | | 14,140 | 0,339 | 3,351 | 0,25 | 0,20 | n.a. |
| **Total:** | | | **135,090** | **1634,964** | **100,00** | **100,00** | |

HPLC chromatogram of **21j**.

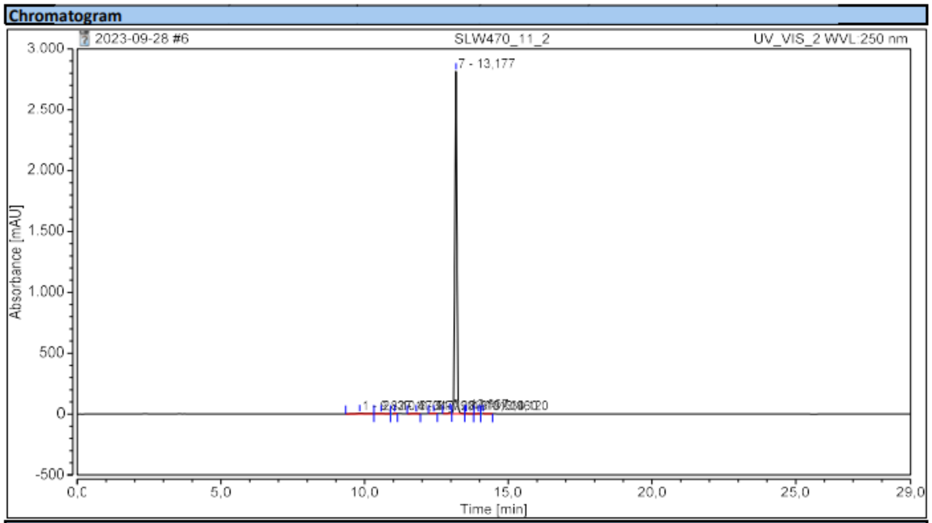


**Integration Results**

| No. | Peak Name | Retention Time min | Area mAU*min | Height mAU | Relative Area % | Relative Height % | Amount n.a. |
|---|---|---|---|---|---|---|---|
| 1 | | 9,827 | 1,738 | 3,821 | 0,70 | 0,13 | n.a. |
| 2 | | 10,573 | 0,535 | 1,821 | 0,22 | 0,06 | n.a. |
| 3 | | 11,047 | 0,115 | 1,309 | 0,05 | 0,05 | n.a. |
| 4 | | 11,790 | 0,195 | 1,189 | 0,08 | 0,04 | n.a. |
| 5 | | 12,407 | 0,187 | 1,404 | 0,08 | 0,05 | n.a. |
| 6 | | 12,967 | 1,035 | 15,222 | 0,42 | 0,54 | n.a. |
| 7 | | 13,177 | 243,591 | 2810,983 | 98,20 | 98,91 | n.a. |
| 8 | | 13,533 | 0,197 | 1,340 | 0,08 | 0,05 | n.a. |
| 9 | | 13,960 | 0,144 | 1,376 | 0,06 | 0,05 | n.a. |
| 10 | | 14,120 | 0,331 | 3,505 | 0,13 | 0,12 | n.a. |
| **Total:** | | | **248,067** | **2841,970** | **100,00** | **100,00** | |

HPLC chromatogram of **21k**.

**Chromatogram**

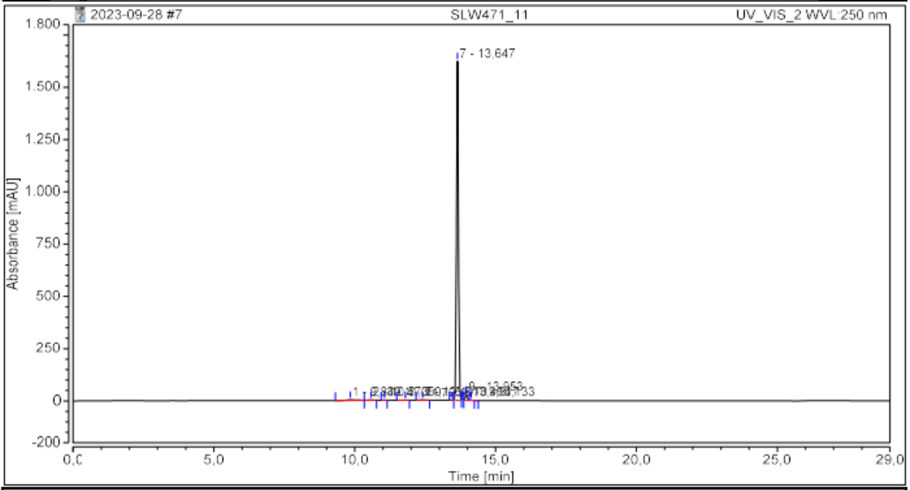


**Integration Results**

| No. | Peak Name | Retention Time min | Area mAU*min | Height mAU | Relative Area % | Relative Height % | Amount n.a. |
|---|---|---|---|---|---|---|---|
| 1 | | 9,840 | 1,710 | 3,849 | 1,26 | 0,23 | n.a. |
| 2 | | 10,573 | 0,421 | 1,609 | 0,31 | 0,10 | n.a. |
| 3 | | 11,050 | 0,129 | 1,414 | 0,10 | 0,08 | n.a. |
| 4 | | 11,793 | 0,186 | 1,159 | 0,14 | 0,07 | n.a. |
| 5 | | 12,407 | 0,293 | 1,751 | 0,22 | 0,10 | n.a. |
| 6 | | 13,460 | 0,117 | 1,486 | 0,09 | 0,09 | n.a. |
| 7 | | 13,647 | 129,715 | 1622,722 | 95,58 | 97,16 | n.a. |
| 8 | | 13,817 | 0,386 | 5,375 | 0,28 | 0,32 | n.a. |
| 9 | | 13,953 | 2,562 | 27,954 | 1,89 | 1,67 | n.a. |
| 10 | | 14,133 | 0,200 | 2,838 | 0,15 | 0,17 | n.a. |
| **Total:** | | | **135,720** | **1670,158** | **100,00** | **100,00** | |

HPLC chromatogram of **21l**.

**Chromatogram**

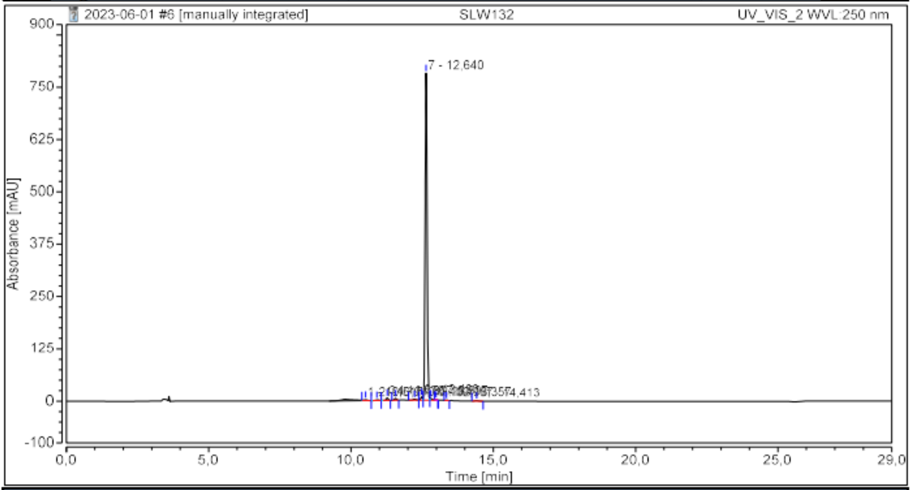


**Integration Results**

| No. | Peak Name | Retention Time min | Area mAU*min | Height mAU | Relative Area % | Relative Height % | Amount n.a. |
|---|---|---|---|---|---|---|---|
| 1 | | 10,517 | 0,329 | 2,068 | 0,51 | 0,25 | n.a. |
| 2 | | 10,910 | 0,181 | 1,448 | 0,28 | 0,18 | n.a. |
| 3 | | 11,273 | 0,473 | 6,134 | 0,73 | 0,75 | n.a. |
| 4 | | 11,567 | 0,394 | 4,868 | 0,61 | 0,60 | n.a. |
| 5 | | 12,233 | 0,343 | 2,671 | 0,53 | 0,33 | n.a. |
| 6 | | 12,480 | 0,626 | 8,056 | 0,97 | 0,99 | n.a. |
| 7 | | 12,640 | 61,420 | 781,478 | 95,15 | 96,02 | n.a. |
| 8 | | 12,807 | 0,461 | 4,197 | 0,71 | 0,52 | n.a. |
| 9 | | 12,987 | 0,053 | 0,784 | 0,08 | 0,10 | n.a. |
| 10 | | 13,357 | 0,065 | 0,693 | 0,10 | 0,09 | n.a. |
| 11 | | 14,413 | 0,208 | 1,476 | 0,32 | 0,18 | n.a. |
| **Total:** | | | **64,554** | **813,872** | **100,00** | **100,00** | |

HPLC chromatogram of SLW132 (**21m**).

(10) Chemical Computing Group ULC. *MOE*, 2022.

(19) Weier, A.-K.; Homrich, M.; Ebbinghaus, S.; Juda, P.; Miková, E.; Hauschild, R.; Zhang, L.; Quast, T.; Mass, E.; Schlitzer, A.; Kolanus, W.; Burgdorf, S.; Gruß, O. J.; Hons, M.; Wieser, S.; Kiermaier, E. Multiple centrosomes enhance migration and immune cell effector functions of mature dendritic cells. *J. Cell Biol.* **2022**, *221* (12). DOI: 10.1083/jcb.202107134.
(20) Waterborg, J. *The Lowry method for protein quantitation*:*The Protein Protocols Handbook*, 2009, 7-10.
(21) Gibson, D. G.; Young, L.; Chuang, R.-Y.; Venter, J. C.; Hutchison, C. A.; Smith, H. O. Enzymatic assembly of DNA molecules up to several hundred kilobases. *Nat. Methods* **2009**, *6* (5), 343–345. DOI: 10.1038/nmeth.1318.
(22) Masuho, I.; Ostrovskaya, O.; Kramer, G. M.; Jones, C. D.; Xie, K.; Martemyanov, K. A. Distinct profiles of functional discrimination among G proteins determine the actions of G protein-coupled receptors. *Sci. Signal.* **2015**, *8* (405), ra123. DOI: 10.1126/scisignal.aab4068.
(23) Chang, Y.; Bach, L.; Hasiuk, M.; Wen, L.; Elmzzahi, T.; Tsui, C.; Gutiérrez-Melo, N.; Steffen, T.; Utzschneider, D. T.; Raj, T.; Jost, P. J.; Heink, S.; Cheng, J.; Burton, O. T.; Zeiträg, J.; Alterauge, D.; Dahlström, F.; Becker, J.-C.; Kastl, M.; Symeonidis, K.; van Uelft, M.; Becker, M.; Reschke, S.; Krebs, S.; Blum, H.; Abdullah, Z.; Paeschke, K.; Ohnmacht, C.; Neumann, C.; Liston, A.; Meissner, F.; Korn, T.; Hasenauer, J.; Heissmeyer, V.; Beyer, M.; Kallies, A.; Jeker, L. T.; Baumjohann, D. TGF-β specifies TFH versus TH17 cell fates in murine CD4+ T cells through c-Maf. *Sci. Immunol.* **2024**, *9* (93), eadd4818. DOI: 10.1126/sciimmunol.add4818.
(24) Kopf, A.; Renkawitz, J.; Hauschild, R.; Girkontaite, I.; Tedford, K.; Merrin, J.; Thorn-Seshold, O.; Trauner, D.; Häcker, H.; Fischer, K.-D.; Kiermaier, E.; Sixt, M. Microtubules control cellular shape and coherence in amoeboid migrating cells. *J. Cell Biol.* **2020**, *219* (6). DOI: 10.1083/jcb.201907154.
